# How and why multicopy genes survive on the human Y chromosome

**DOI:** 10.1101/2024.04.02.587783

**Authors:** Karol Pal, Aleksandra Greshnova, Sukhwan Park, Byung June Ko, Hana Palova, Martin Steinegger, Sergei L. Kosakovsky Pond, Stefan Canzar, Kateryna D. Makova

**Affiliations:** Pennsylvania State University, University Park, PA, USA; University of Regensburg, Regensburg, Germany; Biozentrum, University of Basel, Basel, Switzerland; Seoul National University, Seoul, South Korea; Temple University, Philadelphia, PA, USA

## Abstract

The mammalian Y chromosome is an evolutionary paradox: it cannot recombine across most of its length and has lost many genes, yet it retains multicopy gene families required for sperm production. How these genes survive has been difficult to resolve because their near-identical copies collapse in most genome assemblies and are difficult to analyze accurately with short RNA reads. Using telomere-to-telomere assemblies of seven ape genomes together with long-read testis transcriptomes, protein-structure predictions, and selection tests, we show that these gene families vary extensively but encode structurally conserved proteins. We demonstrate that, while copy number varies across and within species, palindromes and tandem arrays are equally effective in homogenizing copies via gene conversion—a result we could obtain only with complete assemblies. We detect extensive transcript variation arising from alternative splicing and inter-copy sequence differences. Yet this variation collapses into only a few predicted protein-structure clusters per family. Furthermore, we find evidence of purifying selection acting on gene families (and domains) encoding ordered protein structures—the majority of multicopy families on the human Y. Thus, copy number and (to a lesser degree) sequence, and transcript structure and sequence all vary, whereas protein structure, which is important for spermatogenic function, is held nearly constant. Together, our results show that palindrome- and array-mediated homogenization and purifying selection aimed at preserving protein structure jointly enable the survival of fertility-related genes on the non-recombining Y chromosome. These results will inform diagnostic approaches to human Y-linked infertility and studies of the conservation of non-human great apes.

## Introduction

Sex chromosomes originated from autosomes independently in different taxa (Zhu, Younas, and Zhou 2025). The proto-Y chromosome in therian mammals acquired the male-determining *SRY* gene approximately 170 million years ago (Veyrunes et al. 2008). This event led to the suppression of recombination with the proto-X chromosome, followed by the accumulation of deleterious mutations and the degradation of the Y chromosome, including the loss of many genes (Skaletsky et al. 2003; Charlesworth 2003). The present-day human Y chromosome consists of two pseudoautosomal regions (PARs), which still recombine with the X chromosome, and X-transposed (acquired through a recent duplication from the X chromosome), X-degenerate (originating from proto-sex chromosomes), and ampliconic (having high sequence identity to itself) regions (Skaletsky et al. 2003). Ampliconic regions harbor multicopy genes (Y chromosome ampliconic genes, or YAGs) that are expressed in the testis and function in spermatogenesis (Skaletsky et al. 2003), making their preservation critical for reproduction. Highly repetitive ampliconic regions are thought to facilitate the maintenance of functional YAG copies by enabling intrachromosomal recombination (Ohta 1989; Charlesworth 2003; Connallon and Clark 2010; Marais, Campos, and Gordo 2010; Betrán, Demuth, and Williford 2012). Despite this theoretical knowledge, the precise evolutionary dynamics that maintain ampliconic gene integrity across species and how their structural organization affects gene expression and function remain incompletely understood.

Ampliconic regions appear to be a pervasive feature of heterogametic sex chromosomes and are likely critical for maintaining YAGs. These regions are usually organized into two major structural classes—palindromes or tandemly repeated arrays—which arose independently in multiple lineages. Palindromes are inverted repeats where two highly similar sequence arms flank a central spacer. The human Y harbors eight such palindromes (P1-P8) ranging from 30 kb to 2.9 Mb, containing most YAG families (Skaletsky et al. 2003; Trombetta and Cruciani 2017). Palindrome arms maintain >99.9% sequence identity via gene conversion, a type of intrachromosomal recombination (Rozen et al. 2003). Palindromic organization of heterogametic sex chromosomes is frequent across primates (Makova et al. 2024; Hughes, Skaletsky, and Page 2012); the great ape common ancestor already possessed sequences homologous to most human palindromes (Cechova et al. 2020). Palindromes have also evolved on heterogametic sex chromosomes in other species, e.g., in rabbit (Geraldes et al. 2010), some birds, such as sparrows and blackbirds (Davis et al. 2010), and some plants, such as willow (Zhou et al. 2020). In contrast, tandemly repeated arrays (later called “arrays”) are direct repeats of highly similar sequences. They characterize heterogametic sex chromosomes in, for example, mouse (Soh et al. 2014), felids (Brashear, Raudsepp, and Murphy 2018), cattle and sheep (Olagunju et al. 2024), chicken (Huang et al. 2023), and some plants, such as white campion (Hobza et al. 2006). Some species, such as apes, have both palindromes and arrays on their heterogametic sex chromosomes (Makova et al. 2024). However, which organization—in palindromes or arrays—is more efficient at facilitating gene conversion on the Y chromosome remains unexplored. Such information is critical for our understanding of YAG evolution across species and the mechanisms that maintain Y genes in humans.

Palindromes and arrays on the Y are prone to structural rearrangements due to non-allelic homologous recombination (Smith 1976), resulting in YAG copy number variation. Prior studies documented extensive YAG copy number variation across human Y haplogroups (Ye et al. 2018; Lucotte et al. 2018; Shi, Louzada, et al. 2019; Vegesna et al. 2019) and across species and individuals in great apes (Vegesna et al. 2020). Yet, how copy number variation translates into transcript expression and protein function has remained unclear due to technical challenges in resolving these repetitive regions.

Telomere-to-telomere (T2T) genome assemblies have transformed our ability to fully characterize the structure of Y chromosomes. The T2T assembly of the human Y, generated using long-read sequencing technologies, resolved its repetitive structure, including YAG family copies (Rhie et al. 2023). The T2T Y chromosome assemblies of six non-human ape species (bonobo, chimpanzee, gorilla, Sumatran orangutan, Bornean orangutan, and siamang) were generated with the same approach, and their YAG families have been completely deciphered (Makova et al. 2024). Population-level studies using T2T methodology analyzed Y chromosomes from 45 men, revealing substantial structural variation (Hallast et al. 2023). There is now a need to characterize the expression of YAG families while accounting for their organization on T2T-resolved Y chromosomes, as many of these genes are essential for reproduction.

Ampliconic genes encode proteins vital for spermatogenesis, making characterization of their expression patterns directly relevant to male fertility (Colaco and Modi 2018; Krausz and Abrardo 2025). Analysis of human YAG families using short-read RNA sequencing established that men with higher gene copy numbers generally exhibit increased expression levels, though substantial inter-individual variation exists (Vegesna et al. 2019). Comparative analysis across great apes revealed that, despite rapid evolution in copy number, expression levels remain conserved, suggesting dosage regulation (Vegesna et al. 2020). However, short-read approaches cannot reliably resolve transcript isoforms originating from gene copies with high sequence identity. Long-read transcriptomics overcomes this barrier by capturing full-length transcript sequences. The IsoCon algorithm was developed specifically to leverage Pacific Biosciences (PacBio) Iso-Seq data to reconstruct transcripts from multicopy gene families with high copy similarity (Sahlin et al. 2018). Application of this approach to great ape Y chromosomes revealed previously undetectable transcript isoform diversity, including species-specific splicing patterns (Tomaszkiewicz et al. 2023). However, this approach was not informed by the T2T assemblies that fully resolved YAG genomic copies, and thus, it was challenging to assign transcripts to individual gene copies. Therefore, systematic characterization of transcript isoforms across gene copies in ape species and understanding how transcript diversity translates into protein structure remained elusive.

Here, we present a comprehensive analysis of YAGs across great apes, integrating T2T genome assemblies, long-read testis transcriptomes, computational protein-structure predictions, and selection analyses. Building on the T2T Y-chromosome assemblies and coarse selection analysis of our previous study (Makova et al. 2024), we construct copy-resolved phylogenetic trees for each gene family, study the location of each copy in palindromes vs. arrays, produce the most complete repertoire of transcripts for these genes yet, link RNA transcripts to protein structures they produce, and explore selection on gene copies and their structural domains. We find that these genes tolerate extensive variation in copy number, transcript identity, and sequence, yet their sequences are homogenized by gene conversion, and their ordered protein structures are conserved across copies and species by purifying selection. The results of this study provide a possible explanation for the paradox behind the persistence of multicopy genes on the Y chromosome and are relevant to conservation strategies for non-human great apes (all of which are endangered) and human infertility studies.

## Results

### Gene conversion homogenizes Y ampliconic gene copies equally well in palindromes and tandem arrays, despite extensive copy-number variation

The availability of T2T genomes for the Y chromosome in human and other great apes (Rhie et al. 2023; Makova et al. 2024) provides an opportunity to evaluate the evolutionary history and copy number variation of human YAGs. Below we analyzed the seven multicopy families present on the human Y (Skaletsky et al. 2003)—*BPY2, CDY, DAZ, HSFY, RBMY, TSPY,* and *VCY*—in the T2T genomes of human and its closest relatives (bonobo, chimpanzee, gorilla, Bornean orangutan, Sumatran orangutan, and siamang (Yoo et al. 2025)). The *XKRY* and *PRY* gene families were analyzed in previous studies (Skaletsky et al. 2003; Bhowmick, Satta, and Takahata 2007; Lahn and Page 1997; Cechova et al. 2020; Katrien Stouffs et al. 2004; K. Stouffs et al. 2001; Tomaszkiewicz et al. 2023), but not here because they are annotated as “pseudogene” or “non-coding RNA”, respectively, in the latest human and ape T2T assemblies (Table S1). We also searched for non-Y homologs of the seven gene families of interest (Table S2), confirming the transposition origin of *CDY* and *DAZ* and the proto-sex-chromosome origin of *RBMY* and *VCY* (Note S1).

We next studied variation in copy number across the seven YAG families, based on our manual curation of YAG annotations (Methods; Table S3) and, when available, across multiple individuals. For humans, copy numbers were inferred (see Methods) from 45 T2T or nearly T2T Y chromosome assemblies (Hallast et al. 2023). For chimpanzee and gorilla, copy numbers were estimated using `AMPLICONE` (Vegesna et al. 2019) from 24 and 18 individuals, respectively, for which resequencing data are available (see Methods). For bonobo, Bornean orangutan, Sumatran orangutan, and siamang, we counted the number of copies in their reference assemblies (Makova et al. 2024), as the amount and quality of additional sequencing data for these species were limited. Overall, we found (**Figure 1**) that four gene families (*BPY2*, *DAZ*, *HSFY*, and *VCY*) had consistently low copy numbers (≤10 copies). The remaining three gene families (*CDY*, *RBMY*, and *TSPY*) had a wide range of copy numbers. Variability in copy number for chimpanzees, gorillas, and humans was proportional to the average copy number for the gene family in each species (Figure S1). Using `CAFE` (Mendes et al. 2021) on median copy counts across individuals (which reduces outlier effects—e.g., reference HG002 has 45 *TSPY* copies versus a median of 32 across 45 men), we detected significant lineage-specific expansions and contractions in *CDY*, *DAZ*, *RBMY*, and *TSPY* (Table S4A), confirming substantial copy-number variation both within and across species.

**Figure 1.**
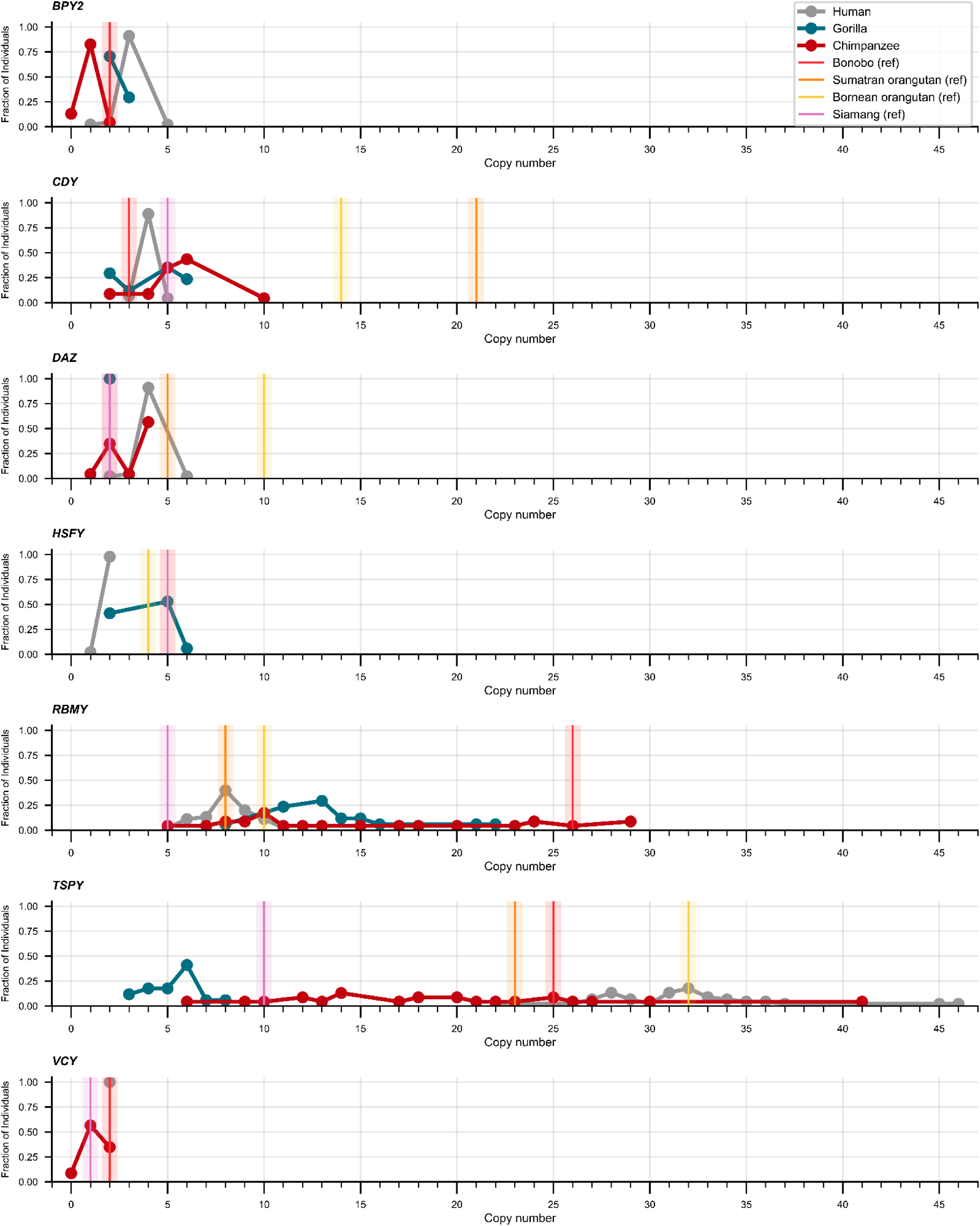
Inter- and intraspecies variation in copy number of Y ampliconic genes. Copy number is plotted on the X axis, and the proportion of individuals on the Y axis. A vertical line was used when only one individual per species was available.

Many human and other ape YAGs are located within palindromes, yet some can be organized into tandemly repeated arrays, hereafter called arrays (Makova et al. 2024; Rhie et al. 2023) the full extent of palindromes and arrays has only been revealed with the assembly of T2T genomes. Each of these two organizations facilitates gene conversion (Lange et al. 2009) (**Figure 2A**), but their effectiveness has not been directly compared before. We hypothesized that being positioned on the opposite arms of the same palindrome provides particularly favorable conditions for homogenization via gene conversion in YAGs (Betrán, Demuth, and Williford 2012; Rozen et al. 2003).

**Figure 2.**
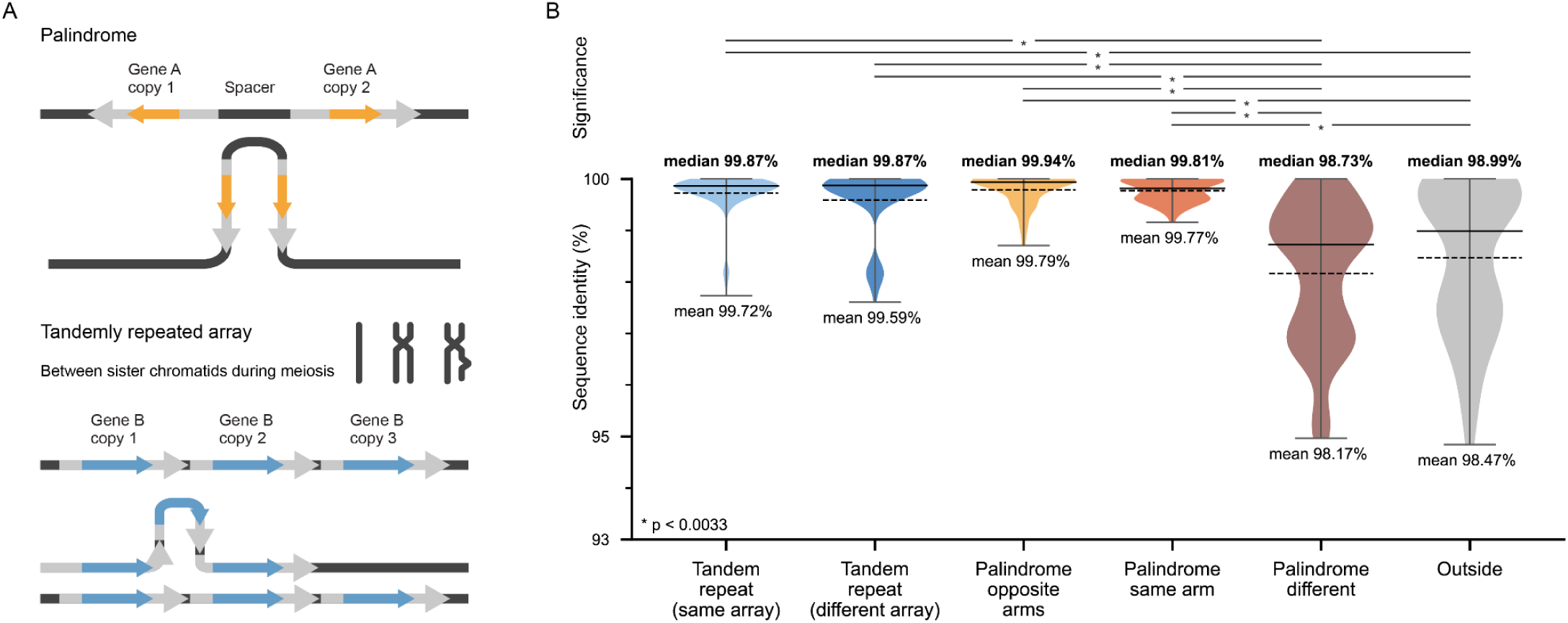
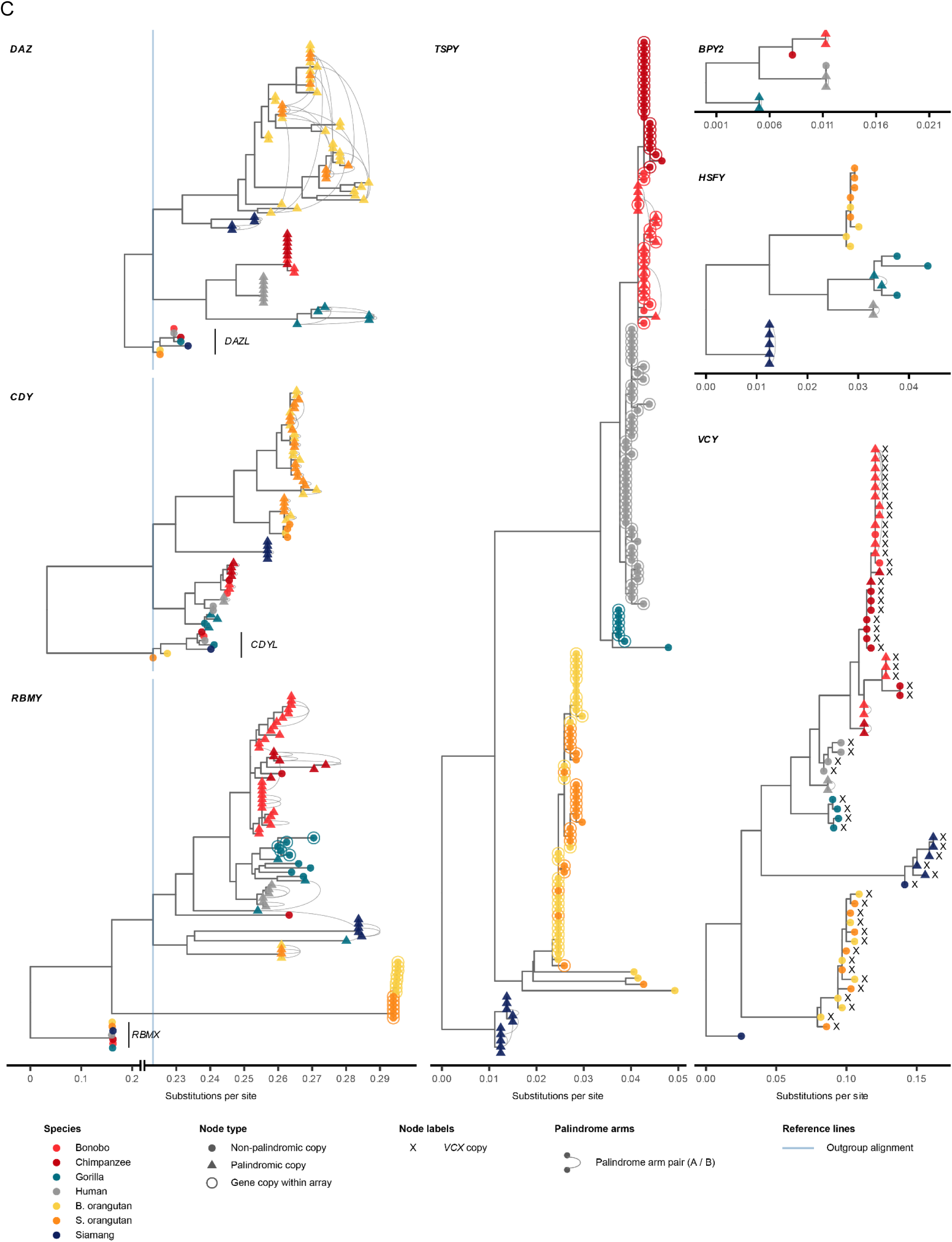
Chromosomal organization of YAGs. (**A**) Two models of homogenization of ampliconic genes via intrachromosomal recombination between palindrome arms and between direct repeats in tandemly repeated arrays. (**B**) Violin plots showing the distribution of pairwise sequence identities and pairwise permutation test results comparing sequence identity for protein-coding sequences of gene copies from the same gene family and from the same species. The analysis was performed across gene copies located within tandem arrays (1,391 comparisons across 136 gene copies), from different arrays (297 comparisons across 69 gene copies), located on opposite arms of the same palindromes (44 comparisons across 80 gene copies; gene copies were compared only if they were in the opposite orientation), on the same arm of a palindrome (10 comparisons across 20 gene copies), within different palindromes (446 comparisons across 78 gene copies), and outside of palindromes and arrays (89 comparisons across 55 genes). For underlying data, see Table S19. (**C**) Phylogenetic trees of seven YAG gene families: *DAZ* RRMs, *CDY*, and *RBMY* (left column, shared interrupted scale); *TSPY* (center column); and *BPY2*, *HSFY*, and *VCY* (right column, shared scale). Trees of *DAZ* RRM and *CDY* were rooted with the closest autosomal paralogue (*DAZL* and *CDYL,* respectively). The *RBMY* tree was rooted with the X-linked paralogue (*RBMX*). All other trees were rooted with Y-chromosome homologs from the most distantly related species (gorilla in *BPY*, siamang in *HSFY*, *TSPY*, and *VCY*). The color of external nodes indicates species. Circles denote non-palindromic gene copies; triangles denote copies residing within a palindrome. A ring surrounding a node indicates that the gene copy is located within a tandemly repeated array. Curved arches connect the two arms (A and B) of the same palindrome within a species. In the left column, the scale is compressed below 0.22 and above 0.29 substitutions per site to accommodate the wide range of branch lengths while preserving resolution in the intermediate range; gray vertical lines mark 0.1 substitutions/site intervals. Trees in the left column are horizontally aligned at the positions of their respective outgroup nodes (blue vertical lines) to facilitate comparison of within-family diversification.

To test this hypothesis, all YAG copies in reference ape genomes were classified as located within palindromes, within arrays, within both palindromes and arrays, or outside of arrays and palindromes (Table S5A). Palindromes were defined with `PALINDROVER` (Makova et al. 2024). Arrays were identified using `AMPLICOVER`, which detects sequence arrays based on genomic spacing between adjacent identical *k*-mers (see Methods). *BPY2*, *CDY*, *DAZ*, *HSFY*, and *VCY* were usually organized into palindromes, *TSPY* into arrays, and *RBMY* into both palindromes and arrays (Table S5A). Whereas human possessed one *TSPY* array, bonobo and orangutans had multiple *TSPY* arrays (three in bonobo, two in Sumatran orangutan, and three in Bornean orangutan; Table S5A). In total, across the seven reference ape Y chromosomes, we detected 26 gene-carrying palindromes, each harboring, on average, 6.3 YAG copies, and 14 gene-carrying arrays, each carrying, on average, 10 YAG copies (Table S5C). Altogether, 163 and 140 YAG copies were located within palindromes and arrays, respectively (not counting 11 gene copies present both within an array and a palindrome), and 46 YAG copies were present in neither palindromes nor arrays. Thus, most YAG copies were located within palindromes or arrays. Moreover, both palindromes and arrays held similar amounts of gene copies. However, palindromes were more numerous, and thus each palindrome, on average, carried fewer genes than an array.

To evaluate whether palindromes or arrays lead to greater sequence homogenization, we examined pairwise sequence identity among gene copies (from the same gene family) within them (**Figure 2B**, Table S6; Methods). The median pairwise nucleotide identity was highest between gene copies on opposite arms of the same palindrome (99.94%), closely followed by that for copies from the same array (99.87%), with no statistically significant difference between these two values (*p* = 0.2608; two-sided permutation test with 10,000 permutations). In fact, there was no statistically significant difference in median pairwise identity between any of the four categories; genes on the same array, genes on different arrays (99.87%), gene copies located on the opposite arms of the a palindrome, and genes located on the same arm of the same palindrome (99.81%). Being located on different palindromes or outside of arrays/palindromes led to a significantly lower pairwise sequence identity (98.73% and 98.99%, respectively) than being located within any of the four previous categories (**Figure 2B**, Table S6). We observed a statistically significant decrease in identity with increasing distance between copies located in the same array (Figure S2). In conclusion, both palindromic organization and tandem repetition are equally effective at homogenizing YAG sequences.

Maximum-likelihood phylogenies of paralogs and orthologs within each family (**Figure 2C**; Note S2) corroborated gene-conversion–driven homogenization. Consistent with the architecture analysis above (**Figure 2B**), copies on opposite arms of the same palindrome formed exclusive monophyletic clades in 22 of 26 cases, as did copies within arrays (**Figure 2C**)—confirming that both architectures drive homogenization. *DAZ* was a notable exception to uniform homogenization, with its RNA-recognition motifs (RRMs) clustering tightly by genus or species while its 72/90-bp exon repeats evolved faster and showed weaker phylogenetic structure (Figure S3). Phylogenies also revealed that sequence homogenization scales with divergence time (Makova et al. 2024): copies remained intermingled between the two *Pongo* species (∼1 MYA divergence) and the two *Pan* species (∼2.5 MYA divergence but resolved into genus-specific monophyletic clades by the *Pan*–human split (∼7 MYA divergence), bracketing the time required to homogenize YAG copies. Together, these results highlight palindromes and tandem arrays as powerful and equivalent drivers of Y-chromosome gene homogenization, modulated by lineage-specific evolutionary histories.

### Extensive transcript variation arises from gene duplication and alternative splicing

After establishing that palindromes and arrays are equally important in sequence homogenization, we next asked how variability in copy number and sequence differences across gene copies manifests in transcript variation. This question could not be addressed prior to the availability of T2T assemblies and long-read YAG transcriptomes. To answer it, we reconstructed transcript isoforms from testis tissue (where these genes are expressed) of the six great ape species analyzed in this study (no testis samples were available for the siamang). We generated long transcript reads with PacBio Iso-Seq and short transcript reads with Illumina RNA-seq (see Methods), and combined them to assemble YAG transcriptomes with `STRINGTIE` (Shumate et al. 2022), using great ape T2T genomes as references (Makova et al. 2024). We performed variant calling on the PacBio reads to assess sequence variation among transcripts originating from different YAG copies (see Methods for details).

### Mechanisms of transcript diversity

We found that transcript diversity results from alternative splicing (i.e., generating *structural isoform variants*) and sequence variation across gene copies (i.e., generating *sequence isoform variants*), and studied how the relative contributions of these two mechanisms differ across gene families and species. The *DAZ* gene family required a separate investigation and is discussed last.

Structural isoform diversity scaled with gene structure: we found a significant positive correlation between the number of exons and structural isoform variant count for YAGs (r = 0.610, *p* = 9.35×10^-4^, Figure S4A). Three gene families (*BPY2*, *RBMY*, and *TSPY*) had abundant structural isoform variants (6, 11, and 7, respectively), whereas three gene families with fewer exons (*CDY*, *HSFY*, and *VCY*) had only two (**Figure 3A**).

**Figure 3.**
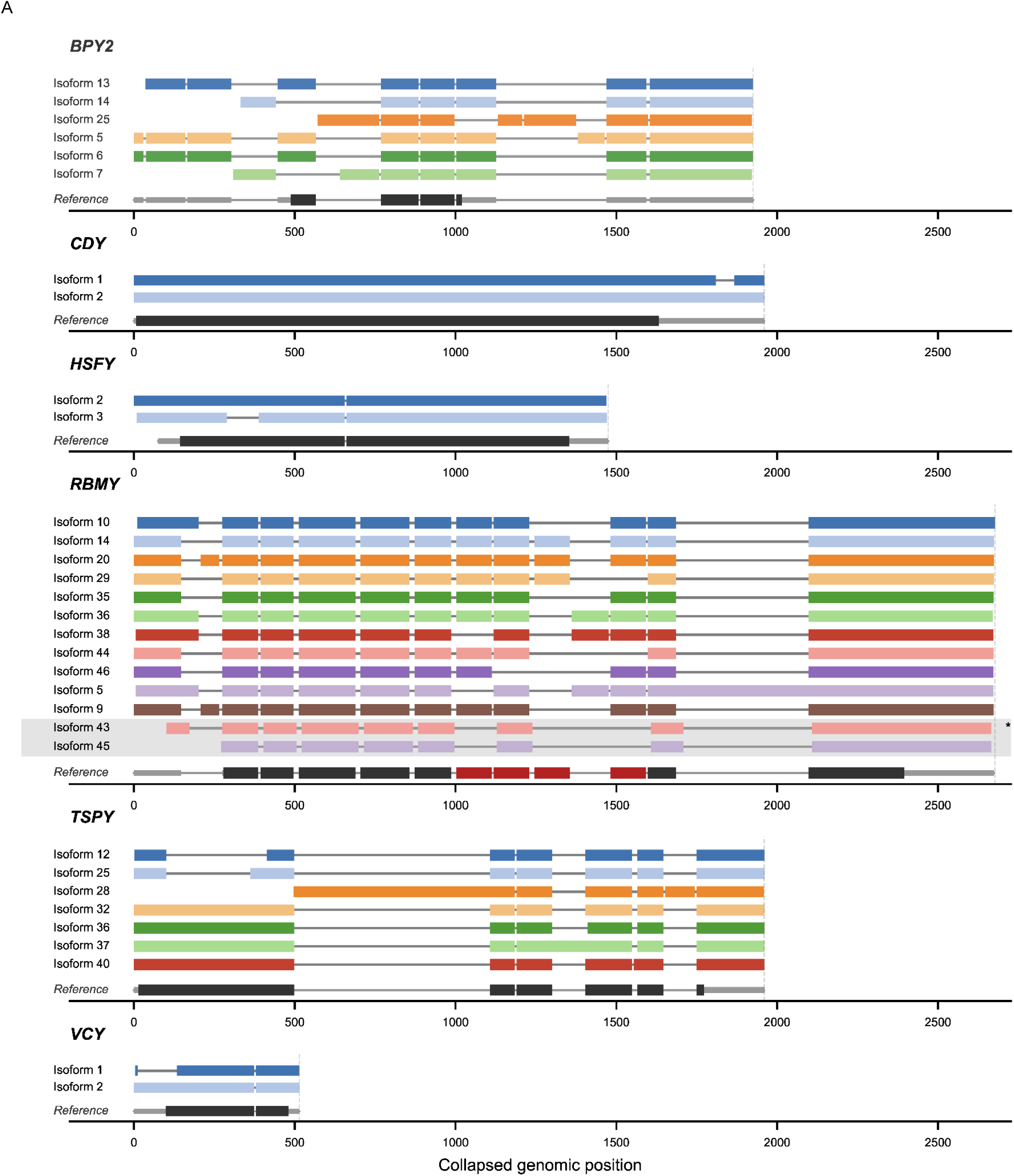
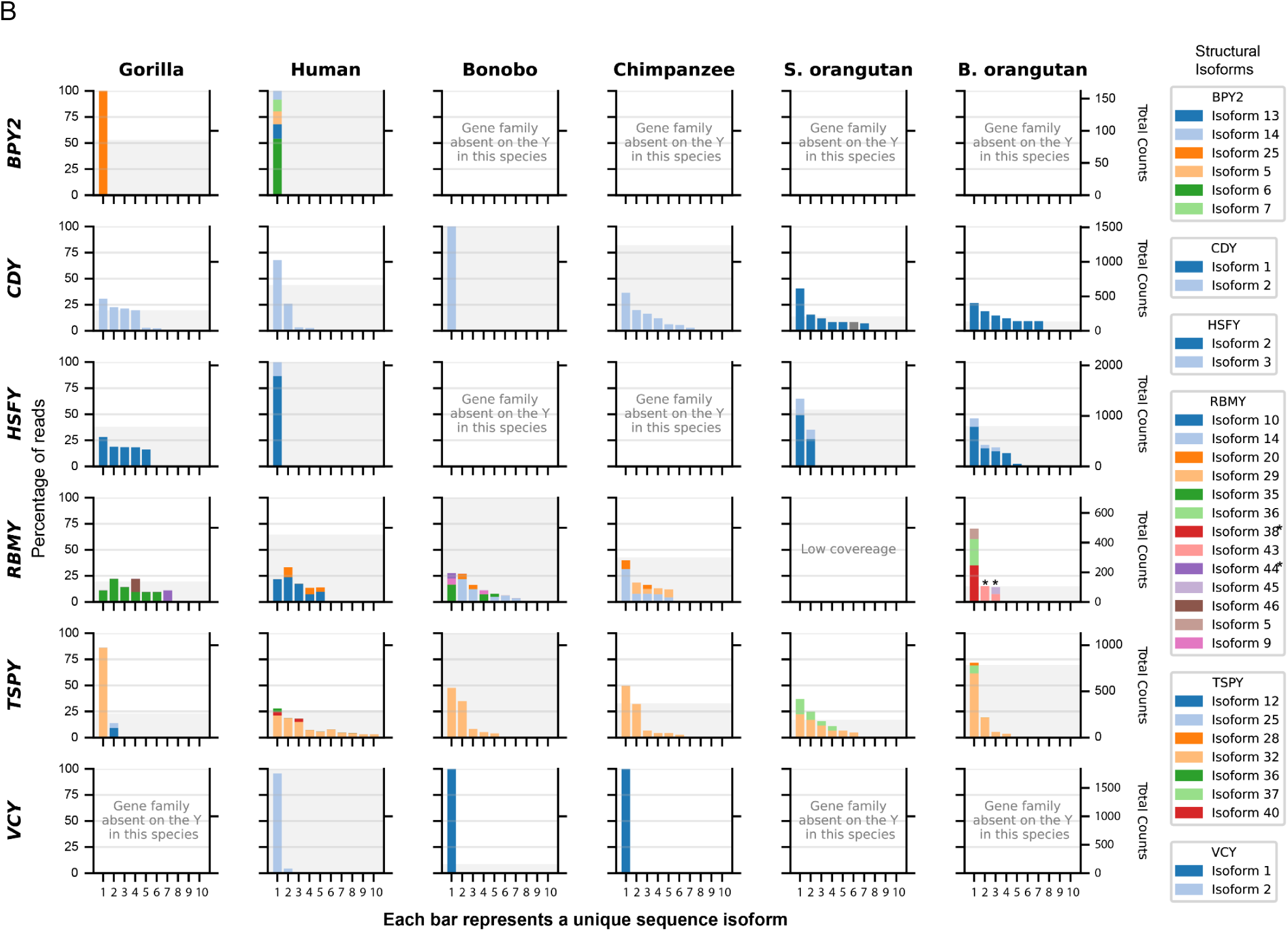
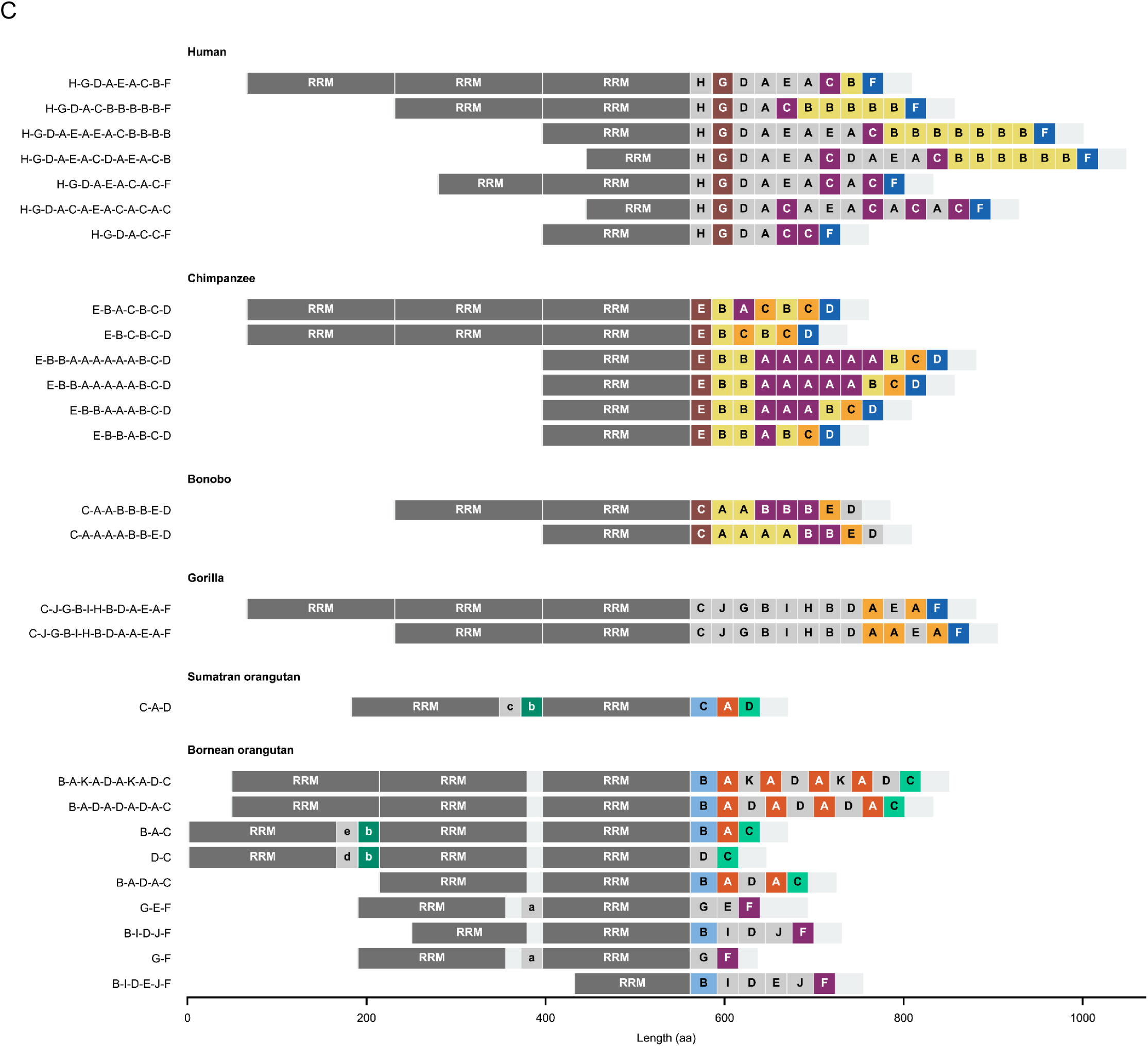
Distribution of structural and sequence RNA isoforms of YAGs. (**A**) Structural transcript isoforms for six gene families (*BPY2*, *CDY*, *HSFY*, *RBMY*, *TSPY*, and *VCY*). Each isoform was aligned to a single human reference copy. Colored rectangles denote RNA sequence aligned to exons, and introns are shown as gray lines. Intronic sequences without aligned RNA sequences are collapsed. Each color represents a distinct structural isoform for each YAG family. Human reference gene copies are shown under the colored isoforms: thicker black boxes represent the coding sequence, whereas thinner gray blocks represent UTRs. The burgundy color represents *RBMY*’s SRGY-box containing 111-bp exon repeat units (Ma et al. 1993; Elliott 2004). (**B**) Each panel displays the relative *sequence isoform* usage (individual bars) for a given gene family (rows) and species (columns). Colors distinguish individual *structural isoforms* within each gene family, as indicated by legends on the right (color scheme matching part A). Stacked bars represent the proportion of reads supporting each structural isoform for a given sequence isoform, normalized to the total number of classified reads at that panel (left y-axis, 0–100%). The light gray shading on each panel indicates the total read depth for a YAG family in a sample from a given species (right y-axis, total counts). (**C**) *DAZ* transcript isoforms. Each horizontal bar represents a unique *DAZ* transcript isoform, aligned at the rightmost RRM domain. Dark gray blocks indicate RRM domains; colored blocks indicate *DAZ* repeat units, with each color representing a distinct repeat sequence variant shared across two or more species. Gray repeat blocks are species-specific variants. Repeat units embedded between RRM domains are also shown and labeled. Uppercase letters identify repeat variants occurring after the RRM motifs; lowercase letters denote inter-RRM repeat variants. Sumatran orangutan *DAZ* expression levels were too low to draw a conclusion. “*” indicates Pongo-specific *RBMY* isoforms.

Sequence isoform diversity scaled with copy number: we found a significant positive correlation between copy number count and the number of sequence isoform variants (r = 0.688, *p* = 1.02×10^-4^, Figure S4B). This relationship could be quantified only once T2T assemblies resolved the individual gene copies. High-copy families (*TSPY*, *RBMY*, and *CDY*) showed extensive isoform diversity (in most species). In contrast, low-copy families *BPY2* and *VCY* showed low to no isoform diversity (across all species, **Figure 3B**). A similar pattern was observed for bonobo *CDY*, human *HSFY*, and gorilla *TSPY* (**Figure 3B**)—all of which also have low copy numbers (**Figure 1**).

### Transcript isoforms for each gene family

We next combined information about structural and sequence isoform variants to detect distinct transcript isoforms and studied their distribution across species (Additional Data File 1; **Figure 3B**, bars represent unique gene copy sequences, while their color represents distinct structural isoforms). Among families consistently expressed across species, *TSPY* and *RBMY* showed the greatest diversity. *TSPY* structural isoform variation was driven mainly by intron retention, with all species favoring structural isoform 32 expressed by multiple sequence isoforms. *RBMY* transcript diversity was concentrated in a 111-bp repeat region and, to a lesser extent, alternative splicing of the 5′ untranslated region (UTR), with some species-specific isoforms present (**Figure 3B**), e.g., orangutans expressed unique isoforms in which a ∼550-bp deletion removes one 111-bp repeat exon while including a different repeat skipped in other species (Figure S5). For *CDY* and *HSFY*, diversity was dominated by sequence rather than structural variation; orangutans additionally exhibited a divergent *CDY* structural isoform with an earlier initiation codon (Figure S6). *BPY2* and *VCY* were each expressed in only a few species, with limited isoform diversity consistent with their low copy number.

*DAZ* isoforms in all species (**Figure 3C**) carried up to three RRMs and a tail of tandemly repeated *DAZ* repeats of variable count (from two in Bornean orangutan to 19 in human), which we studied with a separate approach (see Methods for details; Additional Data File 2). In the *Pongo* lineage, *DAZ* repeats were sometimes also interspersed between RRMs. In human, we recovered isoforms comprising all previously described *DAZ* repeat sequences in (Fu et al. 2015), validating our approach (Additional Data File 2).

In summary, using long-read sequencing, we reconstructed full-length transcripts of great ape YAG families—one of the most complicated repetitive regions of the genome—and have demonstrated that its complexity arises from both gene duplication and alternative splicing. We have also established that the use of an improved, long-read reference transcriptome can affect RNA-seq expression estimates (Note S3, Figure S7, Table S7).

### High transcript diversity translates into a limited number of protein clusters

We next asked whether this transcript diversity extends to protein structure, where the spermatogenic functions of these genes are specified. Overall, based on sequence, and as expected due to the presence of synonymous variants, there were fewer predicted protein isoforms (later called ‘protein isoforms’) than observed transcript isoforms (**Figure 4A**). Using `COLABFOLD` (Mirdita et al. 2022), we predicted protein structures of all unique protein isoforms. Among seven YAG families analyzed, five (*CDY*, *DAZ*, *HSFY*, *RBMY*, and *TSPY*) had high-confidence scores of structure prediction at least over portions of their protein sequences (Figure S8). Two (*BPY2* and *VCY*, Figure S9A-B) had low-confidence scores (Figure S8), suggesting intrinsic disorder (*VCY*) or low confidence in the predicted structure (*BPY2*). We next clustered the predicted protein isoform structures with `FOLDSEEK` (van Kempen et al. 2024), which assigns similar structures to a single representative protein structure via structural alignment (Table S8; **Figure 4A**).

**Figure 4.**
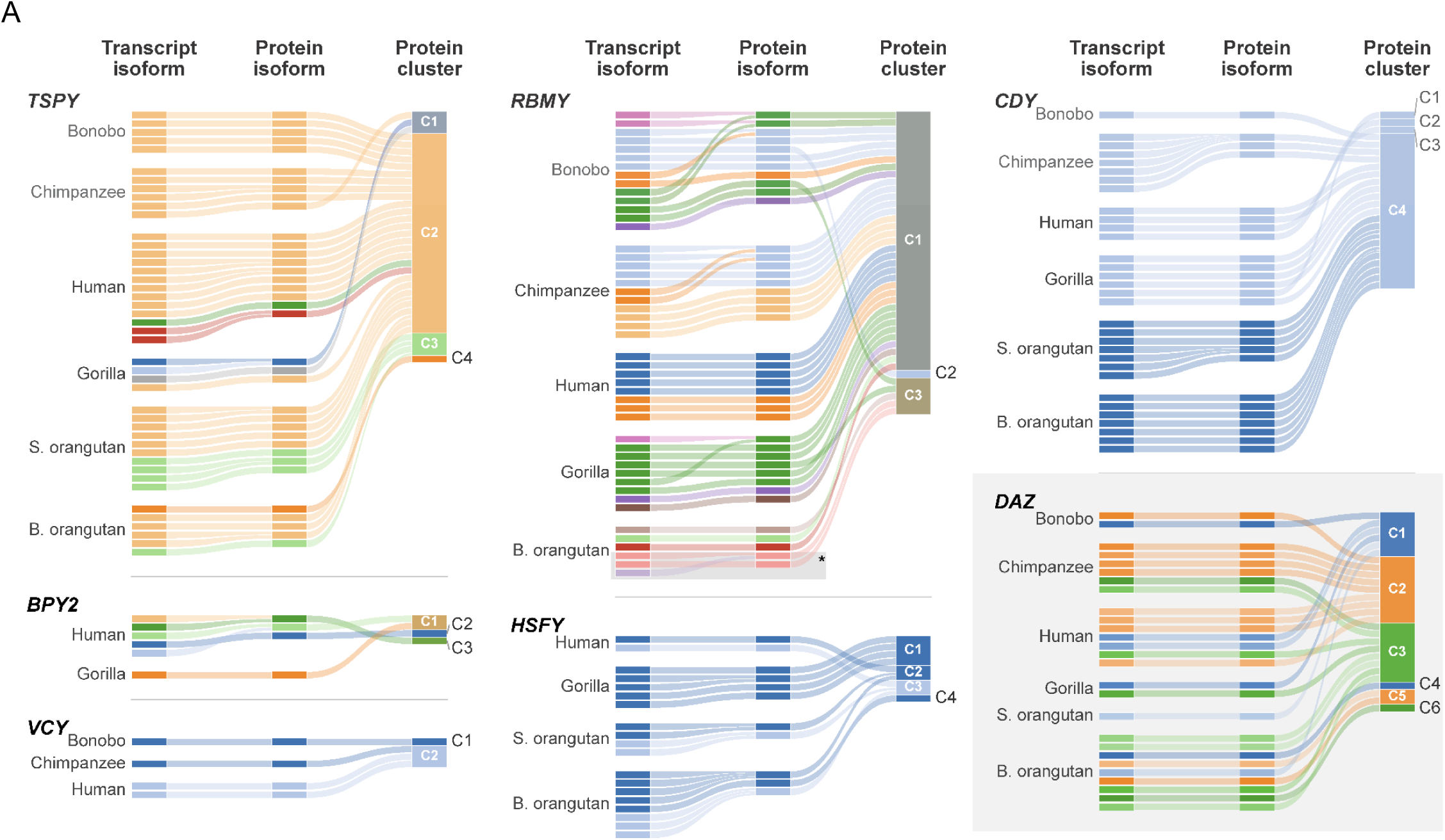
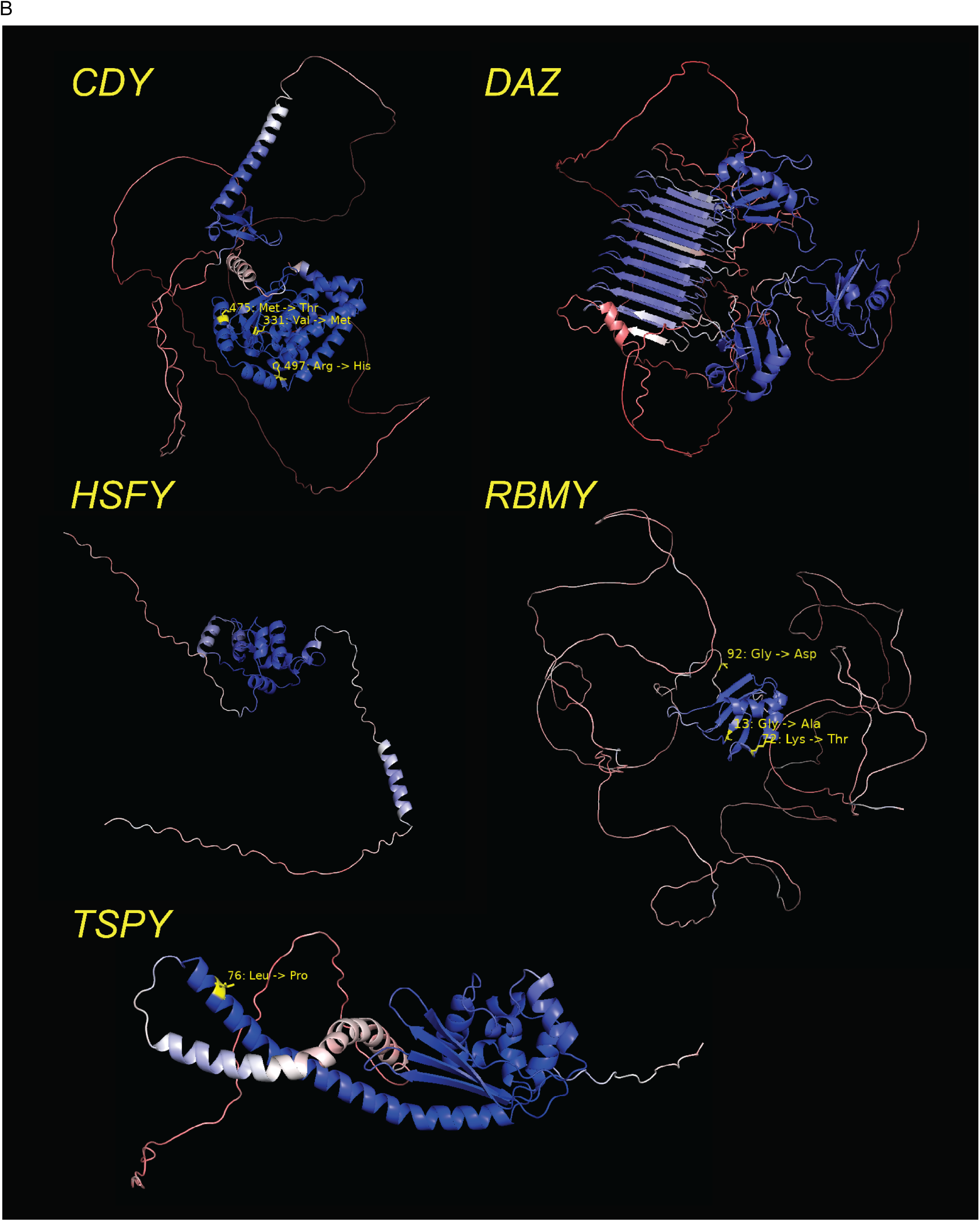
Protein isoforms and confidence levels of structures predicted with ColabFold. (**A**) Relationships between transcript isoforms, protein isoforms, and structure-derived protein clusters. Each subplot—one for each gene family—comprises three columns connected with ribbons. Column one consists of bars representing unique transcript isoforms. For *BPY2, CDY*, *HSFY*, *RBMY*, *TSPY*, and *VCY*, the color of the bars is derived from the structural isoform the transcript represents based on panel A. Multiple bars of the same color represent multiple sequence isoforms of the same structure. Bars in the second column represent protein isoforms, and color ribbons between columns one and two connect transcript isoforms with their respective protein translations. Transcripts resulting in identical proteins (shared reading frame, differing only by synonymous mutations) are collapsed and colored by the most abundant transcript (according to read count). Finally, the third column represents clusters of proteins derived from their 3D structures using Foldseek (van Kempen et al. 2024). Ribbons connect protein isoforms from column two, contributing to clusters in column three. Colors of bars in the third column are derived by mixing the colors of ribbons leading into them. Colors of transcript and protein isoforms for the *DAZ* gene are derived from the cluster they contribute to.Three main clusters C1, C2, and C3 represent *DAZ* proteins with two, one, and three RRMs, respectively. (**B**) Molecular visualization of the dominant cluster of five highly ordered YAG proteins created with PyMol (The PyMOL Molecular Graphics System, Version 1.2r3pre, Schrödinger, LLC.). Seven sites highlighted in yellow show evidence of episodic diversifying selection according to the MEME model (Murrell et al. 2012) and are predicted by SIFT (Sim et al. 2012) to affect protein function with high confidence (Table S10).

Despite their high transcript diversity (**Figure 3B**), the highly copious *CDY*, *RBMY*, and *TSPY* families each maintained similar protein folding structure across great apes (**Figure 4A**). Among 24 *CDY* protein isoforms predicted in our data, 21 clustered together. Of the 42 predicted *RBMY* protein structures, 35 formed a single cluster encompassing structures predicted in all great apes (**Figure 4A**), suggesting conserved folding. The *TSPY* gene family produced 34 distinct protein isoforms grouped in four clusters (**Figure 4A**); the largest cluster (C2) had 29 isoforms. For *DAZ*, protein isoforms clustered by the number of RRMs present (one, two, or three), forming three distinct large clusters (**Figure 4A**). The two transcript isoforms of the *HSFY* gene shared one large, highly ordered domain (**Figure 4B**). Additionally, *HSFY* protein isoforms formed a handful of largely species-specific clusters.

Thus, the most copious families—*CDY*, *RBMY*, and *TSPY*—each maintained a conserved protein structure, with most protein isoforms forming a single dominant cluster despite substantial transcript diversity at the sequence (*CDY*), structural (*TSPY*), or both (*RBMY*) levels. For *TSPY*, *DAZ*, and *HSFY*, protein clusters usually corresponded to the same transcript structural isoforms. This collapse was not uniform: in *CDY* and *HSFY*, some protein isoforms derived from identical transcript structural isoforms nonetheless fell into distinct protein structure clusters, demonstrating that both sequence- and structure-level transcript variation contribute to the diversification of protein architecture. But the dominant pattern was conservation—across gene families and species, despite extensive transcript diversity, most predicted protein isoforms collapse into only a few protein structure clusters per YAG family. Yet what maintains this structural conservation remains unexplained.

### Purifying selection preserves structurally ordered proteins

To analyze selection acting on YAG families, we examined the nonsynonymous-to-synonymous rate ratio (*dN/dS*) using `HYPHY` (Kosakovsky Pond et al. 2020) (see Methods for details). Here, we improved our previous analysis (Makova et al. 2024) by including all individual YAG copies (rather than analyzing consensus protein-coding sequences per species) and the closest non-Y homologs, and by applying both branch- and site-specific models (Note S4). We found that, in general, all ampliconic genes evolve under either neutrality or purifying selection (**Table 1**). The evolution of three gene families—*BPY2*, *TSPY*, and *VCY*—was consistent with neutrality (*dN/dS* was not significantly different from 1). Purifying selection was statistically significant for *CDY*, *HSFY*, *RBMY,* and for the RRM domain of *DAZ*.

**Table 1.**
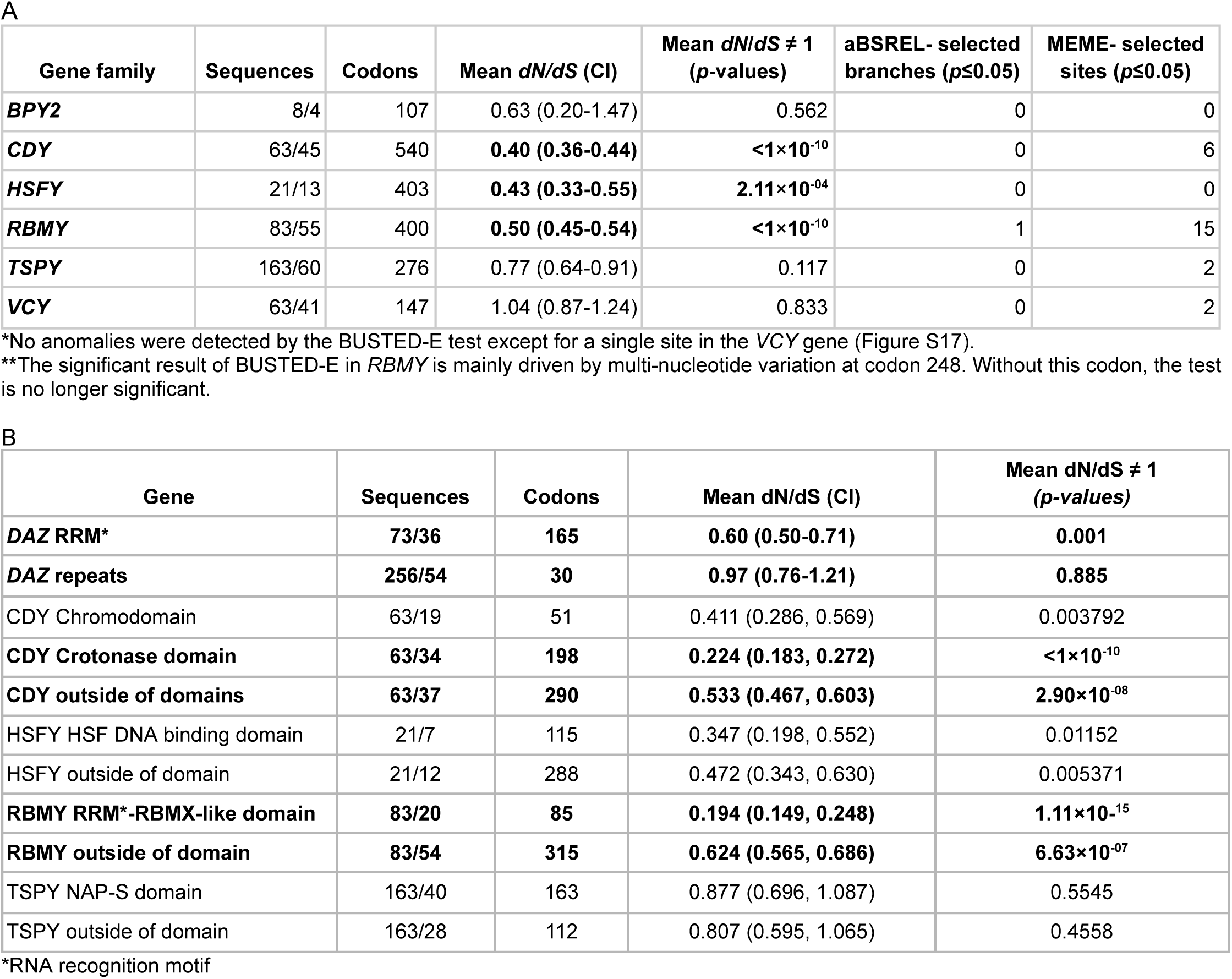
Selection analysis of YAG families. Sequences: Number of all sequences/number of unique sequences considered for the analysis. Codons: number of codons analyzed. *dN*/*dS* CI: 95% confidence intervals for the *dN*/*dS* analysis. Mean *dN*/*dS* ≠ 1: *p*-value for the alternative hypothesis that the alignment-wide and tree-wide *dN*/*dS* is different from 1. Branch heterogeneity: to test for variation in selection pressure across lineages, a local MG94 model (with separate *dN/dS* for each branch) was fitted and compared to the global MG94 model (single *dN/dS* across the tree) using a Likelihood Ratio Test, whose *p*-value is shown. BUSTED-E: the *p*-value for a gene-wide test for episodic diversifying selection. aBSREL: the number of individual branches with evidence of episodic diversifying selection. MEME-selected sites: the number of individual sites that have evidence of episodic diversifying selection. (**A**) Selection analysis performed on the full sequences of YAGs. Statistically significant purifying selection highlighted in bold. (**B**) Domain specific selection of YAGs (*BPY2* and *VCY* do not contain ordered domains. Additional details in Table S9. Genes with significantly higher domain specific selection (non-overlapping confidence intervals) highlighted in bold.

*DAZ* could not be meaningfully analyzed at the whole-gene level because of its complex structure: it cannot be aligned as a single unit because its tandem repeats vary in number, so the RRM and the repeat array were aligned and analyzed separately. We detected evidence of purifying selection within the RRM region, whereas the evolution of *DAZ* repeats was consistent with neutrality (**Table 1**). This result motivated a domain-level analysis across the remaining structured families (*CDY*, *HSFY*, *RBMY*, *TSPY*; Table S9), to test whether selection concentrates on ordered domains more generally.

Following this approach, we performed domain-specific analysis (Table S9A) for all five YAG families with high-confidence predicted structures (Table S9B). As already mentioned, for *DAZ*, structurally ordered RRM and its tandem repeat array evolve under different regimes (purifying selection for the former and neutral evolution for the latter), consistent with prior literature (Bielawski and Yang 2001). For *CDY*, two structured domains were identified by `COLABFOLD` (Mirdita et al. 2022)—crotonase and chromodomain—consistent with prior literature (Dorus et al. 2003) (see Table S9A). Both of them showed significant evidence of purifying selection. The crotonase domain, in particular, exhibited significantly stronger purifying selection (lower *dN/dS* ratio and non-overlapping confidence intervals) than the chromodomain and non-domain region. For *RBMY*, the RRM domain (Elliott 2004) also had significantly stronger purifying selection than the non-domain region. For *HSFY*, the HSF DNA-binding domain exhibited a similar trend, but its selective pressure was not significantly different from that operating on its non-domain region. For *TSPY*, neither the NAP-S domain nor the non-domain region showed statistically significant deviation from neutrality. Thus, across these families, purifying selection on the structurally ordered domains was clearest for *DAZ*, *CDY*, and *RBMY*, weaker and non-significant for *HSFY*, and undetectable for *TSPY*—which retains an ordered predicted structure despite neutral sequence evolution. Thus selection tracks structural order in most, though not all, YAG families.

Using the MEME model (Murrell et al. 2012), we identified 27 sites evolving under episodic positive selection, present in all families but *BPY2* and *HSFY* (*RBMY* had the most, 15, followed by *CDY* with six). Most of these sites were located within the unordered parts of the proteins (17 out of 27; Table S10). Of the 27 sites, however, seven were predicted by `SIFT` (Sim et al. 2012) to affect protein function with high confidence (three in *CDY*, three in *RBMY*, and one in *TSPY*) and six out of seven were located in the ordered regions (Table S10; **Figure 4B**). Thus diversifying selection acts mostly where structure is not changed, but its functional consequences are concentrated at the minority of sites that fall within ordered domains.

Even these functional substitutions, however, are largely buffered: in most families and species, isoforms carrying predicted functional mutations coexist with isoforms derived from non-mutant copies (Figures S10-S15), so the original function is preserved by the latter. The exception is the Met ->Thr substitution at residue 475 for *CDY* in chimpanzee (Figure S16), where the mutant variant is the only form for all gene copies in this species—a case warranting functional investigation.

Together, these results identify protein structure as the level at which YAG function is conserved. The domains predicted to be most ordered are generally those under the strongest purifying selection. Thus, while copy number and transcript identity vary extensively—and DNA sequence somewhat less so—selection maintains the structural integrity of the ordered protein domains that carry out spermatogenic function. This identifies protein structure as the conserved functional level on the otherwise variable Y chromosome.

## Discussion

By integrating T2T genome assemblies, long-read transcriptomics, protein structure prediction, and selection tests—to the best of our knowledge, the first such integration for YAGs—we reconstructed the evolutionary history of these multicopy gene families and evaluated the mechanistic basis and reasons for their survival on the largely non-recombining Y chromosome. Our results resolve a long-standing paradox. We show that Y multicopy genes survive through a system that tolerates extensive variation while constraining its consequences. Although copy number varies widely across species and individuals, and transcript sequence and structure differ across species, this variation is constrained at two levels: gene conversion homogenizes gene copies, and purifying selection preserves protein structure, particularly for the ordered protein domains. This multi-level buffering enables the Y chromosome to tolerate substantial DNA sequence and structural variation due to a lack of recombination and a high mutation rate (Makova and Li 2002; Bergeron et al. 2023), while preserving spermatogenic functions essential for male fertility.

### Palindromes and tandemly repeated arrays represent equivalent evolutionary solutions for YAG homogenization

An important question in Y chromosome biology is whether palindromic or array organization more effectively facilitates gene conversion—the intrachromosomal recombination process that maintains high sequence identity among gene copies while rescuing the functional ones. To address this question thoroughly, one needs to use fully resolved and complete assemblies, as previous Y chromosome assemblies have frequently had palindromes, and likely also arrays, collapsed (Cechova et al. 2020). Here, for this purpose, we utilized ape T2T Y-chromosome assemblies, which have only recently become available (Rhie et al. 2023; Makova et al. 2024). Our analysis revealed that tandem arrays and palindromes achieve comparable levels of sequence homogenization among YAG copies. Moreover, we found that approximately the same number of YAG copies is organized into each of these two structural repeat classes in the apes studied. Our results agree with previous studies demonstrating that YAG copies positioned on opposite arms of the same palindrome are nearly identical (Hallast et al. 2013), whereas those located at nonsymmetric positions within a single palindrome, or on different palindromes, show lower sequence identity (Bhowmick, Satta, and Takahata 2007). Our findings support the notion that gene conversion between tandem direct repeats can proceed through homologous recombination (Grondin, Papadopoulou, and Ouellette 1993; Manz et al. 1993; Xue and Tyler-Smith 2011). Such homogenization on the Y chromosome may occur between sister chromatids. Thus, our results suggest that palindromes and arrays represent alternative but functionally equivalent solutions to the challenge of maintaining genes without interchromosomal recombination.

An alternative explanation is that gene duplication, rather than gene conversion, generates multicopy gene families on the Y. While gene duplication certainly contributes, it cannot alone explain the pattern observed in our phylogenetic trees (**Figure 2C**). We observe genus-specific clades for each multicopy gene family, suggesting sequence homogenization within them. Ancient gene duplications without sequence homogenization would instead produce copy-specific clustering. Without homogenization due to gene conversion, two scenarios could potentially explain our observations, but we consider them to be highly implausible: either the family was already multicopy in the Most Recent Common Ancestor (MRCA) and each species independently retained a different copy while pseudogenizing the others, or the family was single copy in the MRCA and expanded in every species independently after speciation (Katju and Bergthorsson 2013; Cisneros et al. 2026). Both of these scenarios are extremely unlikely, thus we conclude that homogenization via gene conversion is the driving force behind our observations. This conclusion is further strengthened by the fact that homologous recombination during gene conversion itself is known to contribute to variation in copy number (Lange et al. 2009). We note, however, that, in addition to insufficient time for gene conversion, incomplete lineage sorting may contribute to the observed lack of separation of gene copies into species-specific clusters for *Pan* and *Pongo*.

### Transcriptome diversity reflects gene structure and copy number

Our analysis revealed extensive isoform diversity within each YAG family, with the number of distinct transcripts exceeding gene copy number. We demonstrated that this diversity arises from two distinct mechanisms, with their relative contributions varying across gene families. Structural isoform diversity, generated through alternative splicing, correlated with the number of exons per gene. Sequence isoform diversity, reflecting copy-specific variants, correlated with gene copy number. A positive correlation between the number of isoforms and the number of exons was shown in an earlier study (Leung et al. 2021). A correlation between the number of gene copies and the number of sequence isoform variants—which would have been difficult to evaluate accurately before the availability of T2T genomes—to the best of our knowledge, represents a novel finding. For the *DAZ* gene family, transcripts varied in both the number of RRMs and the composition of highly similar *DAZ* repeats. This structural complexity might underlie the functional versatility of *DAZ* in spermatogenesis. In general, the results of our study do not fit any of the three previously described evolutionary models of the interplay between gene duplication and alternative splicing in generating transcript isoform diversity (Iñiguez and Hernández 2017). We propose a new model in which transcript diversity—both at the sequence and structural levels—is relatively conserved among gene copies due to frequent gene conversion on the Y chromosome.

Our use of T2T assemblies for long-read transcript mapping produced fewer, but longer, transcripts than the reference-free IsoCon approach (Table S11) we previously applied (Tomaszkiewicz et al. 2023). This difference can be attributed to the fact that mapping transcripts to the T2T genome results in more accurate full-length transcript reconstruction. We have shown that the availability of such an improved transcriptome affects the estimation of YAG expression levels, revealing some species-specific expression patterns.

### From transcripts to protein clusters

Despite extensive transcript diversity, we predicted only a limited number of protein structure clusters. Indeed, there were fewer predicted protein isoforms than transcript isoforms due to the presence of synonymous variants, and, for all YAG families, protein isoforms formed one or just a handful of clusters with similar structure. This suggests substantial structural conservation among protein isoforms originating from each YAG family.

Previous studies have shown that men with high YAG copy numbers generally exhibit increased YAG transcript levels (Vegesna et al. 2019). However, we previously demonstrated that overall YAG transcript expression levels are conserved across great apes despite rapid copy-number evolution (Vegesna et al. 2020). In contrast, our current study shows some variation in YAG expression levels across the same great ape species (Figure S7). Our protein structure predictions and clustering provide a more complex picture: structurally conserved proteins are expected to perform equivalent functions, so increasing copy number directly increases functional transcript output and likely protein output while maintaining the same molecular activity.

### Selection maintains function through structure conservation in several ampliconic gene families

Our selection analysis revealed a clear dichotomy among YAG families, providing insight into why the Y chromosome has retained these genes despite its overall degeneration. Four gene families—*CDY*, *DAZ*, *HSFY*, and *RBMY* showed evidence of purifying selection. In contrast, the evolution of three other families—*BPY2*, *TSPY*, and *VCY*—was consistent with neutrality. Remarkably, this pattern correlates with protein structure: families evolving under purifying selection encode proteins with highly ordered domains and high AlphaFold confidence scores, whereas *BPY2* might encode ordered proteins but with low confidence scores, and *VCY* encodes intrinsically disordered proteins. *TSPY* represents an intermediate case where protein structure is ordered and conserved despite neutral sequence evolution.

Furthermore, we demonstrate that selection pressure can vary across different functional domains within a single gene. Most notably in *DAZ,* the RRM domains, which mediate RNA binding, showed significant purifying selection and were predicted to be highly ordered, whereas the less ordered *DAZ* repeat region evolved neutrally. This domain-specific pattern, consistent with a prior study in *DAZ* (Bielawski and Yang 2001), suggests that protein-RNA interactions are functionally constrained, whereas *DAZ* repeats are less constrained (or we lack the power to detect selection acting on them due to their short length). In both *CDY* (crotonase domain) and *RBMY* (RRM domain), we observed significantly stronger selection on ordered domains than outside of them. In *HSFY* and TSPY, there was no difference in selection strength between their domains and non-domain regions.

Therefore, highly ordered proteins and domains, in general, evolve under purifying selection, which maintains their structure and function. Disordered proteins and protein domains usually do not experience such sequence constraints and, as a rule, evolve neutrally. The intrinsically disordered proteins encoded by *BPY2* and *VCY* may represent a distinct functional class in which disorder itself is functionally relevant, though their roles in spermatogenesis remain poorly characterized (Tse et al. 2003; Wong et al. 2002; Choi et al. 2007; Shi, Massaia, et al. 2019; Zou et al. 2003; Taguchi et al. 2014). It is also possible that *BPY2* and *VCY* are less functionally constrained because they are on the path to becoming pseudogenes.

### Evolution of YAGs

Our results are consistent with other studies investigating the evolution of YAG families. For instance, YAG families in felids were found to be conserved despite extensive structural variation (Brashear, Raudsepp, and Murphy 2018). Simulations suggested that gene conversion plays an important role in the conservation of essential duplicated genes (Connallon and Clark 2010), and our data support this claim. Moreover, the theory was recently extended to duplicated genes on the Y in the context of Muller’s ratchet (Sakamoto and Innan 2022). In this framework, our results, based on protein structure prediction and clustering, as well as our selection analyses demonstrating purifying selection in several YAG families, strongly argue that gene conversion prevents duplicated genes from losing their function. This scenario is consistent with simulation results under additive dosage effects on fitness (Sakamoto and Innan 2022)—a prediction that should be evaluated in future studies evaluating fitness effects of YAGs with different copy numbers.

Additionally, theoretical models predicted that palindromic organization may accelerate adaptive evolution by increasing targets for beneficial mutations while enabling purging of deleterious variants through gene conversion (Betrán, Demuth, and Williford 2012). In agreement with this prediction, we detected episodic positive selection at multiple sites of YAGs, particularly in *RBMY* and *CDY*, indicating that these gene families experience both purifying and diversifying selection at different positions. Moreover, our `SIFT` (Sim et al. 2012) analysis suggested that sequence variability at several sites evolving under positive selection may affect protein function.

One can draw a parallel between the evolutionary dynamics of ampliconic genes and meiotic drive systems—selfish genetic elements that bias their own transmission (Martí and Larracuente 2023). Sex chromosomes lacking recombination are particularly prone to intragenomic conflict, where segregation distorters trigger arms races between driving and suppressing alleles (Swanepoel and Mueller 2024). The multicopy organization of YAGs may have originated through such conflicts, with gene amplification providing dosage-sensitive responses to drive elements. Indeed, such intragenomic conflicts were suggested to result in genetic redundancy (Martí and Larracuente 2023). Multicopy gene family evolution on primate Y chromosomes shows recurrent amplification and loss at rates higher than those on autosomes, consistent with intragenomic conflict operating at sex chromosomes (Ghenu et al. 2016). Recent work on mouse *t*-haplotypes demonstrates that ampliconic sequences are a defining feature of selfish chromosomes, with amplicon acquisition marking the origins of transmission distortion (Swanepoel et al. 2025).

### Implications for human infertility and great ape conservation

Deletions of ampliconic genes, particularly within the AZF (Azoospermia Factor) regions containing *DAZ*, are established causes of spermatogenic failure associated with human male infertility (Krausz and Abrardo 2025). However, standard genetic testing assesses only the presence or absence of gene families, not copy number, isoform composition, or expression levels (Colaco and Modi 2018). Our finding of transcript diversity among great apes suggests variation in transcriptome expression profiles across men, which may influence spermatogenic efficiency independently of gross deletions. Future clinical studies should consider which isoforms are expressed and at what levels. More precise molecular characterization may enable better prediction of fertility outcomes and inform therapeutic strategies.

All non-human great apes are endangered and rely on captive breeding programs, whose success depends on male fertility. However, the genetic factors contributing to great ape spermatogenesis are not well understood. Our analysis of YAG copy number variation, transcript diversity, and species-specific expression patterns provides a foundation for assessing their reproductive health. Significant inter-individual variation in copy number, observed here in chimpanzees and gorillas, suggests that some males might carry YAG configurations linked to reduced fertility. Establishing the normal range of YAG variation in other species (bonobo and orangutans) will be important in future studies. Furthermore, conservation of YAG protein structures despite sequence divergence indicates that human-based functional assays may be applicable to other great apes, supporting comparative reproductive studies.

### Methodological advances

This study demonstrates the power of combining T2T assemblies with long-read transcriptomics, protein structure prediction, and selection testing for characterizing repetitive gene families. While working towards this goal, we developed the following new methods. First, we developed a *k*-mer-based method to detect tandemly repeated arrays. Second, we developed a method to resolve transcriptome diversity of multicopy genes at both the structural and sequence isoform levels, while incorporating T2T assemblies in the analysis. Third, our approach enabled the characterization of transcripts from the *DAZ* gene family at unprecedented detail, distinguishing highly similar copies with complex internal repeat structures. Previous studies could not fully characterize the repeat composition of individual *DAZ* isoforms. Our approach successfully overcame this limitation and detected novel isoforms, including *DAZ* transcripts with three RRMs in most species, including humans—expanding the known diversity of this clinically important gene family (such isoforms were not detected previously according to https://www.ncbi.nlm.nih.gov/gene/1617, accessed April 24, 2026).

### Limitations

Several limitations should be acknowledged. First, our transcriptomic data are derived from one individual per species, limiting a comprehensive assessment of intraspecific variation in expression patterns. Second, our transcriptome data originate from different individuals than those used for the T2T reference genome reconstruction, preventing a direct one-to-one transcript-to-copy mapping. While acknowledging this limitation, we would like to highlight that collecting matching data would be very challenging. T2T genome sequencing requires a cell line that can be grown to tens of millions of cells, whereas transcriptome characterization would require fresh tissues. In the future, testis cell type-specific lines would have to be established for great apes, which are all endangered (this will be a major undertaking). Third, while `COLABFOLD` (Mirdita et al. 2022) predictions achieve near-experimental accuracy for many protein families (Jumper et al. 2021), predictions may be less reliable for intrinsically disordered proteins and regions. The low confidence scores for *BPY2* and *VCY* likely reflect genuine disorganization rather than prediction failure, but functional implications remain uncertain. Fourth, our analysis focuses on transcript structure and predicted protein properties but does not directly assess protein abundance or functional activity, which may differ from transcript levels due to post-transcriptional regulation. This line of research should be addressed in future work.

## Methods

### Versions of genome assemblies and NCBI annotations

We used the following versions of genome assemblies: GCF_009914755.1-RS_2024_08 (human CHM13+Y), GCF_029281585.2-RS_2024_02 (gorilla), GCF_029289425.2-RS_2024_02 (bonobo), GCF_028858775.2-RS_2024_02 (chimpanzee), GCF_028885655.2-RS_2024_02 (Sumatran orangutan), GCF_028885625.2-RS_2024_02 (Bornean orangutan), and GCF_028878055.3-RS_2024_07 (siamang).

### Assigning genes to gene families

The initial set of gene copies and homologs for all YAGs were determined from gene annotations provided by NCBI for the genome assemblies listed above. In more detail, a `BLASTP` (Camacho et al. 2009) database was created from protein sequences of all protein-coding genes of all species. For each gene, the longest protein isoform was selected. For each species, all protein-coding genes were queried against this database using `BLASTP` (Camacho et al. 2009). Two genes were considered homologs if they reciprocally had at least 50% amino acid sequence identity over at least 35% of their protein lengths (Assis and Bachtrog 2013). Homologous genes were transitively clustered into gene families by identifying connected graph components. If genes (nodes) A and B were identified as homologs (connected by an edge), and so were genes B and C, then genes A, B, and C were assigned to the same gene family (Additional Data File 3).

### Manual curation of individual YAG copies

Highly similar multicopy genes and pseudogenes can be incorrectly annotated and misidentified. In (Makova et al. 2024), we addressed this issue by combining multiple annotation sources and manually curating YAGs in great apes. Since this publication, we have continued working with the NCBI gene annotation team to further curate YAGs of all seven ape species we studied in (Makova et al. 2024) and here. Genes and pseudogenes from the same YAG family and the same species, located within the same palindrome or the same array, were aligned with `MAFFT` v7.526 (Katoh and Standley 2013). When a reading frame in a pseudogene (based on the splice pattern of a homologous protein-coding gene copy) resulted in a transcript without premature stop codons or frameshifts, a change from a pseudogene to a protein-coding gene was proposed. Similarly, truncated copies of annotated protein-coding genes were flagged, and a change from a protein-coding gene to a pseudogene was proposed. These suggestions were reviewed by NCBI (Additional Data File 4) and incorporated into the most recent NCBI Reference Sequence Database (RefSeq). The `GFF` files resulting from such a manual curation of YAGs are available as Additional Data File 5; accession numbers of all protein-coding genes considered are in Table S12.

### Copy number estimation from additional human Y chromosome assemblies

The CDS sequences of all YAG copies annotated on the human reference Y chromosome (HG002) were mapped to the 45 T2T or nearly T2T human Y chromosomes (Hallast et al. 2023) using `MINIMAP2` (Li 2018) in splice-aware mode (`-ax splice`). Sequences of target regions were extracted using `bamtobed` (`BEDTOOLS`) (Quinlan and Hall 2010), translated into protein sequences, and searched against all known human proteins using `BLASTP` (Camacho et al. 2009). As a result, we identified sequences matching YAG proteins. The results of this analysis are summarized in Table S13.

### Copy number estimation from short-read data in chimpanzee and gorilla

`CUTADAPT` (Martin 2011) was used to trim low-quality ends of reads (-q 20) and to trim Illumina TruSeq adaptors from short-read sequencing data analyzed in (Makova et al. 2024) (accession in Table S14). These pre-processed `FASTQ` files were mapped to either the gorilla or chimpanzee T2T reference genomes (versions GCF_029281585.2-RS_2024_02 and GCF_028858775.2-RS_2024_02, respectively) using `BWA-MEM` (Li 2013). `AMPLICONE` (Vegesna et al. 2019) was used to estimate copy numbers of YAG families for each individual. Gene definition files containing the genomic coordinates for each YAG gene copy in the reference ape genomes, as required by `AMPLICONE`, are provided in Table S15.

### Analysis of significant changes in copy numbers along the phylogeny

`CAFE` (Mendes et al. 2021) was used to analyze the evolution of copy number within each YAG family along the phylogeny. We used median copy numbers for species with data for multiple individuals (human, chimpanzee, and gorilla; Tables S5 and S6). For `CAFE`, the Bonferroni-corrected significance threshold of *p* = 0.05/13 = 0.0038 (13 nodes in the tree) was used to identify significant changes in copy number across the phylogeny (Table S4).

### Identification of palindromes and tandemly repeated arrays

Palindromic sequences (Table S16) were identified on the Y chromosome of each species using `LASTZ` (Harris, n.d.) self-alignments, followed by palindrome calling with `PALINDROVER` (Makova et al. 2024) with the following parameters: minimum arm length: 8 kb, minimum sequence identity: 98%, maximum spacer length: 500 kb. In human, a preprocessing step was applied to extract the euchromatic region of the Y chromosome by `samtools faidx` (Li et al. 2009), in order to exclude the highly repetitive heterochromatic region after coordinate 27,449,931 (Rhie et al. 2023). Candidate palindromes with ≥80% overlap with low-complexity sequences, simple repeats, or satellite DNA (based on REPEATMASKER (Smit, Hubley, and Green 2021) annotations) were excluded.

Tandemly repeated arrays on each Y chromosome were identified using `AMPLICOVER` v0.1.1-alpha (https://github.com/makovalab-psu/amplicover). `AMPLICOVER` detects tandemly repeated sequence copies (i.e., arrays) by identifying periodicity in the genomic distance between adjacent identical *k*-mers. Each chromosome was analyzed with *k*=21 and default parameters, via array signal calling, array boundary detection, and unit identification, as implemented in `AMPLICOVER` (Table S17).

### Multiple sequence alignments

Multiple sequence alignments (MSAs) for phylogenetic and selection analyses were built from CDS sequences of all identified protein-coding copies of YAGs and their autosomal and X-chromosome homologs using `MAFFT` v7.526 (Katoh and Standley 2013) with the `--globalpair and --maxiterate 1000` options, in a multi-step manner. For each YAG family, one MSA was built for the Y homologs and an additional one for all non-Y homologs (if present). Next, for each gene family, the MSAs of Y and non-Y homologs were combined into the final MSA using the --`merge` parameter. This approach was used for all genes except for *DAZ* and *RBMY*.

The *DAZ* gene contains an RRM domain (exons 2-6) and a 72-bp (or 90-bp) exon repeated multiple times (Fu et al. 2015), ranging from four copies in the Bornean orangutan to 16 in human (Fu et al. 2015). Variable numbers of *DAZ* repeats between individual gene copies make alignments of full CDSs challenging. Hence, two MSAs were constructed for the *DAZ* gene family: one comprising all RRMs from all gene copies and another one comprising all short *DAZ* repeats from all gene copies.

The *RBMY* gene contains an internal repetitive region, SRGY (Ma et al. 1993; Elliott 2004), which complicates aligning coding sequences. To distinguish between repeat units across homologs, the full genomic sequences (exons and introns) of all *RBMY* homologs were aligned using `MAFFT` (Katoh and Standley 2013). With this approach, introns served as alignment anchors, simplifying the alignment process. The coding sequences of individual gene copies were then extracted. In the final MSA, we considered only exons annotated across all gene copies.

For all YAG families studied, final alignments were produced by trimming both ends of the MSAs, when needed, based on visual inspection. The 5’ end was trimmed to the most common initiation codon, and the 3’ end was trimmed when it was substantially different among homologs. Final alignments are presented in https://github.com/makovalab-psu/YAGs-survival/tree/main/selection_full.

### Phylogenetic analysis

Maximum-likelihood phylogenies from the alignments for each gene family were inferred by `IQ-TREE` v2.3.6 (Minh et al. 2020) using automatic best-fit model selection with `MODELFINDER` (Kalyaanamoorthy et al. 2017). Branch support was assessed with 1,000 ultrafast bootstrap replicates.

### YAG sequence similarity and permutation analysis

Pairwise nucleotide identity between genes of the same YAG family and species was calculated based on the number of mismatching bases (excluding gaps) from a pairwise alignment performed with `MAFFT` (Katoh and Standley 2013) in global mode, only gene pairs with a length difference of <10% relative to the longer sequence were included in the analysis. Copies within both arrays and palindromes were excluded from comparisons, and *DAZ* gene copies were excluded due to their unique structure. Two-sided permutation test with 10,000 permutations was used to compare medians and means between YAGs in the following groups: genes located within the same tandemly repeated array, genes from the same gene family but located on distinct tandemly repeated arrays, genes on opposite arms of a palindrome, genes on the same arm of a palindrome, genes located on different palindromes, and finally genes outside of either palindromes or arrays. 15 comparisons were made and Bonferroni correction of ⍺ = 0.05/15

### RNA sequencing

RNA sequencing was applied genome-wide and separately targeting these specific gene families. Targeted (enriched with hybridization probes for YAGs) long-read Iso-Seq data for testis samples for six species (one sample per species, for all species studied but siamang) were from (Tomaszkiewicz et al. 2023), in which two technical replicates per sample were generated. Whole-genome long (PacBio) and short (Illumina) read expression data for the same testis samples from five non-human great ape species were from (Makova et al. 2024). For the human testis sample (A119, provided by the Cooperative Human Tissue Network (CHTN) under Penn State IRB STUDY00005084), targeted data were generated in (Tomaszkiewicz et al. 2023). Here, we generated an additional untargeted dataset from this sample. To do this, total RNA was isolated using the RNeasy Mini Kit (Qiagen, Germany) according to the manufacturer’s instructions. Tissue was homogenized using a BeadBug 6 homogenizer (Benchmark Scientific, NJ, USA). RNA quantity and integrity were assessed using a TapeStation system (Agilent, CA, USA). For short-read RNA sequencing, libraries were prepared using the Illumina Stranded mRNA Library Prep kit and sequenced in paired-end mode (2×150 nt) on a NextSeq 2000, yielding 123.7 million read pairs. For long-read RNA sequencing, RNA libraries were generated using the Kinnex Full-Length RNA Kit, following the manufacturer’s MAS-Seq protocol. Sequencing was performed on a Sequel IIe system (Pacific Biosciences) using a single SMRT Cell, yielding 21.5 million reads.

Raw reads from both targeted and untargeted PacBio approaches were preprocessed using the Iso-Seq3 pipeline (PacBio) to produce full-length non-chimeric (FLNC) reads. For Illumina short-read RNA-seq, adapters were removed using `TRIMGALORE` (Krueger et al. 2023). Sample information and accession numbers are provided in Table S18.

### Identification of YAG transcript reads in long-read data

PacBio FLNC reads were converted from `BAM` to `FASTA` format using `SAMTOOLS` (Danecek et al. 2021) and aligned to the corresponding species reference genome assemblies (listed above) in the splice-aware mode of `MINIMAP2` (Li 2018). Quality-trimmed Illumina reads, as described in the previous paragraph, were mapped to the corresponding reference genomes using `HISAT2` (Kim et al. 2019), and reads mapping to the Y chromosome were retained.

### Transcriptome assembly and quality control

For each species, a T2T reference-informed transcriptome assembly was constructed using `STRINGTIE` (Shumate et al. 2022) in hybrid mode, integrating long and short reads. The three independently assembled transcript sets per species (two targeted—one per each technical replicate—and one untargeted) were compared using `GFFCOMPARE` (Pertea and Pertea 2020). For each species, we identified a union of transcripts from these three sets, which was used in subsequent analyses. Transcripts were evaluated using `SQANTI3` (Pardo-Palacios et al. 2024), and transcripts flagged as likely artifacts or degradation products were excluded.

### Assignment of transcripts to YAG families

For each species, each transcript (represented as GFF coordinates) was assigned to a YAG family using one of two methods. In the first method, we searched for overlap between transcript coordinates and annotated YAG loci in the reference genomes using a custom script (annotation_based_on_region_intersection.R available on GitHub). If no overlap was found, we applied the second method. Here, we used transcript coordinates (in GFF format) to extract transcript sequences from the reference genome, using `GFFREAD` (Pertea and Pertea 2020). Next, we aligned the extracted sequences against a database of known YAG gene sequences we built from great ape T2T assembly annotations (extract_gene_families.py available on GitHub), using `LASTZ` (Harris, n.d.). These alignments were filtered to retain matches with ≥80% sequence identity over ≥100 bp, and used to assign the transcripts to YAG families.

### ORF prediction

For all YAG transcripts identified in the previous step, ORFs were predicted in transcripts using `GETORF` from the `EMBOSS` suite (Rice, Longden, and Bleasby 2000), requiring a minimum size of 50 amino acids for the resulting protein. For additional validation, we used `BLASTP` (Camacho et al. 2009) to align the predicted ORF sequences against a reference database of great ape YAG proteins (create_protein_db.txt available on GitHub) that we constructed to include experimentally validated and predicted proteins for the seven YAG families, as available at NCBI.

### Comparison of isoform structures

To identify alternative splicing patterns for each gene family, transcripts with ORFs were aligned to the longest human gene copy in each YAG family using `ULTRA` (Sahlin and Mäkinen 2021). This has allowed us to project all transcripts from the same family onto a common coordinate system. Structural isoforms were identified by converting alignments to the `GFF` format with `BAMTOBED` (Quinlan and Hall 2010), and comparing each transcript’s exon-intron architecture against the full set using `GFFCOMPARE` (Pertea and Pertea 2020). Transcripts with identical exon-intron structure were assigned the same isoform identifier.

### Identification of sequence isoforms from Iso-Seq reads

Transcript sequence diversity in YAG families was determined by identifying sequence-specific signatures. For this analysis, YAG-specific Iso-Seq reads were used (see the “Identification of YAG transcript reads in long-read data” section).

For each of the *BPY2, CDY, HSFY, RBMY, TSPY,* and *VCY* gene families, first, gene-family-specific transcript reads were aligned to the longest copy for each gene family in each species using the splice-aware mode of `MINIMAP2` (Li 2018) to enable comparisons across all copies within a species. `FREEBAYES` (Garrison and Marth 2012) was used to perform variant calling on the alignments. Variants with transcript read depth greater than 30× were used in the signature analysis. Variable positions among reference gene copies were also noted. Second, each transcript read was processed to determine whether a variant was present or absent at positions defined in the previous step. The combination of variants specified a transcript sequence signature, or a fingerprint. The results of this analysis were compiled into count tables summarizing the frequency distributions of signatures across samples (Additional Data File 6). Third, for these transcripts, which now have assigned sequence signatures, we identified splicing patterns and matched them to structural isoforms as defined in “Comparison of isoform structures”. To do this, transcript reads (with signatures) from each species were mapped to the corresponding T2T reference genome assemblies using `MINIMAP2` (Li 2018). As a result of this step, each transcript read had a previously assigned structural identifier. Summarizing these results, we built tables with rows representing sequence signatures and columns representing transcript structural isoforms (Additional Data File 1). Finally, we filtered and translated the observed transcript isoforms sequences into protein isoforms using the following three steps. First, a transcript isoform was considered only if it was represented by at least 2% of the total number of full-length reads identified for the given gene family within a species, or by at least five reads, whichever was higher. Second, we required that each transcript isoform be supported by at least two of the three independently assembled transcript sets (see section “Transcriptome assembly and quality control”). Third, to exclude sequencing errors, reads representing each retained transcript isoform were collected, and their consensus sequences were called using `SPOA` (Vaser et al. 2017). An ORF within each transcript isoform was identified with `ORF-FINDER` (Rombel et al. 2002). Finally, the retrieved ORFs were translated using the `BIOPYTHON` (Cock et al. 2009) library, using the default codon table—NCBI Genetic Code Table 1 (https://www.ncbi.nlm.nih.gov/Taxonomy/Utils/wprintgc.cgi).

For the *DAZ* gene family, first, transcript reads containing at least part of the RRM domain were identified. Given the composition of *DAZ* genes 5’(RRM)*_N_*(*DAZ* repeat)*_M_*3’, where *N* is the number of RRMs (*N*=1-3) and *M* is the number of *DAZ* repeats, and considering reads are sequences from the poly A tail (3’ -> 5’), this ensures that the retained reads contain the full sequence of *DAZ* repeats. Second, for each species, all *DAZ* repeat sequences were collected, and an MSA of just these sequences was produced using `MAFFT` (Katoh and Standley 2013) in global alignment mode. Third, `HMMBUILD` (`HMMER` 3.4, (Eddy 2011)) was used to build the Hidden Markov Model (`HMM`) profile of *DAZ* repeats. Fourth, utilizing this `HMM` profile, `HMMSEARCH` (`HMMER` 3.4, (Eddy 2011)) was used to identify individual *DAZ* repeats in each read. Fifth, unique *DAZ* repeat sequences occurring within each species were collected and sorted by the number of occurrences (Additional Data File 2). The sorted list of identified *DAZ* repeats (Additional Data File 2) contains a small number of high-abundance *DAZ* repeat sequences and a long list of low-abundance sequences. For each species, the top *N* most abundant sequences were considered true repeat sequences. *N* was manually set for each species after visual inspection of the count tables (Additional Data File 2) (to identify a natural breakpoint in abundance between signal and noise) as follows: 10 for human and gorilla, 20 for bonobo and chimpanzee, 10 for Bornean orangutan, and 3 for Sumatran orangutan. The ‘true’ (i.e., high-abundance repeat sequences) were labeled A, B, C, etc. The remaining low-abundance sequences in the count tables received the label of the closest (according to the Hamming distance) true repeat sequence. Sixth, a signature consisting of a pattern of *DAZ* repeat labels based on the repeat content was constructed for each read. Seventh, a consensus sequence was called from reads clustered by their *DAZ* repeat content signatures using `SPOA` (Vaser et al. 2017), and a protein sequence was extracted from an open reading frame identified by `ORF-FINDER` (Rombel et al. 2002). The scripts for variant calling, signature extraction, mapping, splice pattern matching, alignment, HMM, and consensus calling are available on GitHub (https://github.com/makovalab-psu/reconstruct-isoform-sequence).

### Comparison of reference-augmented structural isoforms with a set of previously *de novo* assembled structural isoforms

For this comparison, we used transcripts from the section “Identification of sequence isoforms from Iso-Seq reads.”) from Table S11. Here, we identified 180 replicate-supported transcripts, whereas in (Tomaszkiewicz et al. 2023) we identified 1,510 transcripts. On average, transcripts identified by (Tomaszkiewicz et al. 2023) were shorter (mean length of 820 bp in (Tomaszkiewicz et al. 2023), as compared to those in our reference-based approach (mean length = 1,215 bp). Considering our approach filters reads based on read length, a fairer comparison of methods is provided in (Note S3).

### RNA-seq expression quantification

Expression of YAG transcripts was quantified from testis short RNA-seq reads generated in (Makova et al. 2024) (Table S18) using `SALMON` v1.10.3 (Patro et al. 2017). To assess the effect of using a genome reference on YAG expression estimates, two transcriptomes were constructed per species. The first transcriptome consisted of the NCBI RefSeq gene annotations. The second, reference-augmented transcriptome was constructed by adding species-specific novel YAG transcript sequences (which we identified by incorporating T2T reference genome assemblies into analysis, see sections “Identification of YAG transcript reads from long-read data” through “Identification of sequence isoforms from Iso-Seq reads.”) to the standard RefSeq reference. `SALMON` indices were built with a *k*-mer size of 31.

Quantification was performed using selective alignment (`--validateMappings`), correcting for sequence-specific biases (`--seqBias`), and fragment-level GC biases (`--gcBias`). Expression differences between standard and reference-augmented transcriptomes were reported in read counts and in TPM. Fold differences were computed as log2((augmented + 1) / (standard + 1)), with a pseudocount of 1 to handle zero counts, and are reported in Table S7.

### Protein folding predictions and clustering

All transcripts from the previous step that contained an ORF were used for protein structure prediction, which was then used in the following steps. First, by collapsing transcripts differing by synonymous variants, we identified and included only unique predicted protein isoforms. Second, `COLABFOLD` v1.5.5 (Mirdita et al. 2022) was run to produce structure predictions, MSAs of related proteins, and AlphaFold2 structure prediction confidence measures (pLDDT). Third, the predicted protein structures were clustered using `FOLDSEEK` commit 10.941cd33 (van Kempen et al. 2024) with default parameters.

### Selection analysis

All alignments from the “Multiple sequence alignments” step were examined for the presence of premature stop codons and the presence of identical gene copies—only unique copies were retained, using `HYPHY cln` (Kosakovsky Pond et al. 2020). For each gene family, the alignment and the phylogenetic trees (from the “Phylogenetic analysis” step above) were fitted to the standard MG94 (Muse and Gaut 1994) model to estimate a mean *dN*/*dS* and its confidence intervals using `HYPHY`. BUSTED-E within `HYPHY` was used to detect episodic diversifying selection and to identify errors caused by misalignments. Each gene family was tested with the following tests:

1. Mean *dN*/*dS* ≠ 1: a likelihood ratio test under the MG94 global model.
2. Branch heterogeneity: a test for variation in selection pressure across branches, via fitting of a local MG94 model (separate *dN*/*dS* for each branch) and comparing it to the global MG94 model (single *dN/dS* across the phylogeny) using a Likelihood Ratio Test.
3. BUSTED-E: a gene-wide test for episodic diversifying selection that also estimates whether or not there is a non-zero error component.
4. aBSREL-selected branches: a test for evidence of episodic diversifying selection on individual branches.
5. MEME-selected sites: a test of episodic diversifying selection on individual sites.

For each of the *CDY, HSFY, RBMY,* and *TSPY* genes, we extracted protein sequences corresponding to known functional domains (which largely correspond to the predicted structured domains), using `CD-SEARCH` at NCBI. As a result, we extracted the chromodomain and crotonase domain for *CDY*, the HSF DNA-binding domain for *HSFY*, the RRM domain for *RBMY*, and the NAP-S domain for *TSPY*, as well as the remaining “no domain” sequence (Table S9A). Next, we aligned these retrieved partial protein sequences to the consensus MSA sequences of the corresponding nucleotide alignments using Biophython pairwise aligner (Cock et al. 2009). Data and code are in https://github.com/makovalab-psu/YAGs-survival/tree/main/selection_on_domains). For each region, we executed the same selection analysis steps as for the full genic region above.

### Assessment of functional impact of non-synonymous substitutions

For each gene, a consensus protein sequence was derived from an MSA of coding sequences across species (see section “Multiple sequence alignments”). Protein sequence and substitutions of interest were submitted to the `SIFT` web server (<u>sift.bii.a-star.edu.sg</u>, (Sim et al. 2012)). `SIFT` (Kumar, Henikoff, and Ng 2009) predictions and confidence scores were retrieved from the server output and formatted into a table format. Substitutions were classified as “Affect Protein Function” (score < 0.05) or “Tolerated” (score ≥ 0.05). Some predictions were flagged as low confidence (insufficient sequence diversity). All code used for processing of `SIFT` results is available at https://github.com/makovalab-psu/YAGs-survival/tree/main/SIFT_annotation.

## Acknowledgements

We would like to thank Françoise Thibaud-Nissen, Kelly McGarvey, and Shashikant Pujar of the NCBI gene annotation team for feedback, assistance, and review during gene curation. Next, we would like to thank Linnéa Smeds for careful reading of the manuscript and helpful comments. Finally, we would like to acknowledge Kerry Hair and Dan Hannon for their help with Illumina mRNA library preparation sequencing and Kinnex (MAS-seq), respectively. We also thank the Huck Institutes’ Genomics Core Facility (RRID:SCR_023645) for use of the Illumina NextSeq 2000 and PacBio Sequel IIe. Human testis tissue sample used for generating expression sequencing data was provided by the NCI Cooperative Human Tissue Network (CHTN). Other investigators may have received specimens from the same tissue specimen. The research in Kateryna D. Makova’s laboratory is supported by the grant R35GM151945 from the NIH and by the Willaman Chair Endowment Fund from the Eberly College of Science. Computations were performed at the Penn State Institute of Computational Data Sciences (RRID:SCR_025154), which provided access to computational research infrastructure within the Roar Core Facility (RRID:SCR_026424). Sergei L. Kosakovsky Pond acknowledges support from NIH/NIGMS (GM151683) and NSF DBI (2419522). Martin Steinegger acknowledges support by the National Research Foundation of Korea grants (2020M3A9G7103933, RS-2021-NR061659 and RS-2021-NR056571, RS-2024-00396026), Novo Nordisk Foundation (NNF24SA0092560), and Creative-Pioneering Researchers Program through Seoul National University.

## References

Assis, Raquel, and Doris Bachtrog. 2013. “Neofunctionalization of Young Duplicate Genes in *Drosophila*.” Proceedings of the National Academy of Sciences of the United States of America 110 (43): 17409–14.

Bergeron, Lucie A., Søren Besenbacher, Jiao Zheng, Panyi Li, Mads Frost Bertelsen, Benoit Quintard, Joseph I. Hoffman, Zhipeng Li, Judy St Leger, Changwei Shao, Josefin Stiller, M. Thomas, P. Gilbert, Mikkel H. Schierup, and Guojie Zhang. 2023. “Evolution of the Germline Mutation Rate across Vertebrates.” Nature 615 (7951): 285–91.

Betrán, Esther, Jeffery P. Demuth, and Anna Williford. 2012. “Why Chromosome Palindromes?” International Journal of Evolutionary Biology 2012 (July):207958.

Bhowmick, Bejon Kumar, Yoko Satta, and Naoyuki Takahata. 2007. “The Origin and Evolution of Human Ampliconic Gene Families and Ampliconic Structure.” Genome Research 17 (4): 441–50.

Bielawski, J. P., and Z. Yang. 2001. “Positive and Negative Selection in the DAZ Gene Family.” Molecular Biology and Evolution 18 (4): 523–29.

Brashear, Wesley A., Terje Raudsepp, and William J. Murphy. 2018. “Evolutionary Conservation of Y Chromosome Ampliconic Gene Families despite Extensive Structural Variation.” Genome Research 28 (12): 1841–51.

Camacho, Christiam, George Coulouris, Vahram Avagyan, Ning Ma, Jason Papadopoulos, Kevin Bealer, and Thomas L. Madden. 2009. “BLAST+: Architecture and Applications.” BMC Bioinformatics 10 (1): 421.

Cechova Monika, Rahulsimham Vegesna, Marta Tomaszkiewicz, Robert S. Harris, Di Chen, Samarth Rangavittal, Paul Medvedev, and Kateryna D. Makova. 2020. “Dynamic Evolution of Great Ape Y Chromosomes.” Proceedings of the National Academy of Sciences of the United States of America 117 (42): 26273–80.

Charlesworth, Brian. 2003. “The Organization and Evolution of the Human Y Chromosome.” Genome Biology 4 (9): 226.

Choi, Jin, Eitetsu Koh, Hiromi Suzuki, Yuji Maeda, Atsumi Yoshida, and Mikio Namiki. 2007. “Alu Sequence Variants of the BPY2 Gene in Proven Fertile and Infertile Men with Sertoli Cell-Only Phenotype: Alu Sequence Variants of BPY2.” International Journal of Urology: Official Journal of the Japanese Urological Association 14 (5): 431–35.

Cisneros, Angel F., Soham Dibyachintan, Frédéric Bédard, Simon Aubé, Pascale Lemieux, and Christian R. Landry. 2026. “Evolutionary Causes and Consequences of Gene Duplication.” Nature Reviews. Genetics 27 (7): 512–29.

Cock Peter J. A., Tiago Antao, Jeffrey T. Chang, Brad A. Chapman, Cymon J. Cox, Andrew Dalke, Iddo Friedberg, Thomas Hamelryck, Frank Kauff, Bartek Wilczynski, and Michiel J. L. de Hoon. 2009. “Biopython: Freely Available Python Tools for Computational Molecular Biology and Bioinformatics.” *Bioinformatics (Oxford*, England) 25 (11): 1422–23.

Colaco, Stacy, and Deepak Modi. 2018. “Genetics of the Human Y Chromosome and Its Association with Male Infertility.” Reproductive Biology and Endocrinology: RB&E 16 (1): 14.

Connallon, Tim, and Andrew G. Clark. 2010. “Gene Duplication, Gene Conversion and the Evolution of the Y Chromosome.” Genetics 186 (1): 277–86.

Danecek, Petr, James K. Bonfield, Jennifer Liddle, John Marshall, Valeriu Ohan, Martin O. Pollard, Andrew Whitwham, Thomas Keane, Shane A. McCarthy, Robert M. Davies, and Heng Li. 2021. “Twelve Years of SAMtools and BCFtools.” GigaScience 10 (2). 10.1093/gigascience/giab008.

Davis, Jamie K., Pamela J. Thomas, NISC Comparative Sequencing Program, and James W. Thomas. 2010. “A W-Linked Palindrome and Gene Conversion in New World Sparrows and Blackbirds.” Chromosome Research: An International Journal on the Molecular, Supramolecular and Evolutionary Aspects of Chromosome Biology 18 (5): 543–53.

Dorus, Steve, Sandra L. Gilbert, Michele L. Forster, Robert J. Barndt, and Bruce T. Lahn. 2003. “The CDY-Related Gene Family: Coordinated Evolution in Copy Number, Expression Profile and Protein Sequence.” Human Molecular Genetics 12 (14): 1643–50.

Eddy, Sean R. 2011. “Accelerated Profile HMM Searches.” PLoS Computational Biology 7 (10): e1002195.

Elliott David J. 2004. “The Role of Potential Splicing Factors Including RBMY, RBMX, hnRNPG-T and STAR Proteins in Spermatogenesis.” International Journal of Andrology 27 (6): 328–34.

Fu, Xia-Fei, Shun-Feng Cheng, Lin-Qing Wang, Shen Yin, Massimo De Felici, and Wei Shen. 2015. “DAZ Family Proteins, Key Players for Germ Cell Development.” International Journal of Biological Sciences 11 (10): 1226–35.

Garrison, Erik, and Gabor Marth. 2012. “Haplotype-Based Variant Detection from Short-Read Sequencing.” arXiv [q-bio.GN]. 10.48550/ARXIV.1207.3907.

Geraldes, Armando, Teri Rambo, Rod A. Wing, Nuno Ferrand, and Michael W. Nachman. 2010. “Extensive Gene Conversion Drives the Concerted Evolution of Paralogous Copies of the SRY Gene in European Rabbits.” Molecular Biology and Evolution 27 (11): 2437–40.

Ghenu, Ana-Hermina, Benjamin M. Bolker, Don J. Melnick, and Ben J. Evans. 2016. “Multicopy Gene Family Evolution on Primate Y Chromosomes.” BMC Genomics 17 (1): 157.

Grondin, K., B. Papadopoulou, and M. Ouellette. 1993. “Homologous Recombination between Direct Repeat Sequences Yields P-Glycoprotein Containing Amplicons in Arsenite Resistant Leishmania.” Nucleic Acids Research 21 (8): 1895–1901.

Hallast, Pille, Patricia Balaresque, Georgina R. Bowden, Stéphane Ballereau, and Mark A. Jobling. 2013. “Recombination Dynamics of a Human Y-Chromosomal Palindrome: Rapid GC-Biased Gene Conversion, Multi-Kilobase Conversion Tracts, and Rare Inversions.” PLoS Genetics 9 (7): e1003666.

Hallast, Pille, Peter Ebert, Mark Loftus, Feyza Yilmaz, Peter A. Audano, Glennis A. Logsdon, Marc Jan Bonder, Weichen Zhou, Wolfram Höps, Kwondo Kim, Chong Li, Savannah J. Hoyt, Philip C. Dishuck, David Porubsky, Fotios Tsetsos, Jee Young Kwon, Qihui Zhu, Katherine M. Munson, Patrick Hasenfeld, William T. Harvey, Alexandra P. Lewis, Jennifer Kordosky, Kendra Hoekzema, Human Genome Structural Variation Consortium (HGSVC), Rachel J. O’Neill, Jan O. Korbel, Chris Tyler-Smith, Evan E. Eichler, Xinghua Shi, Christine R. Beck, Tobias Marschall, Miriam K. Konkel, and Charles Lee. 2023. “Assembly of 43 Human Y Chromosomes Reveals Extensive Complexity and Variation.” Nature 621 (7978): 355–64.

Harris, Bob. n.d. Lastz: Program for Aligning DNA Sequences, a Pairwise Aligner. Github. Accessed December 12, 2024. https://github.com/lastz/lastz.

Hobza, Roman, Martina Lengerova, Julia Svoboda, Hana Kubekova, Eduard Kejnovsky, and Boris Vyskot. 2006. “An Accumulation of Tandem DNA Repeats on the Y Chromosome in Silene Latifolia during Early Stages of Sex Chromosome Evolution.” Chromosoma 115 (5): 376–82.

Huang, Zhen, Zaoxu Xu, Hao Bai, Yongji Huang, Na Kang, Xiaoting Ding, Jing Liu, Haoran Luo, Chentao Yang, Wanjun Chen, Qixin Guo, Lingzhan Xue, Xueping Zhang, Li Xu, Meiling Chen, Honggao Fu, Youling Chen, Zhicao Yue, Tatsuo Fukagawa, Shanlin Liu, Guobin Chang, and Luohao Xu. 2023. “Evolutionary Analysis of a Complete Chicken Genome.” Proceedings of the National Academy of Sciences of the United States of America 120 (8): e2216641120.

Hughes, Jennifer F., Helen Skaletsky, and David C. Page. 2012. “Sequencing of Rhesus Macaque Y Chromosome Clarifies Origins and Evolution of the DAZ (Deleted in AZoospermia) Genes.” *BioEssays: News and Reviews in Molecular*, Cellular and Developmental Biology 34 (12): 1035–44.

Iñiguez, Luis P., and Georgina Hernández. 2017. “The Evolutionary Relationship between Alternative Splicing and Gene Duplication.” Frontiers in Genetics 8 (February):14.

Jumper John, Richard Evans, Alexander Pritzel, Tim Green, Michael Figurnov, Olaf Ronneberger, Kathryn Tunyasuvunakool, Russ Bates, Augustin Žídek, Anna Potapenko, Alex Bridgland, Clemens Meyer, Simon A. A. Kohl, Andrew J. Ballard, Andrew Cowie, Bernardino Romera-Paredes, Stanislav Nikolov, Rishub Jain, Jonas Adler, Trevor Back, Stig Petersen, David Reiman, Ellen Clancy, Michal Zielinski, Martin Steinegger, Michalina Pacholska, Tamas Berghammer, Sebastian Bodenstein, David Silver, Oriol Vinyals, Andrew W. Senior, Koray Kavukcuoglu, Pushmeet Kohli, and Demis Hassabis. 2021. “Highly Accurate Protein Structure Prediction with AlphaFold.” Nature 596 (7873): 583–89.

Kalyaanamoorthy Subha, Bui Quang Minh, Thomas K. F. Wong, Arndt von Haeseler, and Lars S. Jermiin. 2017. “ModelFinder: Fast Model Selection for Accurate Phylogenetic Estimates.” Nature Methods 14 (6): 587–89.

Katju, Vaishali, and Ulfar Bergthorsson. 2013. “Copy-Number Changes in Evolution: Rates, Fitness Effects and Adaptive Significance.” Frontiers in Genetics 4 (December):273.

Katoh, Kazutaka, and Daron M. Standley. 2013. “MAFFT Multiple Sequence Alignment Software Version 7: Improvements in Performance and Usability.” Molecular Biology and Evolution 30 (4): 772–80.

Kempen, Michel van, Stephanie S. Kim, Charlotte Tumescheit, Milot Mirdita, Jeongjae Lee, Cameron L. M. Gilchrist, Johannes Söding, and Martin Steinegger. 2024. “Fast and Accurate Protein Structure Search with Foldseek.” Nature Biotechnology 42 (2): 243–46.

Kim, Daehwan, Joseph M. Paggi, Chanhee Park, Christopher Bennett, and Steven L. Salzberg. 2019. “Graph-Based Genome Alignment and Genotyping with HISAT2 and HISAT-Genotype.” Nature Biotechnology 37 (8): 907–15.

Kosakovsky Pond, Sergei L., Art F. Y. Poon, Ryan Velazquez, Steven Weaver, N. Lance Hepler, Ben Murrell, Stephen D. Shank, Brittany Rife Magalis, Dave Bouvier, Anton Nekrutenko, Sadie Wisotsky, Stephanie J. Spielman, Simon D. W. Frost, and Spencer V. Muse. 2020. “HyPhy 2.5-A Customizable Platform for Evolutionary Hypothesis Testing Using PHYlogenies.” Molecular Biology and Evolution 37 (1): 295–99.

Krausz, Csilla, and Chiara Abrardo. 2025. “The Y Chromosome: Male Reproduction and beyond.” Fertility and Sterility 123 (6): 921–32.

Krueger, Felix, Frankie James, Phil Ewels, Ebrahim Afyounian, Michael Weinstein, Benjamin Schuster-Boeckler, Gert Hulselmans, and sclamons. 2023. FelixKrueger/TrimGalore: v0.6.10 - Add Default Decompression Path. Zenodo. 10.5281/ZENODO.7598955.

Kumar, Prateek, Steven Henikoff, and Pauline C. Ng. 2009. “Predicting the Effects of Coding Non-Synonymous Variants on Protein Function Using the SIFT Algorithm.” Nature Protocols 4 (7): 1073–81.

Lahn, B. T., and D. C. Page. 1997. “Functional Coherence of the Human Y Chromosome.” *Science (New York*, N.Y.) 278 (5338): 675–80.

Lange Julian, Helen Skaletsky, Saskia K. M. van Daalen, Stephanie L. Embry, Cindy M. Korver, Laura G. Brown, Robert D. Oates, Sherman Silber, Sjoerd Repping, and David C. Page. 2009. “Isodicentric Y Chromosomes and Sex Disorders as Byproducts of Homologous Recombination That Maintains Palindromes.” Cell 138 (5): 855–69.

Leung, Szi Kay, Aaron R. Jeffries, Isabel Castanho, Ben T. Jordan, Karen Moore, Jonathan P. Davies, Emma L. Dempster, Nicholas J. Bray, Paul O’Neill, Elizabeth Tseng, Zeshan Ahmed, David A. Collier, Erin D. Jeffery, Shyam Prabhakar, Leonard Schalkwyk, Connor Jops, Michael J. Gandal, Gloria M. Sheynkman, Eilis Hannon, and Jonathan Mill. 2021. “Full-Length Transcript Sequencing of Human and Mouse Cerebral Cortex Identifies Widespread Isoform Diversity and Alternative Splicing.” Cell Reports 37 (7): 110022.

Li, Heng. 2013. “Aligning Sequence Reads, Clone Sequences and Assembly Contigs with BWA-MEM.” *arXiv [q-bio.GN]*. arXiv. http://arxiv.org/abs/1303.3997.

Li, Heng. 2018. “Minimap2: Pairwise Alignment for Nucleotide Sequences.” Bioinformatics (Oxford, England) 34 (18): 3094–3100.

Li, Heng, Bob Handsaker, Alec Wysoker, Tim Fennell, Jue Ruan, Nils Homer, Gabor Marth, Goncalo Abecasis, Richard Durbin, and 1000 Genome Project Data Processing Subgroup. 2009. “The Sequence Alignment/Map Format and SAMtools.” Bioinformatics (Oxford, England) 25 (16): 2078–79.

Lucotte, Elise A., Laurits Skov, Jacob Malte Jensen, Moisès Coll Macià, Kasper Munch, and Mikkel H. Schierup. 2018. “Dynamic Copy Number Evolution of X- and Y-Linked Ampliconic Genes in Human Populations.” Genetics 209 (3): 907–20.

Ma, K., J. D. Inglis, A. Sharkey, W. A. Bickmore, R. E. Hill, E. J. Prosser, R. M. Speed, E. J. Thomson, M. Jobling, and K. Taylor. 1993. “A Y Chromosome Gene Family with RNA-Binding Protein Homology: Candidates for the Azoospermia Factor AZF Controlling Human Spermatogenesis.” Cell 75 (7): 1287–95.

Makova, Kateryna D., and Wen-Hsiung Li. 2002. “Strong Male-Driven Evolution of DNA Sequences in Humans and Apes.” Nature 416 (6881): 624–26.

Makova, Kateryna D., Brandon D. Pickett, Robert S. Harris, Gabrielle A. Hartley, Monika Cechova, Karol Pal, Sergey Nurk, Dongahn Yoo, Qiuhui Li, Prajna Hebbar, Barbara C. McGrath, Francesca Antonacci, Margaux Aubel, Arjun Biddanda, Matthew Borchers, Erich Bornberg-Bauer, Gerard G. Bouffard, Shelise Y. Brooks, Lucia Carbone, Laura Carrel, Andrew Carroll, Pi-Chuan Chang, Chen-Shan Chin, Daniel E. Cook, Sarah J. C. Craig, Luciana de Gennaro, Mark Diekhans, Amalia Dutra, Gage H. Garcia, Patrick G. S. Grady, Richard E. Green, Diana Haddad, Pille Hallast, William T. Harvey, Glenn Hickey, David A. Hillis, Savannah J. Hoyt, Hyeonsoo Jeong, Kaivan Kamali, Sergei L. Kosakovsky Pond, Troy M. LaPolice, Charles Lee, Alexandra P. Lewis, Yong-Hwee E. Loh, Patrick Masterson, Kelly M. McGarvey, Rajiv C. McCoy, Paul Medvedev, Karen H. Miga, Katherine M. Munson, Evgenia Pak, Benedict Paten, Brendan J. Pinto, Tamara Potapova, Arang Rhie, Joana L. Rocha, Fedor Ryabov, Oliver A. Ryder, Samuel Sacco, Kishwar Shafin, Valery A. Shepelev, Viviane Slon, Steven J. Solar, Jessica M. Storer, Peter H. Sudmant, Sweetalana, Alex Sweeten, Michael G. Tassia, Françoise Thibaud-Nissen, Mario Ventura, Melissa A. Wilson, Alice C. Young, Huiqing Zeng, Xinru Zhang, Zachary A. Szpiech, Christian D. Huber, Jennifer L. Gerton, Soojin V. Yi, Michael C. Schatz, Ivan A. Alexandrov, Sergey Koren, Rachel J. O’Neill, Evan E. Eichler, and Adam M. Phillippy. 2024. “The Complete Sequence and Comparative Analysis of Ape Sex Chromosomes.” Nature 630 (8016): 401–11.

Manz, E., F. Schnieders, A. M. Brechlin, and J. Schmidtke. 1993. “TSPY-Related Sequences Represent a Microheterogeneous Gene Family Organized as Constitutive Elements in DYZ5 Tandem Repeat Units on the Human Y Chromosome.” Genomics 17 (3): 726–31.

Marais, Gabriel A. B., Paulo R. A. Campos, and Isabel Gordo. 2010. “Can Intra-Y Gene Conversion Oppose the Degeneration of the Human Y Chromosome? A Simulation Study.” Genome Biology and Evolution 2 (0): 347–57.

Martí, Emiliano, and Amanda M. Larracuente. 2023. “Genetic Conflict and the Origin of Multigene Families: Implications for Sex Chromosome Evolution.” *Proceedings*. Biological Sciences 290 (2010): 20231823.

Martin, Marcel. 2011. “Cutadapt Removes Adapter Sequences from High-Throughput Sequencing Reads.” EMBnet.journal 17 (1): 10.

Mendes, Fábio K., Dan Vanderpool, Ben Fulton, and Matthew W. Hahn. 2021. “CAFE 5 Models Variation in Evolutionary Rates among Gene Families.” *Bioinformatics (Oxford*, England) 36 (22-23): 5516–18.

Minh, Bui Quang, Heiko A. Schmidt, Olga Chernomor, Dominik Schrempf, Michael D. Woodhams, Arndt von Haeseler, and Robert Lanfear. 2020. “IQ-TREE 2: New Models and Efficient Methods for Phylogenetic Inference in the Genomic Era.” Molecular Biology and Evolution 37 (5): 1530–34.

Mirdita, Milot, Konstantin Schütze, Yoshitaka Moriwaki, Lim Heo, Sergey Ovchinnikov, and Martin Steinegger. 2022. “ColabFold: Making Protein Folding Accessible to All.” Nature Methods 19 (6): 679–82.

Murrell, Ben, Joel O. Wertheim, Sasha Moola, Thomas Weighill, Konrad Scheffler, and Sergei L. Kosakovsky Pond. 2012. “Detecting Individual Sites Subject to Episodic Diversifying Selection.” PLoS Genetics 8 (7): e1002764.

Muse, S. V., and B. S. Gaut. 1994. “A Likelihood Approach for Comparing Synonymous and Nonsynonymous Nucleotide Substitution Rates, with Application to the Chloroplast Genome.” Molecular Biology and Evolution 11 (5): 715–24.

Ohta, T. 1989. “The Mutational Load of a Multigene Family with Uniform Members.” Genetical Research 53 (2): 141–45.

Olagunju, Temitayo A., Benjamin D. Rosen, Holly L. Neibergs, Gabrielle M. Becker, Kimberly M. Davenport, Christine G. Elsik, Tracy S. Hadfield, Sergey Koren, Kristen L. Kuhn, Arang Rhie, Katie A. Shira, Amy L. Skibiel, Morgan R. Stegemiller, Jacob W. Thorne, Patricia Villamediana, Noelle E. Cockett, Brenda M. Murdoch, and Timothy P. L. Smith. 2024. “Telomere-to-Telomere Assemblies of Cattle and Sheep Y-Chromosomes Uncover Divergent Structure and Gene Content.” Nature Communications 15 (1): 8277.

Pardo-Palacios, Francisco J., Angeles Arzalluz-Luque, Liudmyla Kondratova, Pedro Salguero, Jorge Mestre-Tomás, Rocío Amorín, Eva Estevan-Morió, Tianyuan Liu, Adalena Nanni, Lauren McIntyre, Elizabeth Tseng, and Ana Conesa. 2024. “SQANTI3: Curation of Long-Read Transcriptomes for Accurate Identification of Known and Novel Isoforms.” Nature Methods 21 (5): 793–97.

Patro, Rob, Geet Duggal, Michael I. Love, Rafael A. Irizarry, and Carl Kingsford. 2017. “Salmon Provides Fast and Bias-Aware Quantification of Transcript Expression.” Nature Methods 14 (4): 417–19.

Pertea, Geo, and Mihaela Pertea. 2020. “GFF Utilities: GffRead and GffCompare.” F1000Research 9 (304): 304.

Quinlan, Aaron R., and Ira M. Hall. 2010. “BEDTools: A Flexible Suite of Utilities for Comparing Genomic Features.” Bioinformatics (Oxford, England) 26 (6): 841–42.

Rhie, Arang, Sergey Nurk, Monika Cechova, Savannah J. Hoyt, Dylan J. Taylor, Nicolas Altemose, Paul W. Hook, Sergey Koren, Mikko Rautiainen, Ivan A. Alexandrov, Jamie Allen, Mobin Asri, Andrey V. Bzikadze, Nae-Chyun Chen, Chen-Shan Chin, Mark Diekhans, Paul Flicek, Giulio Formenti, Arkarachai Fungtammasan, Carlos Garcia Giron, Erik Garrison, Ariel Gershman, Jennifer L. Gerton, Patrick G. S. Grady, Andrea Guarracino, Leanne Haggerty, Reza Halabian, Nancy F. Hansen, Robert Harris, Gabrielle A. Hartley, William T. Harvey, Marina Haukness, Jakob Heinz, Thibaut Hourlier, Robert M. Hubley, Sarah E. Hunt, Stephen Hwang, Miten Jain, Rupesh K. Kesharwani, Alexandra P. Lewis, Heng Li, Glennis A. Logsdon, Julian K. Lucas, Wojciech Makalowski, Christopher Markovic, Fergal J. Martin, Ann M. Mc Cartney, Rajiv C. McCoy, Jennifer McDaniel, Brandy M. McNulty, Paul Medvedev, Alla Mikheenko, Katherine M. Munson, Terence D. Murphy, Hugh E. Olsen, Nathan D. Olson, Luis F. Paulin, David Porubsky, Tamara Potapova, Fedor Ryabov, Steven L. Salzberg, Michael E. G. Sauria, Fritz J. Sedlazeck, Kishwar Shafin, Valery A. Shepelev, Alaina Shumate, Jessica M. Storer, Likhitha Surapaneni, Angela M. Taravella Oill, Françoise Thibaud-Nissen, Winston Timp, Marta Tomaszkiewicz, Mitchell R. Vollger, Brian P. Walenz, Allison C. Watwood, Matthias H. Weissensteiner, Aaron M. Wenger, Melissa A. Wilson, Samantha Zarate, Yiming Zhu, Justin M. Zook, Evan E. Eichler, Rachel J. O’Neill, Michael C. Schatz, Karen H. Miga, Kateryna D. Makova, and Adam M. Phillippy. 2023. “The Complete Sequence of a Human Y Chromosome.” Nature 621 (7978): 344–54.

Rice, P., I. Longden, and A. Bleasby. 2000. “EMBOSS: The European Molecular Biology Open Software Suite.” Trends in Genetics: TIG 16 (6): 276–77.

Rombel, Irene T., Kathryn F. Sykes, Simon Rayner, and Stephen Albert Johnston. 2002. “ORF-FINDER: A Vector for High-Throughput Gene Identification.” Gene 282 (1-2): 33–41.

Rozen, Steve, Helen Skaletsky, Janet D. Marszalek, Patrick J. Minx, Holland S. Cordum, Robert H. Waterston, Richard K. Wilson, and David C. Page. 2003. “Abundant Gene Conversion between Arms of Palindromes in Human and Ape Y Chromosomes.” Nature 423 (6942): 873–76.

Sahlin, Kristoffer, and Veli Mäkinen. 2021. “Accurate Spliced Alignment of Long RNA Sequencing Reads.” Bioinformatics (Oxford, England) 37 (24): 4643–51.

Sahlin, Kristoffer, Marta Tomaszkiewicz, Kateryna D. Makova, and Paul Medvedev. 2018. “Deciphering Highly Similar Multigene Family Transcripts from Iso-Seq Data with IsoCon.” Nature Communications 9 (1): 4601.

Sakamoto, Takahiro, and Hideki Innan. 2022. “Muller’s Ratchet of the Y Chromosome with Gene Conversion.” Genetics 220 (1). 10.1093/genetics/iyab204.

Shi, Wentao, Sandra Louzada, Marina Grigorova, Andrea Massaia, Elena Arciero, Laura Kibena, Xiangyu Jack Ge, Yuan Chen, Qasim Ayub, Olev Poolamets, Chris Tyler-Smith, Margus Punab, Maris Laan, Fengtang Yang, Pille Hallast, and Yali Xue. 2019. “Evolutionary and Functional Analysis of RBMY1 Gene Copy Number Variation on the Human Y Chromosome.” Human Molecular Genetics 28 (16): 2785–98.

Shi, Wentao, Andrea Massaia, Sandra Louzada, Juliet Handsaker, William Chow, Shane McCarthy, Joanna Collins, Pille Hallast, Kerstin Howe, Deanna M. Church, Fengtang Yang, Yali Xue, and Chris Tyler-Smith. 2019. “Birth, Expansion, and Death of VCY-Containing Palindromes on the Human Y Chromosome.” Genome Biology 20 (1): 207.

Shumate, Alaina, Brandon Wong, Geo Pertea, and Mihaela Pertea. 2022. “Improved Transcriptome Assembly Using a Hybrid of Long and Short Reads with StringTie.” PLoS Computational Biology 18 (6): e1009730.

Sim, Ngak-Leng, Prateek Kumar, Jing Hu, Steven Henikoff, Georg Schneider, and Pauline C. Ng. 2012. “SIFT Web Server: Predicting Effects of Amino Acid Substitutions on Proteins.” Nucleic Acids Research 40 (Web Server issue): W452–57.

Skaletsky, Helen, Tomoko Kuroda-Kawaguchi, Patrick J. Minx, Holland S. Cordum, Ladeana Hillier, Laura G. Brown, Sjoerd Repping, Tatyana Pyntikova, Johar Ali, Tamberlyn Bieri, Asif Chinwalla, Andrew Delehaunty, Kim Delehaunty, Hui Du, Ginger Fewell, Lucinda Fulton, Robert Fulton, Tina Graves, Shun-Fang Hou, Philip Latrielle, Shawn Leonard, Elaine Mardis, Rachel Maupin, John McPherson, Tracie Miner, William Nash, Christine Nguyen, Philip Ozersky, Kymberlie Pepin, Susan Rock, Tracy Rohlfing, Kelsi Scott, Brian Schultz, Cindy Strong, Aye Tin-Wollam, Shiaw-Pyng Yang, Robert H. Waterston, Richard K. Wilson, Steve Rozen, and David C. Page. 2003. “The Male-Specific Region of the Human Y Chromosome Is a Mosaic of Discrete Sequence Classes.” Nature 423 (6942): 825–37.

Smit, A. F. A., R. Hubley, and P. Green. 2021. “2013–2015. RepeatMasker Open-4.0.” https://scholar.google.com/citations?user=SOqGFioAAAAJ&hl=en&oi=sra.

Smith, G. P. 1976. “Evolution of Repeated DNA Sequences by Unequal Crossover.” Science (New York, N.Y.) 191 (4227): 528–35.

Soh, Y. Q. Shirleen, Jessica Alföldi, Tatyana Pyntikova, Laura G. Brown, Tina Graves, Patrick J. Minx, Robert S. Fulton, Colin Kremitzki, Natalia Koutseva, Jacob L. Mueller, Steve Rozen, Jennifer F. Hughes, Elaine Owens, James E. Womack, William J. Murphy, Qing Cao, Pieter de Jong, Wesley C. Warren, Richard K. Wilson, Helen Skaletsky, and David C. Page. 2014. “Sequencing the Mouse Y Chromosome Reveals Convergent Gene Acquisition and Amplification on Both Sex Chromosomes.” Cell 159 (4): 800–813.

Stouffs Katrien, Willy Lissens, Greta Verheyen, Lisbet Van Landuyt, Annieta Goossens, Herman Tournaye, André Van Steirteghem, and Inge Liebaers. 2004. “Expression Pattern of the Y-Linked PRY Gene Suggests a Function in Apoptosis but Not in Spermatogenesis.” Molecular Human Reproduction 10 (1): 15–21.

Stouffs, K., W. Lissens, L. Van Landuyt, H. Tournaye, A. Van Steirteghem, and I. Liebaers. 2001. “Characterization of the Genomic Organization, Localization and Expression of Four PRY Genes (PRY1, PRY2, PRY3 and PRY4).” Molecular Human Reproduction 7 (7): 603–10.

Swanepoel, Callie M., and Jacob L. Mueller. 2024. “Out with the Old, in with the New: Meiotic Driving of Sex Chromosome Evolution.” Seminars in Cell & Developmental Biology 163 (November):14–21.

Swanepoel, Callie M., Gaojianyong Wang, Lucy Zhang, Björn Brändl, Hermann Bauer, Pavel Tsaytler, Franz-Josef Müller, Bernhard G. Herrmann, and Jacob L. Mueller. 2025. “Acquisition of Ampliconic Sequences Marks a Selfish Mouse T-Haplotype.” bioRxivorg. 10.1101/2025.01.28.635315.

Taguchi, Ayumu, Allen D. Taylor, Jaime Rodriguez, Müge Celiktaş, Hui Liu, Xiaotu Ma, Qing Zhang, Chee-Hong Wong, Alice Chin, Luc Girard, Carmen Behrens, Wan L. Lam, Stephen Lam, John D. Minna, Ignacio I. Wistuba, Adi F. Gazdar, and Samir M. Hanash. 2014. “A Search for Novel Cancer/testis Antigens in Lung Cancer Identifies VCX/Y Genes, Expanding the Repertoire of Potential Immunotherapeutic Targets.” Cancer Research 74 (17): 4694–4705.

Tomaszkiewicz, Marta, Kristoffer Sahlin, Paul Medvedev, and Kateryna D. Makova. 2023. “Transcript Isoform Diversity of Ampliconic Genes on the Y Chromosome of Great Apes.” Genome Biology and Evolution 15 (11). 10.1093/gbe/evad205.

Trombetta, Beniamino, and Fulvio Cruciani. 2017. “Y Chromosome Palindromes and Gene Conversion.” Human Genetics 136 (5): 605–19.

Tse, J. Y. M., E. Y. M. Wong, A. N. Y. Cheung, W. S. O P. C. Tam, and W. S. B. Yeung. 2003. “Specific Expression of VCY2 in Human Male Germ Cells and Its Involvement in the Pathogenesis of Male Infertility.” Biology of Reproduction 69 (3): 746–51.

Vaser, Robert, Ivan Sović, Niranjan Nagarajan, and Mile Šikić. 2017. “Fast and Accurate de Novo Genome Assembly from Long Uncorrected Reads.” Genome Research 27 (5): 737–46.

Vegesna, Rahulsimham, Marta Tomaszkiewicz, Paul Medvedev, and Kateryna D. Makova. 2019. “Dosage Regulation, and Variation in Gene Expression and Copy Number of Human Y Chromosome Ampliconic Genes.” PLoS Genetics 15 (9): e1008369.

Vegesna, Rahulsimham, Marta Tomaszkiewicz, Oliver A. Ryder, Rebeca Campos-Sánchez, Paul Medvedev, Michael DeGiorgio, and Kateryna D. Makova. 2020. “Ampliconic Genes on the Great Ape Y Chromosomes: Rapid Evolution of Copy Number but Conservation of Expression Levels.” Genome Biology and Evolution 12 (6): 842–59.

Veyrunes, Frédéric, Paul D. Waters, Pat Miethke, Willem Rens, Daniel McMillan, Amber E. Alsop, Frank Grützner, Janine E. Deakin, Camilla M. Whittington, Kyriena Schatzkamer, Colin L. Kremitzki, Tina Graves, Malcolm A. Ferguson-Smith, Wes Warren, and Jennifer A. Marshall Graves. 2008. “Bird-like Sex Chromosomes of Platypus Imply Recent Origin of Mammal Sex Chromosomes.” Genome Research 18 (6): 965–73.

Wong, Elaine Y. M., Jenny Y. M. Tse, Kwok-Ming Yao, Po-Chor Tam, and William S. B. Yeung. 2002. “VCY2 Protein Interacts with the HECT Domain of Ubiquitin-Protein Ligase E3A.” Biochemical and Biophysical Research Communications 296 (5): 1104–11.

Xue, Yali, and Chris Tyler-Smith. 2011. “An Exceptional Gene: Evolution of the TSPY Gene Family in Humans and Other Great Apes.” Genes 2 (1): 36–47.

Ye, Danling, Arslan A. Zaidi, Marta Tomaszkiewicz, Kate Anthony, Corey Liebowitz, Michael DeGiorgio, Mark D. Shriver, and Kateryna D. Makova. 2018. “High Levels of Copy Number Variation of Ampliconic Genes across Major Human Y Haplogroups.” Genome Biology and Evolution 10 (5): 1333–50.

Yoo, Dongahn, Arang Rhie, Prajna Hebbar, Francesca Antonacci, Glennis A. Logsdon, Steven J. Solar, Dmitry Antipov, Brandon D. Pickett, Yana Safonova, Francesco Montinaro, Yanting Luo, Joanna Malukiewicz, Jessica M. Storer, Jiadong Lin, Abigail N. Sequeira, Riley J. Mangan, Glenn Hickey, Graciela Monfort Anez, Parithi Balachandran, Anton Bankevich, Christine R. Beck, Arjun Biddanda, Matthew Borchers, Gerard G. Bouffard, Emry Brannan, Shelise Y. Brooks, Lucia Carbone, Laura Carrel, Agnes P. Chan, Juyun Crawford, Mark Diekhans, Eric Engelbrecht, Cedric Feschotte, Giulio Formenti, Gage H. Garcia, Luciana de Gennaro, David Gilbert, Richard E. Green, Andrea Guarracino, Ishaan Gupta, Diana Haddad, Junmin Han, Robert S. Harris, Gabrielle A. Hartley, William T. Harvey, Michael Hiller, Kendra Hoekzema, Marlys L. Houck, Hyeonsoo Jeong, Kaivan Kamali, Manolis Kellis, Bryce Kille, Chul Lee, Youngho Lee, William Lees, Alexandra P. Lewis, Qiuhui Li, Mark Loftus, Yong Hwee Eddie Loh, Hailey Loucks, Jian Ma, Yafei Mao, Juan F. I. Martinez, Patrick Masterson, Rajiv C. McCoy, Barbara McGrath, Sean McKinney, Britta S. Meyer, Karen H. Miga, Saswat K. Mohanty, Katherine M. Munson, Karol Pal, Matt Pennell, Pavel A. Pevzner, David Porubsky, Tamara Potapova, Francisca R. Ringeling, Joana L. Rocha, Oliver A. Ryder, Samuel Sacco, Swati Saha, Takayo Sasaki, Michael C. Schatz, Nicholas J. Schork, Cole Shanks, Linnéa Smeds, Dongmin R. Son, Cynthia Steiner, Alexander P. Sweeten, Michael G. Tassia, Françoise Thibaud-Nissen, Edmundo Torres-González, Mihir Trivedi, Wenjie Wei, Julie Wertz, Muyu Yang, Panpan Zhang, Shilong Zhang, Yang Zhang, Zhenmiao Zhang, Sarah A. Zhao, Yixin Zhu, Erich D. Jarvis, Jennifer L. Gerton, Iker Rivas-González, Benedict Paten, Zachary A. Szpiech, Christian D. Huber, Tobias L. Lenz, Miriam K. Konkel, Soojin V. Yi, Stefan Canzar, Corey T. Watson, Peter H. Sudmant, Erin Molloy, Erik Garrison, Craig B. Lowe, Mario Ventura, Rachel J. O’Neill, Sergey Koren, Kateryna D. Makova, Adam M. Phillippy, and Evan E. Eichler. 2025. “Complete Sequencing of Ape Genomes.” Nature 641 (8062): 401–18.

Zhou, Ran, David Macaya-Sanz, Craig H. Carlson, Jeremy Schmutz, Jerry W. Jenkins, David Kudrna, Aditi Sharma, Laura Sandor, Shengqiang Shu, Kerrie Barry, Gerald A. Tuskan, Tao Ma, Jianquan Liu, Matthew Olson, Lawrence B. Smart, and Stephen P. DiFazio. 2020. “A Willow Sex Chromosome Reveals Convergent Evolution of Complex Palindromic Repeats.” Genome Biology 21 (1): 38.

Zhu, Zexian, Lubna Younas, and Qi Zhou. 2025. “Evolution and Regulation of Animal Sex Chromosomes.” Nature Reviews. Genetics 26 (1): 59–74.

Zou, Sheng Wei, Jian Chao Zhang, Xiao Dong Zhang, Shi Ying Miao, Shu Dong Zong, Qi Sheng, and Lin Fang Wang. 2003. “Expression and Localization of VCX/Y Proteins and Their Possible Involvement in Regulation of Ribosome Assembly during Spermatogenesis.” Cell Research 13 (3): 171–77.

