## Supplemental Information for "How and why multicopy genes survive on the human Y chromosome"

### Supplementary Information

### Supplementary Notes

#### Note S1. Autosomal homologs and X gametologs of YAGs

While determining the complete repertoire of YAGs, we also identified YAG homologs on other chromosomes (Table S20). We confirmed the X-chromosome homologs of *RBMY* and *VCY (Bhowmick, Satta, and Takahata 2007; Vallender and Lahn 2004)*. These two gene families displayed the opposite expansion patterns: *RBMX*/*RBMY* expanded mostly on the Y chromosome, whereas *VCX*/*VCY* expanded mostly on the X chromosome (Table S20). Additionally, we identified several autosomal *RBMY* homologs: the more closely related genes *RBMXL1-RBMXL3,* which originated by retrotransposition of *RBMX* to autosomes (Lingenfelter et al. 2001; Elliott et al. 2019), and more distantly related genes (*RBMR3*, *CIRBP*, *TRA2A/B*, and *MSI1/2*; Figure S18), which share a highly conserved RNA-binding domain. Consistent with prior literature (Lahn and Page 1999; Saxena et al. 1996), we identified two autosomal paralogs for each of the *CDY* and *DAZ* gene families—*CDYL* and *CDYL2*, and *DAZL* and *BOL*, respectively. No autosomal or X chromosome protein-coding homologs were identified for *BPY*2, *HSFY*, and *TSPY*. *BPY2* (also called *VCY*2) was suggested to be derived from a non-protein-coding gene that gained protein-coding function (Cao et al. 2015). Both *HSFY* and *TSPY* have homologs on the X chromosome according to prior studies (Bhowmick, Satta, and Takahata 2007; Delbridge et al. 2004); however, we could not detect them because their X-Y divergence occurred prior to or during early mammalian radiation (Bhowmick, Satta, and Takahata 2007; Delbridge et al. 2004; Hughes et al. 2020), resulting in sequence identity below our detection threshold of 35% similarity. This is shown in Additional Data File 5 sheet 6; pairwise identity between the *HSFY*, *HSFX*1/2, and *HSFX*3/4 gene families forming three separate blocks with pairwise identity <30%, and in Additional Data File 5 sheet 7, where we observe two separate clusters formed by the *TSPY* gene family and the *TSPYL2* (sometimes referred to as *TSPX*) paralogs, with pairwise identity <35%. In summary, our analysis of T2T ape genomes confirms the transposition origin of *CDY* and *DAZ,* and the proto-sex-chromosome origin of *RBMY* and *VCY.*

### Note S2. A detailed description of the organization and phylogenetic patterns for YAG families

To investigate duplication history and gene conversion–driven homogenization within YAG families, we analyzed phylogenetic relationships (using the maximum likelihood approach, see Methods) among paralogs and orthologs within each YAG family (**Figure 2C**). This approach has not yet been applied to T2T genomes, except for the *TSPY* gene family, where phylogeny was still not linked to array versus palindrome organization (Makova et al. 2024). For *BPY2*, nearly identical palindromic gene pairs clustered tightly within species, with chimpanzee retaining only a single copy. *CDY* copies largely formed species-specific clades, though complex repeat structures in orangutans indicated duplications predating speciation with ongoing palindrome-mediated homogenization. Two separate phylogenetic trees were built for the two domains of the *DAZ* gene (**Figure 2C** and Figure S3). *DAZ* showed striking domain-specific patterns: RNA-recognition motifs (RRMs) were highly homogenized and clustered by genus or species (**Figure 2C**), whereas 72- or 90-bp exon repeats (*DAZ* repeats) evolved faster, were less homogenized, and showed weaker phylogenetic structure (Figure S3). *HSFY* copies were homogenized within siamang, human, and gorilla but remained intermingled between orangutan species, reflecting insufficient divergence time for homogenization. *RBMY* exhibited extensive within-species homogenization across palindromes and tandem arrays, with orangutans retaining two distinct gene variants from ancient duplication events. *TSPY*, one of the most abundant Y gene families, was highly homogenized within species despite variable organization across palindromes and arrays. Finally, *VCY* showed evidence of X–Y gene conversion (consistent with (Iwase et al. 2010)), as Y-linked copies did not form a distinct clade from their X-linked homologs. Together, these results highlight palindromes and tandem arrays as powerful drivers of Y-chromosome gene homogenization, modulated by lineage-specific evolutionary histories.. Detailed description of individual gene families follows:

***BPY2.*** The *BPY2* gene family is present only in Homininae (bonobo, chimpanzee, human, and gorilla). Thus, we rooted its phylogenetic tree with the gorilla gene copies. We discovered that, in each species except for chimpanzee, *BPY2* gene copies are present in pairs on the opposite arms of palindromes. Such pairs are identical (Table S21) and group together on the phylogenetic tree. Chimpanzee has only one *BPY2* gene copy. In humans, in addition to the two copies located on the opposite arms of palindrome P2, there is also a third copy located outside of palindromes or arrays. All three human gene copies have high sequence identity and cluster together on the phylogenetic tree.

***CDY***. *CDY* gene copies are mostly located within palindromes. On the phylogenetic tree, which we rooted with the *CDYL*—the closest autosomal homolog of *CDY (Dorus et al. 2003)*—gene copies form species-specific clades (except in orangutans). Within these clades, most gene copy pairs located on opposite arms of the same palindrome cluster together. *CDY* gene copies in orangutans are numerous (13 in Bornean and 22 in Sumatran) and are located in a highly complex region, containing multiple levels of repeats and inversions (Figure S19 A, B, C) and also harboring *DAZ* and *HSFY* copies. On the phylogenetic tree, the *CDY* gene copies form several clades, including sequences from both orangutan species, suggesting duplications that predated their speciation. However, within such clades, gene copies from the same species located on opposite arms of the same palindrome cluster together, suggesting palindrome-aided sequence homogenization. All four gorilla gene copies have high sequence identity and form a cluster on the phylogenetic tree; three of the copies were newly added as compared to (Makova et al. 2024), and are located within a complex double-palindrome structure with a potential for intrachromosomal sequence homogenization.

***DAZ***. In the T2T ape genomes (Makova et al. 2024), *DAZ* gene copies—all located within palindromes—contain up to three 495-bp RNA-recognition motifs (‘RRMs’) and up to 16 72- or 90-bp repeats

(‘*DAZ* repeats’) (Saxena et al. 1996) (Table S22). Therefore, two separate phylogenetic trees were constructed—for the RRMs (including each RRM for each gene copy separately) and for *DAZ* repeats (including each *DAZ* repeat for each gene copy separately). The *DAZ* RRM tree was rooted with *DAZL*, the closest autosomal homolog of *DAZ* (Xu, Moore, and Pera 2001; Hughes, Skaletsky, and Page 2012). Since the *DAZL* repeats did not form a monophyletic clade, the *DAZ* repeats tree was unrooted.

The RRM tree has species-specific clades for human, gorilla, and siamang, and genus-specific clades for *Pongo* and *Pan*. All human RRMs have an identical DNA sequence, suggesting homogenization via gene conversion. In gorilla, domain-copy-specific clades were observed (i.e., for RRMI and for RRMII). In chimpanzee and bonobo, all RRM domain copies formed a single monophyletic clade with highly similar sequences across species. Within the larger monophyletic clade for orangutans, RRM copies usually cluster depending on their position within the *DAZ* gene, creating the RRMI, RRMII, and RRMIII groups, likely due to palindrome-aided sequence homogenization. This is clearly demonstrated between *DAZ* copies in Bornean orangutan (Figure S20), where we observe 100% sequence identity between RRMs directly across each other on opposing palindrome arms, while RRM duplications within the same arm show some divergence (with sequence identity of 96.8-98.4%). Interestingly, *DAZ* is the only gene consistently found in palindromic configuration within the complex region on the Y in orangutans (Figure S19).

In contrast, the phylogenetic tree of *DAZ* repeats does not cluster by species, and generally has lower bootstrap values and higher overall evolutionary distance than those on the RRM tree, suggesting less homogenization (Figure S3). We could still observe some *Pongo*-specific and Homininae-specific clades, as well as several clades with two repeats located on the opposite arms of the same palindrome. In summary, we observe two very different phylogenetic trees constructed from different parts of the same gene. The RRM domain shows clear signs of homogenization and forms distinct genus- or species-specific clades, whereas *DAZ* repeats appear to be shuffled, evolve more rapidly, and are less homogenized.

***HSFY***. *HSFY* gene copies are located within palindromes in the siamang, human, and gorilla (except for one copy). In orangutans, they are located in a complex repetitive region with homologous sequences that cannot be classified into simple palindromes or repeat arrays (this region also harbors the *CDY* and *DAZ* genes; Figure S19). Our examination of the *HSFY* phylogenetic tree, rooted with siamang gene copies, revealed the following patterns. In siamang, human, and gorilla, the gene copies are very similar within each species and cluster into species-specific groups. This indicates that enough time has passed since human and gorilla diverged for gene copies to become homogenized within each species. In contrast, orangutan gene copies do not separate by species and instead form a single mixed group, suggesting that their divergence was too recent for homogenization to fully shape the *HSFY* phylogenetic pattern.

***RBMY***. Copies of the *RBMY* gene are located in palindromes (in bonobo, chimpanzee, human, and siamang) and in both palindromes and tandemly repeated gene arrays (in gorilla and the orangutans). In human, all copies are located on Inverted Repeat 2 (IR2) and palindrome P3 (Skaletsky et al. 2003), and form a cluster on the phylogenetic tree with additional groupings corresponding to gene copy pairs located on the opposite arms of the same palindrome. In both bonobo and chimpanzee, similar inverted repetitive structures are observed, with additional amplification in bonobo (Figure S21). We observed a chimpanzee-specific clade and two bonobo-specific clades on the phylogenetic tree (Figure S22A). The two bonobo clades were likely driven by distinct nucleotide differences (Figure S22B), possibly originating from two distant paralogs, however, gene copies originating from the two clades are not spatially separated (Figure S22C), suggesting homogenization due to location in palindromes and tandem repeats. Siamang copies are located on palindrome Q3, where arm B contains an internal duplication creating an unbalanced palindrome. Siamang copies have high sequence identity and form a separate cluster on the phylogenetic tree. In gorilla, the palindromic and array gene copies create a monophyletic cluster on the phylogenetic tree. Orangutans have two distinct variants of the *RBMY* gene. The first variant comprises four identical copies (two in each species), is located within palindromes, and groups with copies from the other species. The second variant, previously identified in (Makova et al. 2024), is organized in arrays and forms a separate cluster outside of the rest of the tree, suggesting an early duplication event. Gene copies within this array are highly similar to each other, suggesting extensive homogenization. Overall, the *RBMY* tree appears highly homogenized within each species, although orangutan gene copies differ between the two gene copy variants. Two gorilla and one chimpanzee copy group outside of species-specific clusters because their sequences are truncated.

***TSPY***. The organization and phylogenetic patterns of the *TSPY* gene family were described in detail in (Makova et al. 2024). Since then, we have identified nine additional *TSPY* copies in bonobo and five in siamang. Moreover, our gene copies (one copy in each of bonobo, chimpanzee, Sumatran orangutan, and Bornean orangutan) previously considered protein-coding are now considered pseudogenes. *TSPY* is among the most copious gene families in apes, yet copies are highly homogenized within species. In most species (chimpanzee, human, gorilla, and both orangutans), copies of this gene are organized in tandem arrays. In siamang, all copies of the gene are located within palindromes. In bonobo, some copies are present in palindromes, others are present in tandem arrays, and yet others are present in arrays located on palindromes shared with *RBMY* and can form unbalanced repeats (occurring only on one arm of a palindrome) resembling small tandem arrays (Figure S23). On the phylogenetic tree we built using siamang Y copies as an outgroup, we observed species-specific clades for human and gorilla, and genus-specific clades for *Pan* and *Pongo*. We observed high levels of sequence homogenization among gene copies within the same tandem array.

***VCY***. Finally, the *VCY* gene family is present only in the siamang, where a single copy was detected, and in Hominini (human, bonobo, and chimpanzee), where two palindromic copies are present in each species. It was likely lost from the Y in gorilla and orangutans. Interestingly, the X-linked homolog of *VCY*—*VCX*—is present in multiple copies across all studied species (ranging from four in gorilla to 16 in bonobo). The *VCX* copies are located within palindromes in all species except the two orangutans. Notably, on the phylogenetic trees, the *VCY* gene copies are interspersed among the *VCX* gene copies and do not form a separate phylogeny, consistent with gene conversion between the X and Y paralogs, as was suggested previously (Bhowmick, Satta, and Takahata 2007).

### Note S3. Long-read reference transcriptome construction and affect RNA-seq expression estimates

**Comparison of `STRINGTIE`-produced structural isoforms with previous non-reference results.** For this comparison, we used replicate-supported transcripts (present in at least two of the three transcript sets, see section "Transcriptome assembly and quality control") from Table S23. Our results identified only 445 replicate-supported transcripts, whereas Tomaszkiewicz et al. (2023) identified 1,510. On average, transcripts identified by the *de novo* assembly approach were more than two times shorter (mean length of  820 bp, Tomaszkiewicz et al., 2023), as compared to those in our reference-based approach (mean length = 1,893).

**Transcript expression levels.** To evaluate the effect of our reference-augmented transcriptome on capturing YAG expression in great apes, we mapped testis short-read RNA-seq data to it and (separately) to the standard NCBI RefSeq coding sequence (CDS) set. Expression estimates were compared at the YAG family level (separately for each species) to assess whether the reference-augmented transcriptome recovered additional signal (Table S10). Most gene families were unaffected by the inclusion of novel transcripts. However, in some cases, the number of mapped reads increased, most notably for *DAZ* in human and Bornean orangutan. In other cases, TPM  (Transcripts Per Million) values changed despite identical read counts, as observed for *TSPY* in Bornean orangutan (Figure S7; Table S10). *VCY* was the most highly expressed family in chimpanzee and bonobo, *HSFY* dominated in human and Sumatran orangutan, and *TSPY* showed the highest expression in gorilla and Bornean orangutan. Overall, these results suggest that expression estimates are sensitive to transcriptome choice and that accurately modeling YAG transcript structures is important for interpreting RNA-seq data.

### Note S4. Examination of branch-specific selection

When analyzing the complete gene sequences for each family, we observed significant variability in selection pressure (i.e., significant branch-heterogeneity test results) across branches for CDY and RBMY. However, no individual branch displayed statistically significant evidence of positive selection for CDY, and only one such branch was detected (with aBSREL analysis (Smith et al. 2015)) in RBMY (it led to one gene copy in gorilla; Figure S24)

### Supplementary Figures

#### Figure S1. The relationship between the mean and the standard deviation of copy number for YAG families

Correlation between the mean and standard deviation of copy number for each gene family in the three species (gorilla, human, and chimpanzee), for which the data from multiple individuals were available.

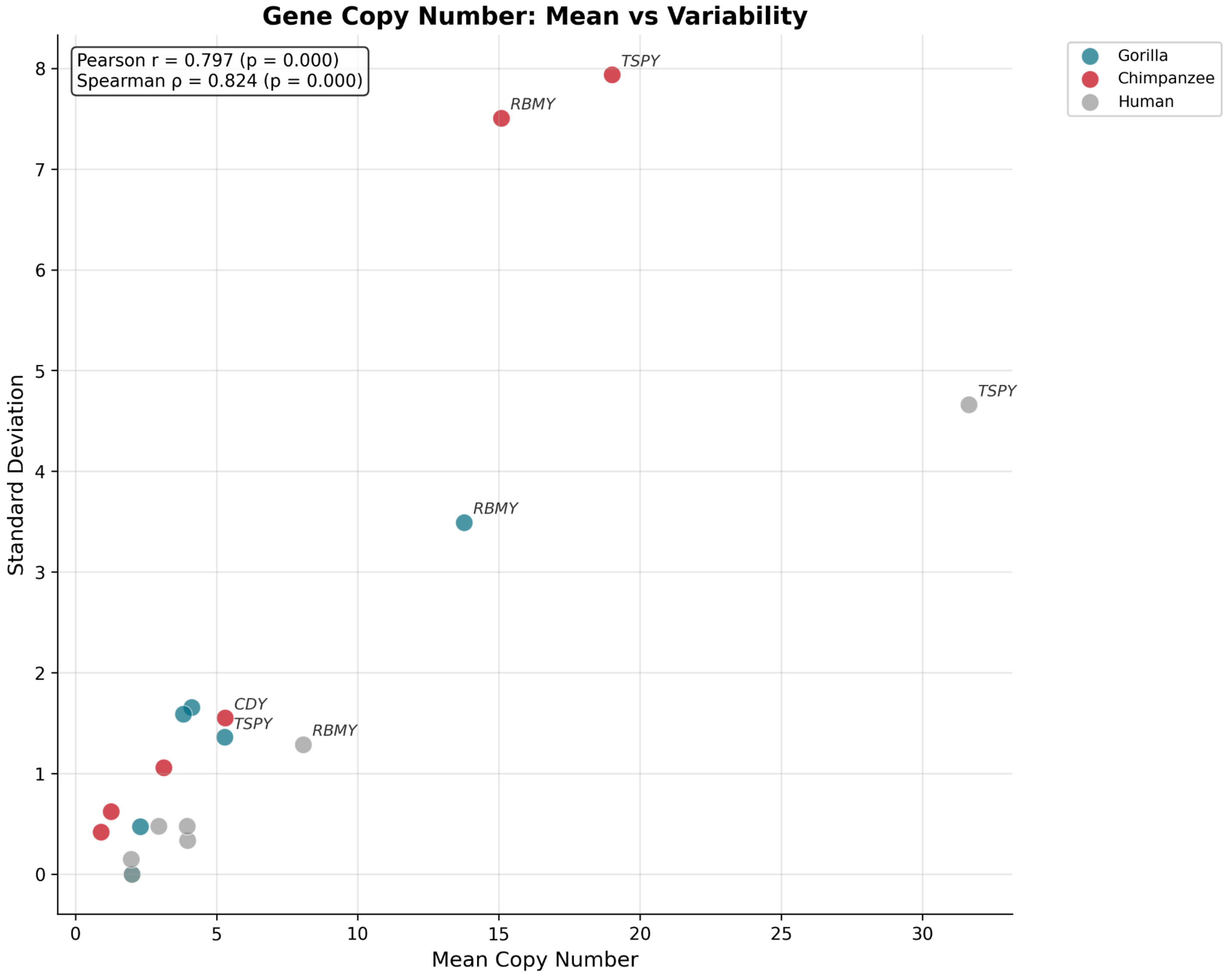

Figure S2. A correlation between pairwise sequence identity and pairwise distance between gene copies belonging to the same YAG family in the same species, for gene copies located in tandemly repeated arrays

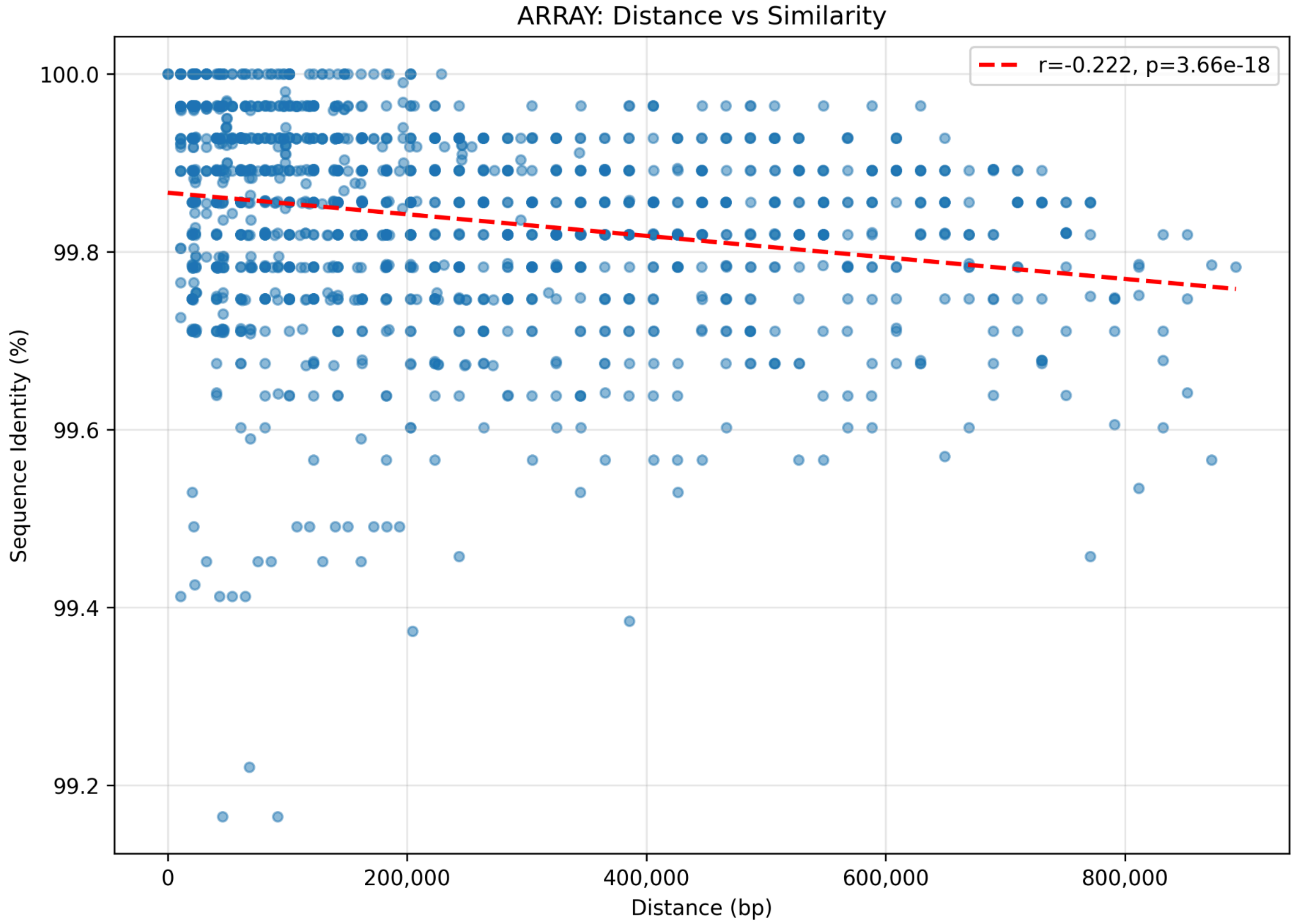

### Figure S3. The phylogenetic relationship among *DAZ* 72- or 90-bp repeats

Autosomal copies (highlighted in a circle) do not form a distinct clade, hence the phylogenetic tree is unrooted.

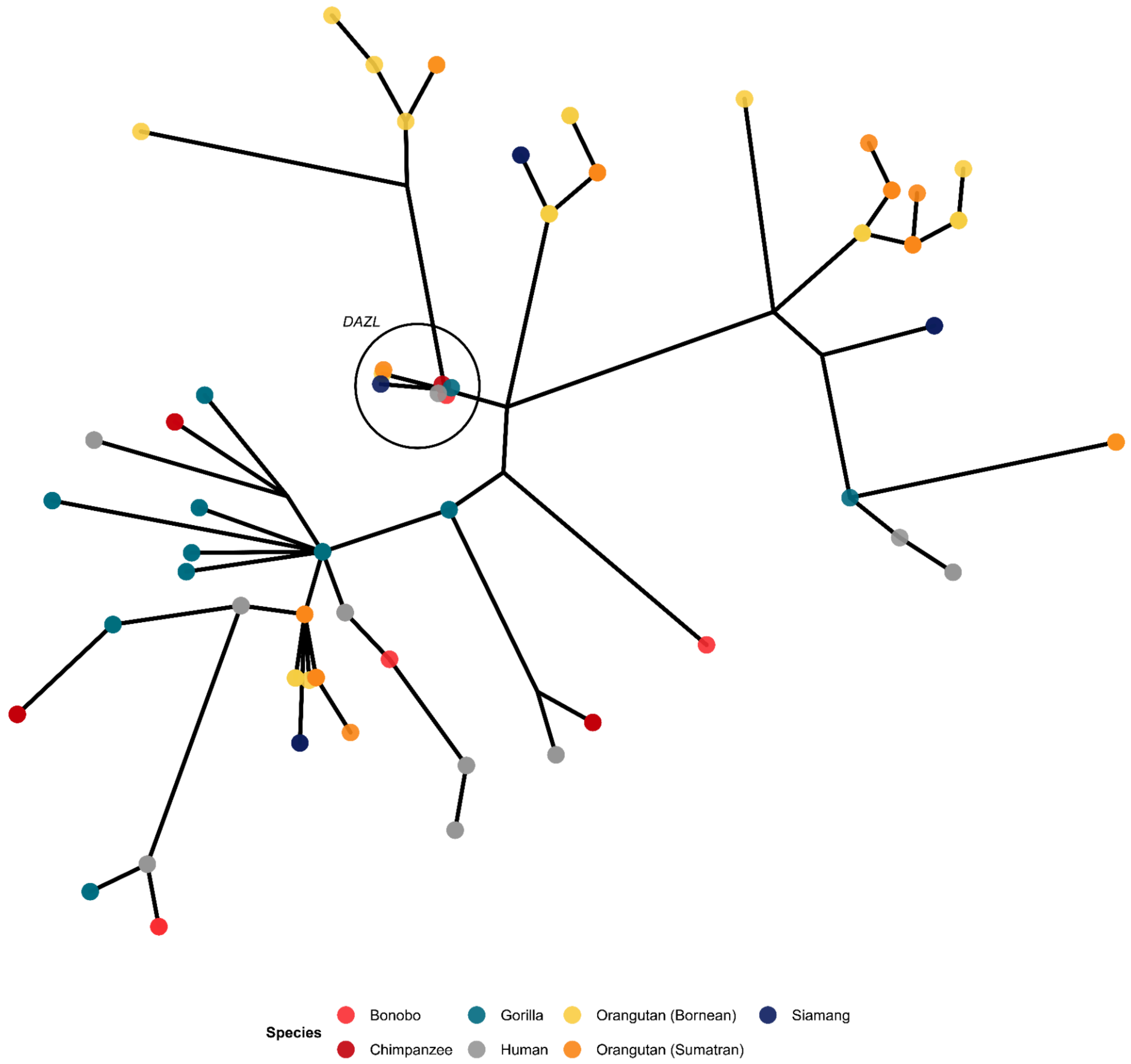

### Figure S4. Structural and sequence transcript isoform diversity of YAGs

(**A**) Correlation between the number of non-overlapping exons (in the longest human reference gene) and the number of observed **structural** isoforms. Each combination of YAG families and species was considered separately. The dataset excluded the *DAZ* gene, which contains complex internal duplications, and was analyzed using a modified approach that omitted the alignment step used to compare sequence isoforms. (**B**) Correlation between the number of gene copies (counts in reference genomes for bonobo, and Sumatran and Bornean orangutans, averages from multiple individuals for human, gorilla, and chimpanzee) and the number of **sequence** isoforms.

A

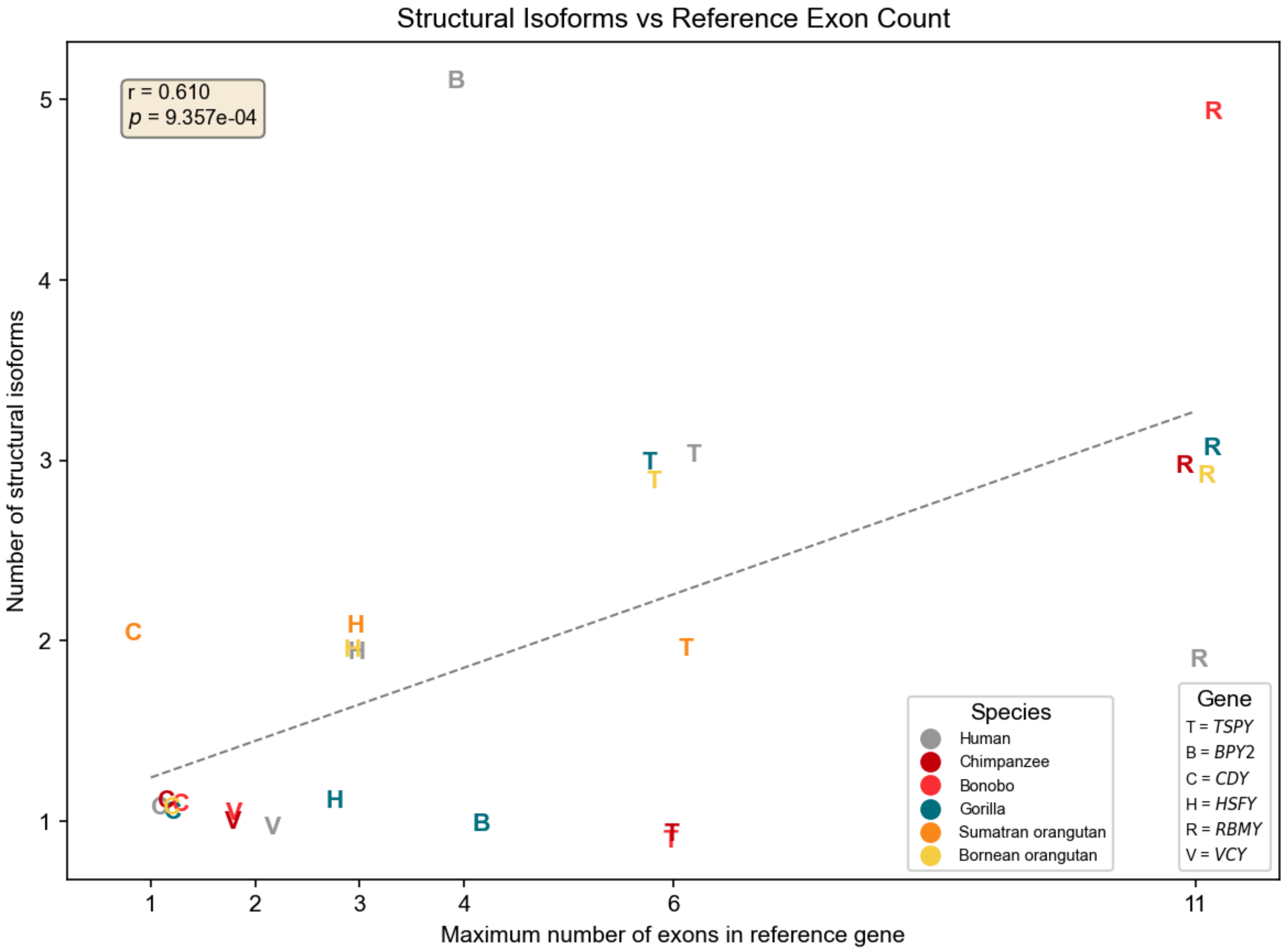

B

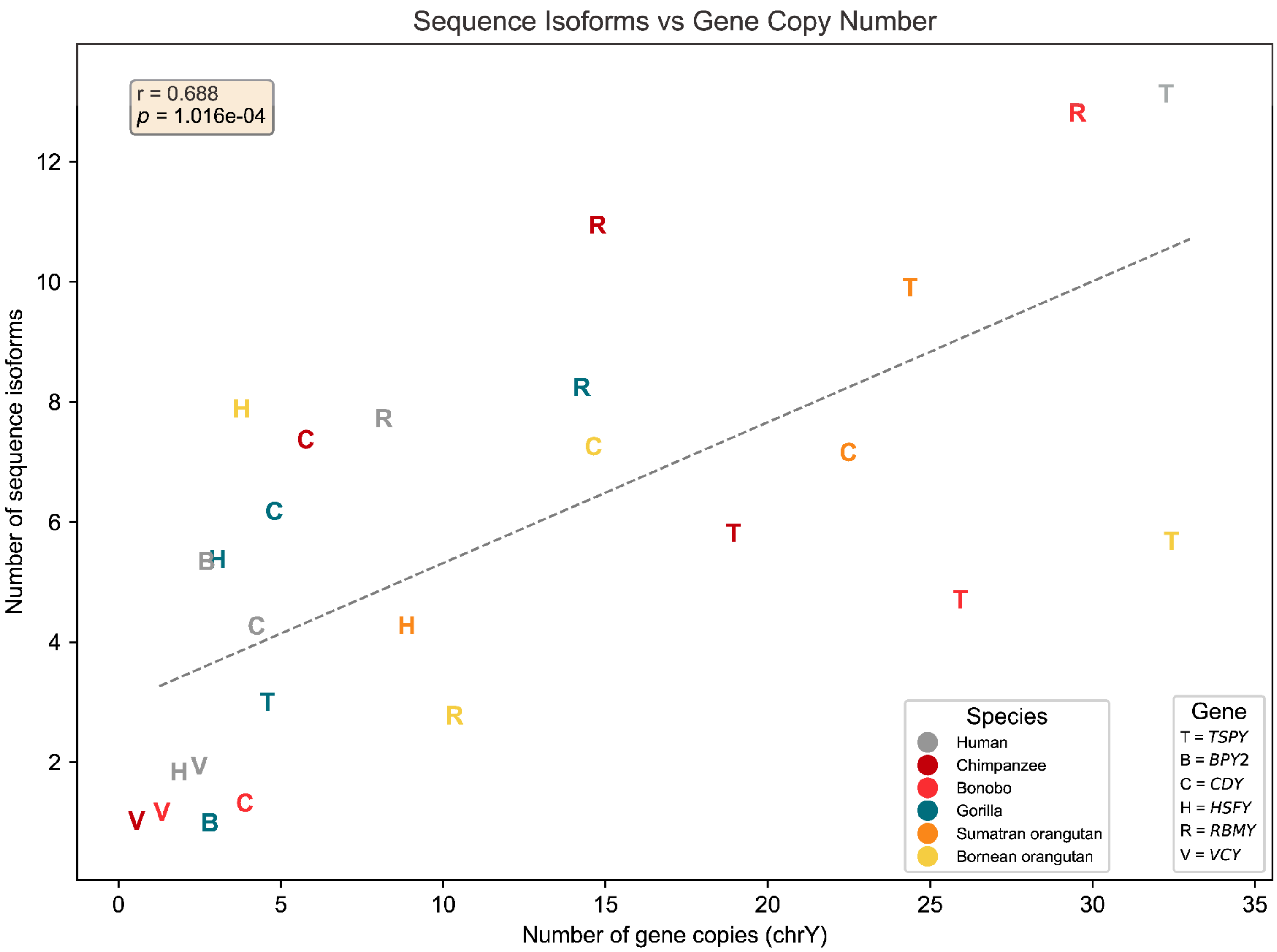

Sequence Isoforms vs Gene Copy Number
r = 0.688
p = 1.016e-04
Number of sequence isoforms
Number of gene copies (chrY)
Species
Human
Chimpanzee
Bonobo
Gorilla
Sumatran orangutan
Bornean orangutan
Gene
T = TSPY
B = BPY2
C = CDY
H = HSFY
R = RBMY
V = VCY

### Figure S5. Alternative splicing of *RBMY* in the *Pongo* lineage

(**A**) Alternative splicing of *RBMY* in the *Pongo* lineage due to the deletion of one copy of the 111-bp repeated exon. The plot shows DNA sequences (gray line with black highlights for mismatches and thin black line for deletions). Yellow boxes represent annotated exons. An inset zoom-in on the sequence at the nucleotide resolution shows the absence of a splice site. (**B**) Pairwise similarity between genomic regions. With each exon, the following ~440 intron region was used. The deleted region shows high homology with repeat A, while the newly formed *Pongo*-specific exon shows high homology with the intronic counterpart in human.

A

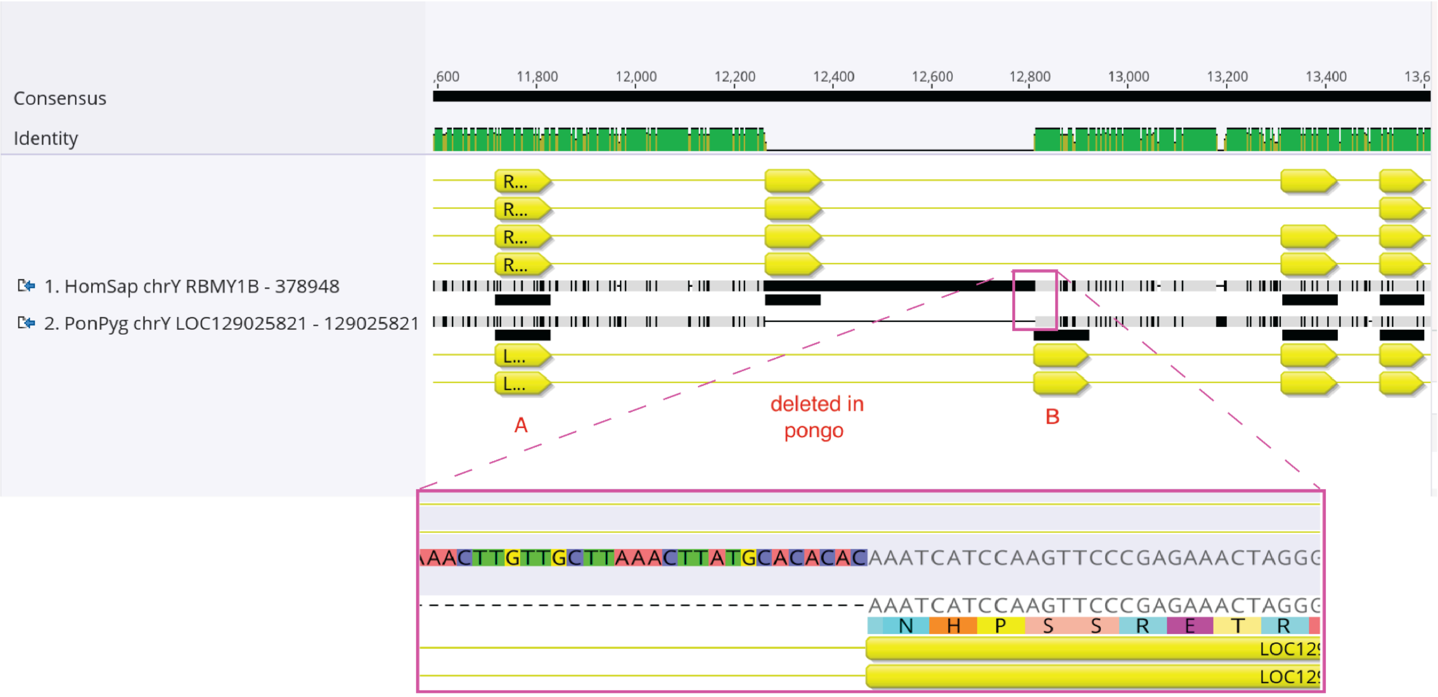

B

| | A_human_full | A_pongo_full | Deleted_in_... | B_human_full | B_pongo_full |
|---|---|---|---|---|---|
| A_human_full | | 92.50% | 92.70% | 72.53% | 71.73% |
| A_pongo_full | 92.50% | | 89.82% | 72.99% | 72.52% |
| Deleted_in_pongo_full | 92.70% | 89.82% | | 70.89% | 69.97% |
| B_human_full | 72.53% | 72.99% | 70.89% | | 89.04% |
| B_pongo_full | 71.73% | 72.52% | 69.97% | 89.04% | |

### Figure S6. Early initiation codon in *CDY* in the *Pongo* lineage

Alignment of *CDY* sequences from all seven species. Gene copies from Bornean and Sumatran orangutans do not cluster separately and are highlighted. We observe several orangutan-specific copies with an earlier initiation codon. Some copies have a mutation that affects the original initiation codon, highlighted in a red rectangle.

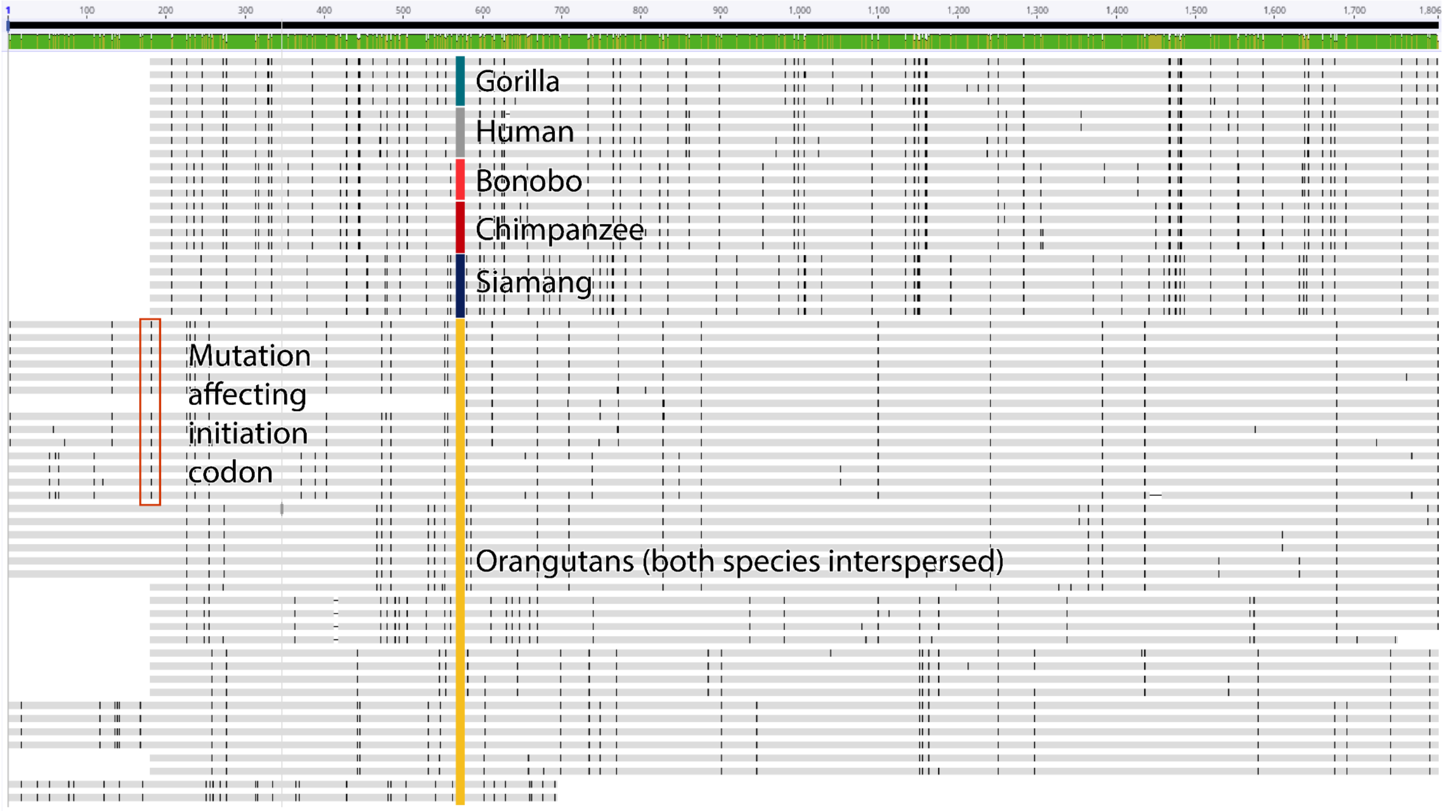

### Figure S7. Short read RNA-seq expression levels of YAGs

Expression levels of YAGs estimated from short read RNA-seq data using standard transcriptome (full colored bars) and using transcriptome augmented with newly identified isoforms (single black line).

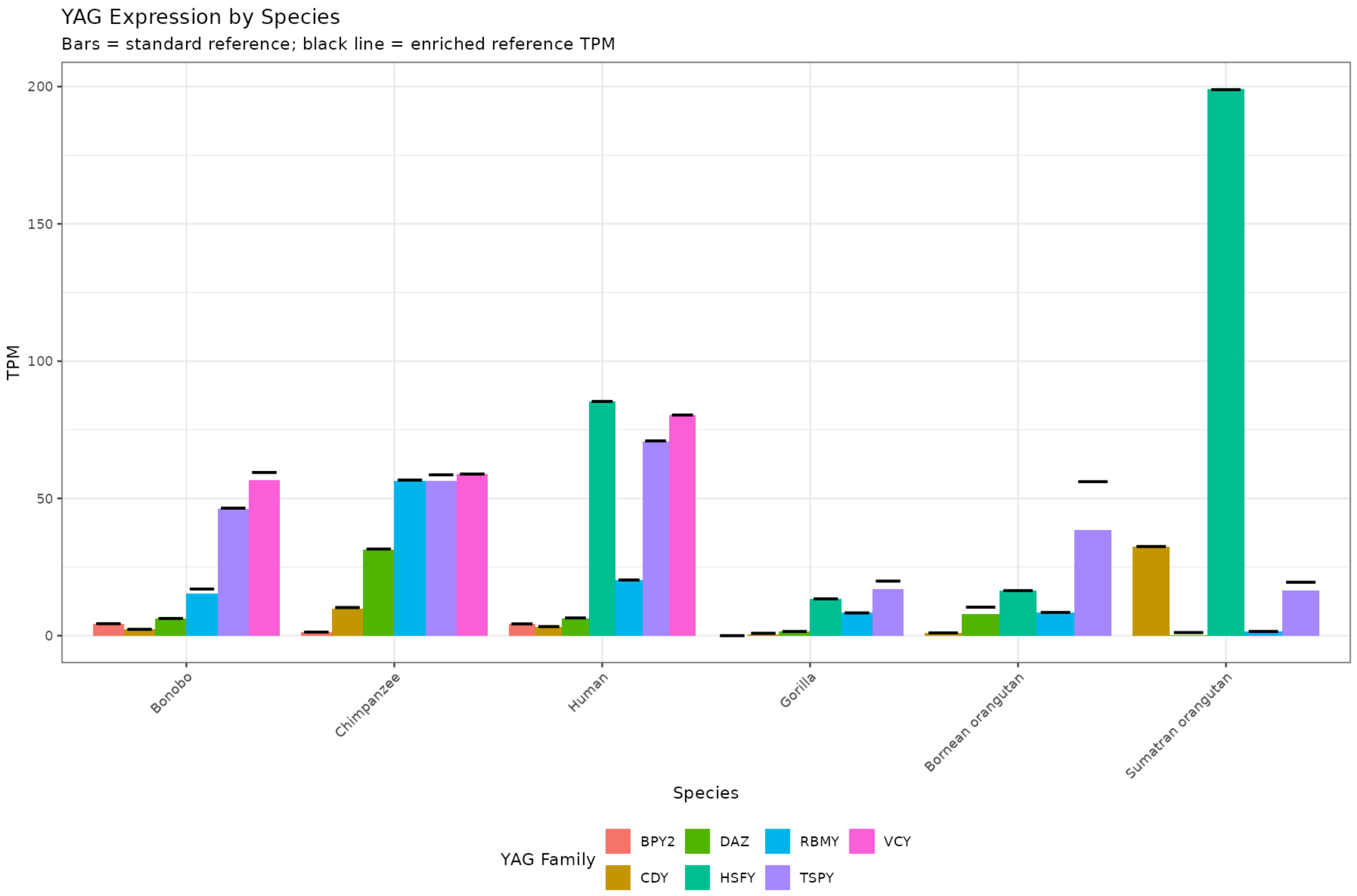

### Figure S8. Confidence levels of ColabFold predicted protein structures across YAG isoforms

(**A**) pLDDT (per-residue measure of local confidence) scores from AlphaFold2 (Jumper et al. 2021) were projected onto the genomic exon-intron diagrams (from Figure 3A). The resulting per-nucleotide pLDDT traces are rendered as colored line segments above each isoform, using a red–white–blue color scale shown in the figure.
(**B**) pLDDT (per-residue measure of local confidence) scores from AlphaFold2 (Jumper et al. 2021) were projected onto the diagrams of individual *DAZ* isoforms (from Figure 3C).

A

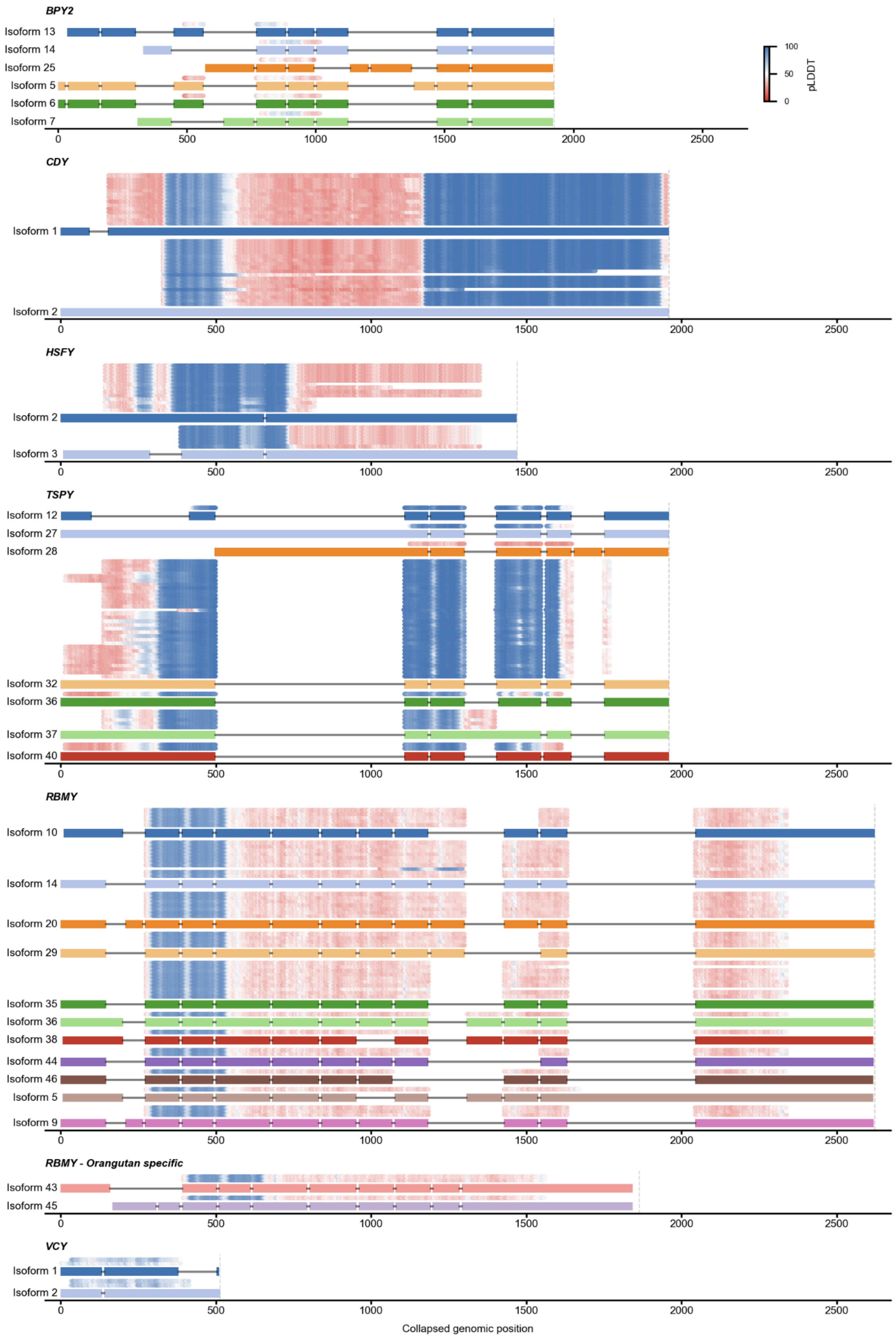

BPY2
Isoform 13
Isoform 14
Isoform 25
Isoform 5
Isoform 6
Isoform 7
100
50
0
pLDDT
CDY
Isoform 1
Isoform 2
HSFY
Isoform 2
Isoform 3
TSPY
Isoform 12
Isoform 27
Isoform 28
Isoform 32
Isoform 36
Isoform 37
Isoform 40
RBMY
Isoform 10
Isoform 14
Isoform 20
Isoform 29
Isoform 35
Isoform 36
Isoform 38
Isoform 44
Isoform 46
Isoform 5
Isoform 9
RBMY - Orangutan specific
Isoform 43
Isoform 45
VCY
Isoform 1
Isoform 2
0
500
1000
1500
2000
2500
Collapsed genomic position

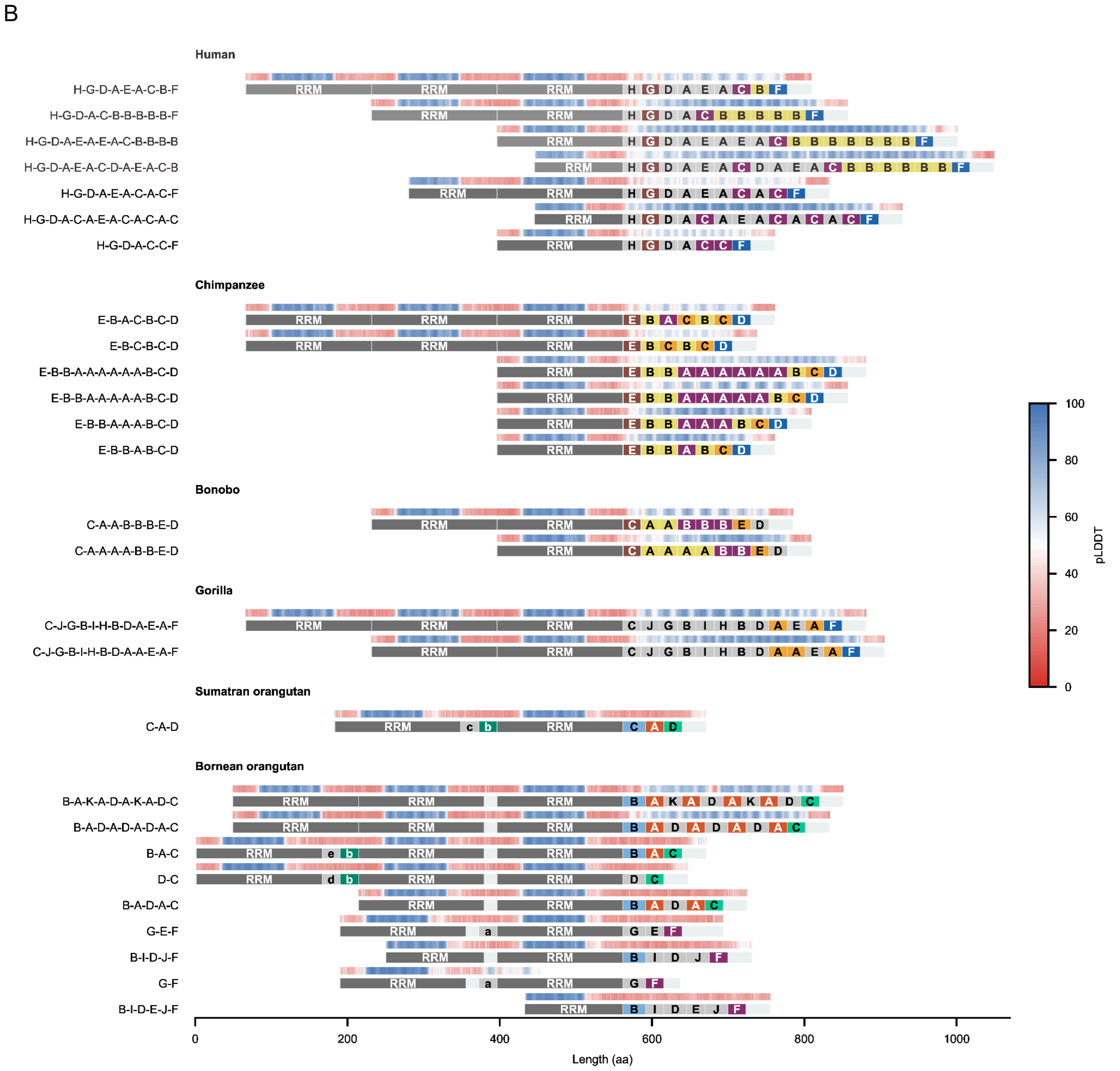

B
Human
H-G-D-A-E-A-C-B-F
H-G-D-A-C-B-B-B-B-B-F
H-G-D-A-E-A-E-A-C-B-B-B-B
H-G-D-A-E-A-C-D-A-E-A-C-B
H-G-D-A-E-A-C-A-C-F
H-G-D-A-C-A-E-A-C-A-C-A-C
H-G-D-A-C-C-F
Chimpanzee
E-B-A-C-B-C-D
E-B-C-B-C-D
E-B-B-A-A-A-A-A-A-B-C-D
E-B-B-A-A-A-A-A-B-C-D
E-B-B-A-A-A-B-C-D
E-B-B-A-B-C-D
Bonobo
C-A-A-B-B-B-E-D
C-A-A-A-A-B-B-E-D
Gorilla
C-J-G-B-I-H-B-D-A-E-A-F
C-J-G-B-I-H-B-D-A-A-E-A-F
Sumatran orangutan
C-A-D
Bornean orangutan
B-A-K-A-D-A-K-A-D-C
B-A-D-A-D-A-D-A-C
B-A-C
D-C
B-A-D-A-C
G-E-F
B-I-D-J-F
G-F
B-I-D-E-J-F
RRM
pLDDT
100
80
60
40
20
0
0
200
400
600
800
1000
Length (aa)

### Figure S9. Predicted protein structures.

A) *BPY2*
B) *VCY*
C) *RBMY* - *Pongo*-specific copies

A

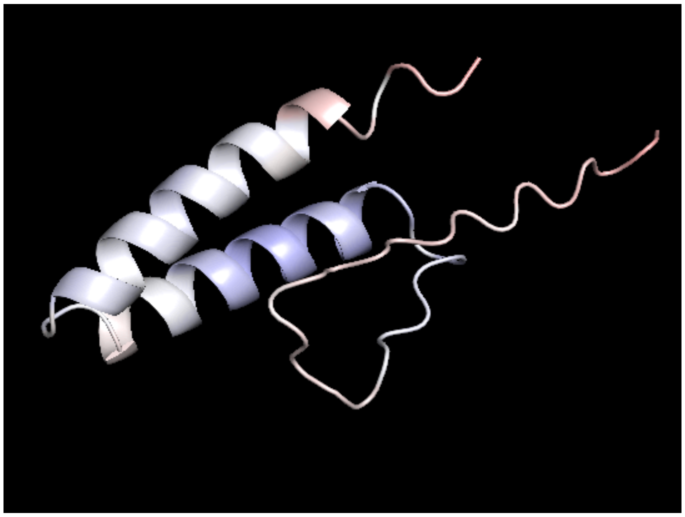

B

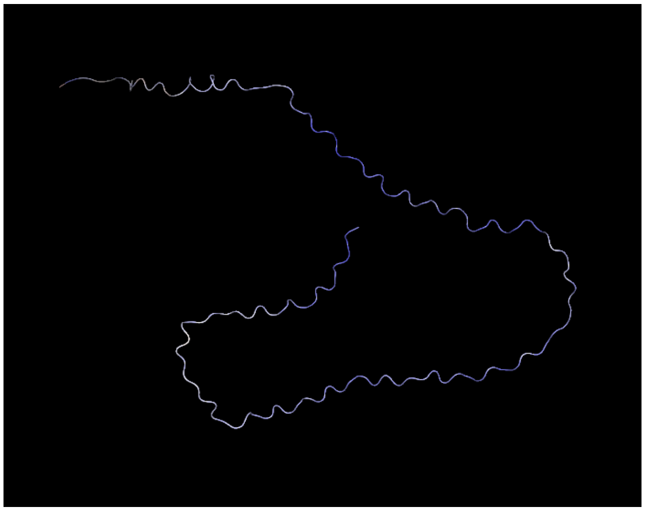

C

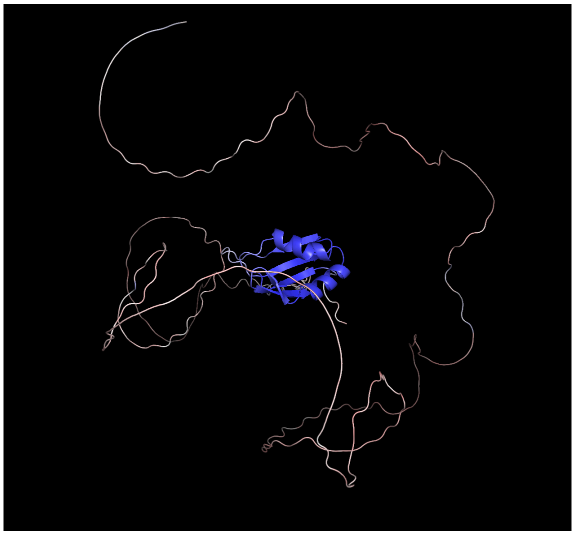

### Figure S10. Episodic diversifying selection on residue 331 of *CDY*

(**A**) Residue 331 in the cartoon representation of the folder protein in PyMol (Schrödinger, LLC 2026). (**B**) Two branches under diversifying selection at residue 331 are drawn with a thicker dark red stroke.

A

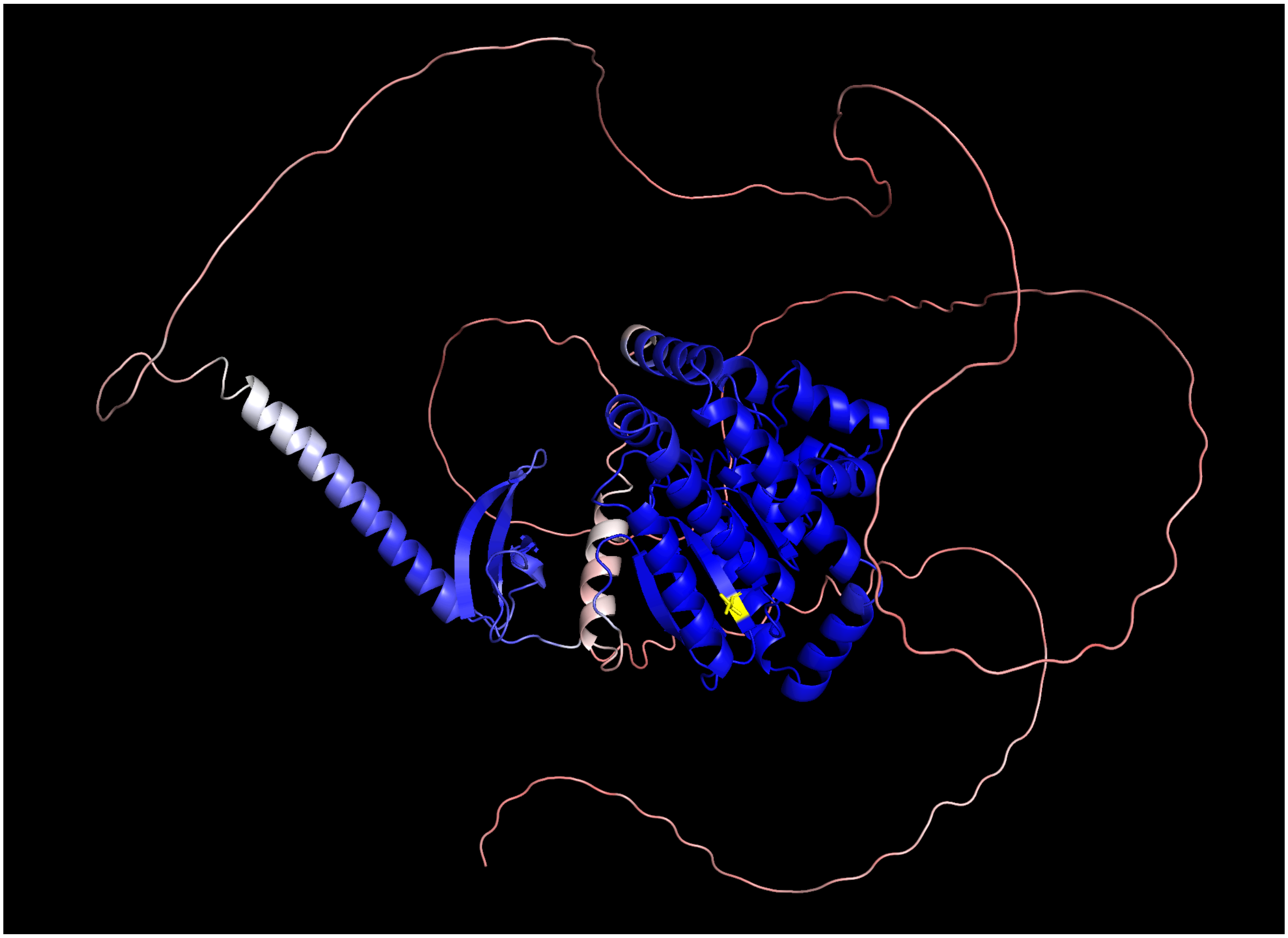

B

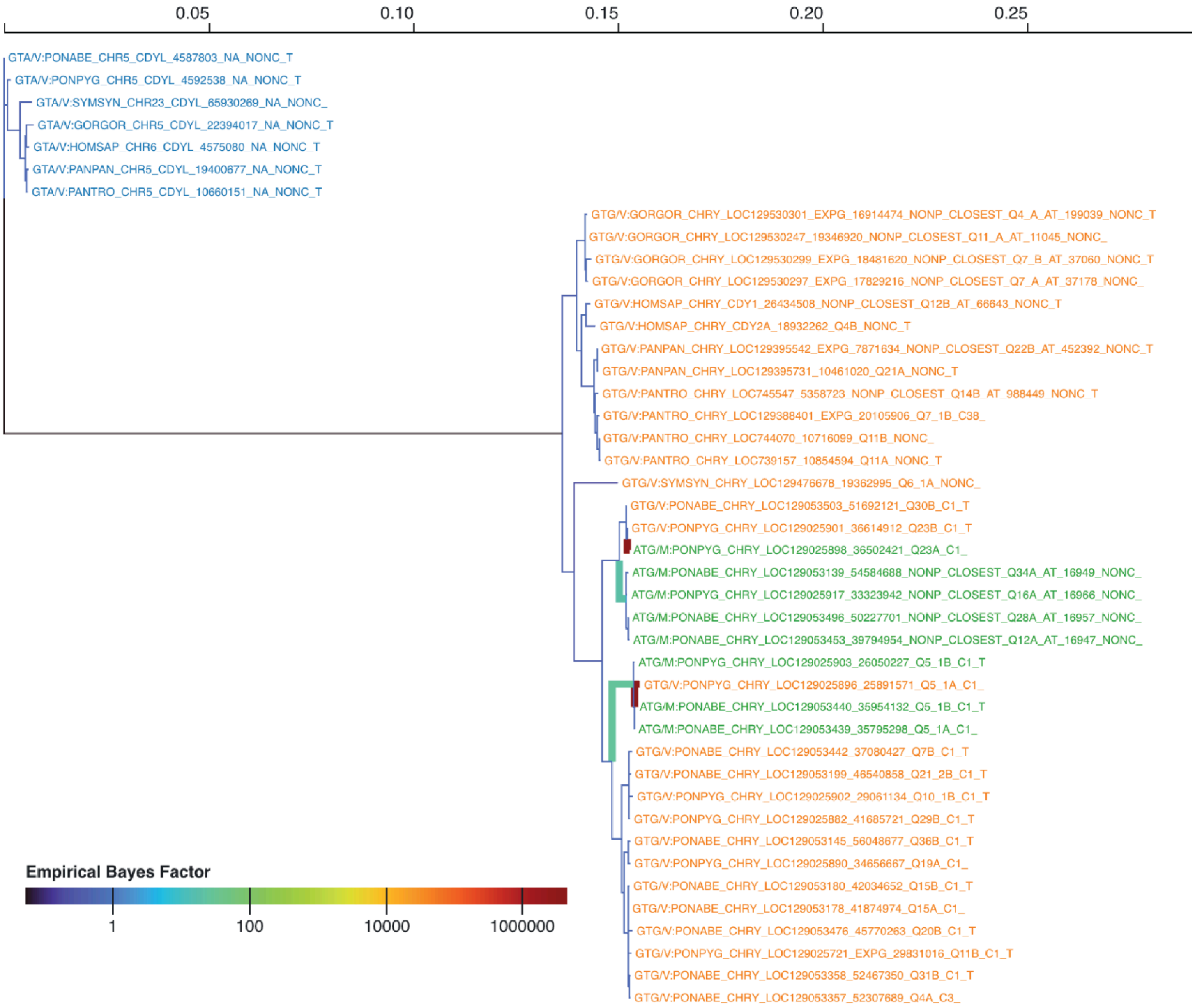
0.05
0.10
0.15
0.20
0.25
GTA/V:PONABE_CHR5_CDYL_4587803_NA_NONC_T
GTA/V:PONPYG_CHR5_CDYL_4592538_NA_NONC_T
GTA/V:SYMSYN_CHR23_CDYL_65930269_NA_NONC_
GTA/V:GORGOR_CHR5_CDYL_22394017_NA_NONC_T
GTA/V:HOMSAP_CHR6_CDYL_4575080_NA_NONC_T
GTA/V:PANPAN_CHR5_CDYL_19400677_NA_NONC_T
GTA/V:PANTRO_CHR5_CDYL_10660151_NA_NONC_T
GTG/V:GORGOR_CHRY_LOC129530301_EXPG_16914474_NONP_CLOSEST_Q4_A_AT_199039_NONC_T
GTG/V:GORGOR_CHRY_LOC129530247_19346920_NONP_CLOSEST_Q11_A_AT_11045_NONC_
GTG/V:GORGOR_CHRY_LOC129530299_EXPG_18481620_NONP_CLOSEST_Q7_B_AT_37060_NONC_T
GTG/V:GORGOR_CHRY_LOC129530297_EXPG_17829216_NONP_CLOSEST_Q7_A_AT_37178_NONC_
GTG/V:HOMSAP_CHRY_CDY1_26434508_NONP_CLOSEST_Q12B_AT_66643_NONC_T
GTG/V:HOMSAP_CHRY_CDY2A_18932262_Q4B_NONC_T
GTG/V:PANPAN_CHRY_LOC129395542_EXPG_7871634_NONP_CLOSEST_Q22B_AT_452392_NONC_T
GTG/V:PANPAN_CHRY_LOC129395731_10461020_Q21A_NONC_T
GTG/V:PANTRO_CHRY_LOC745547_5358723_NONP_CLOSEST_Q14B_AT_988449_NONC_T
GTG/V:PANTRO_CHRY_LOC129388401_EXPG_20105906_Q7_1B_C38_
GTG/V:PANTRO_CHRY_LOC744070_10716099_Q11B_NONC_
GTG/V:PANTRO_CHRY_LOC739157_10854594_Q11A_NONC_T
GTG/V:SYMSYN_CHRY_LOC129476678_19362995_Q6_1A_NONC_
GTG/V:PONABE_CHRY_LOC129053503_51692121_Q30B_C1_T
GTG/V:PONPYG_CHRY_LOC129025901_36614912_Q23B_C1_T
ATG/M:PONPYG_CHRY_LOC129025898_36502421_Q23A_C1_
ATG/M:PONABE_CHRY_LOC129053139_54584688_NONP_CLOSEST_Q34A_AT_16949_NONC_
ATG/M:PONPYG_CHRY_LOC129025917_33323942_NONP_CLOSEST_Q16A_AT_16966_NONC_
ATG/M:PONABE_CHRY_LOC129053496_50227701_NONP_CLOSEST_Q28A_AT_16957_NONC_
ATG/M:PONABE_CHRY_LOC129053453_39794954_NONP_CLOSEST_Q12A_AT_16947_NONC_
ATG/M:PONPYG_CHRY_LOC129025903_26050227_Q5_1B_C1_T
GTG/V:PONPYG_CHRY_LOC129025896_25891571_Q5_1A_C1_
ATG/M:PONABE_CHRY_LOC129053440_35954132_Q5_1B_C1_T
ATG/M:PONABE_CHRY_LOC129053439_35795298_Q5_1A_C1_
GTG/V:PONABE_CHRY_LOC129053442_37080427_Q7B_C1_T
GTG/V:PONABE_CHRY_LOC129053199_46540858_Q21_2B_C1_T
GTG/V:PONPYG_CHRY_LOC129025902_29061134_Q10_1B_C1_T
GTG/V:PONPYG_CHRY_LOC129025882_41685721_Q29B_C1_T
GTG/V:PONABE_CHRY_LOC129053145_56048677_Q36B_C1_T
GTG/V:PONPYG_CHRY_LOC129025890_34656667_Q19A_C1_
GTG/V:PONABE_CHRY_LOC129053180_42034652_Q15B_C1_T
GTG/V:PONABE_CHRY_LOC129053178_41874974_Q15A_C1_
GTG/V:PONABE_CHRY_LOC129053476_45770263_Q20B_C1_T
GTG/V:PONPYG_CHRY_LOC129025721_EXPG_29831016_Q11B_C1_T
GTG/V:PONABE_CHRY_LOC129053358_52467350_Q31B_C1_T
GTG/V:PONABE_CHRY_LOC129053357_52307689_Q4A_C3_
Empirical Bayes Factor
1
100
10000
1000000

### Figure S11. Episodic diversifying selection on individual sites of the *CDY* residue 497

(**A**) Residue 497 in the cartoon representation of the folder protein in PyMol (Schrödinger, LLC 2026). (**B**) One branch under diversifying selection at residue 475 drawn with a thicker dark red stroke.

A

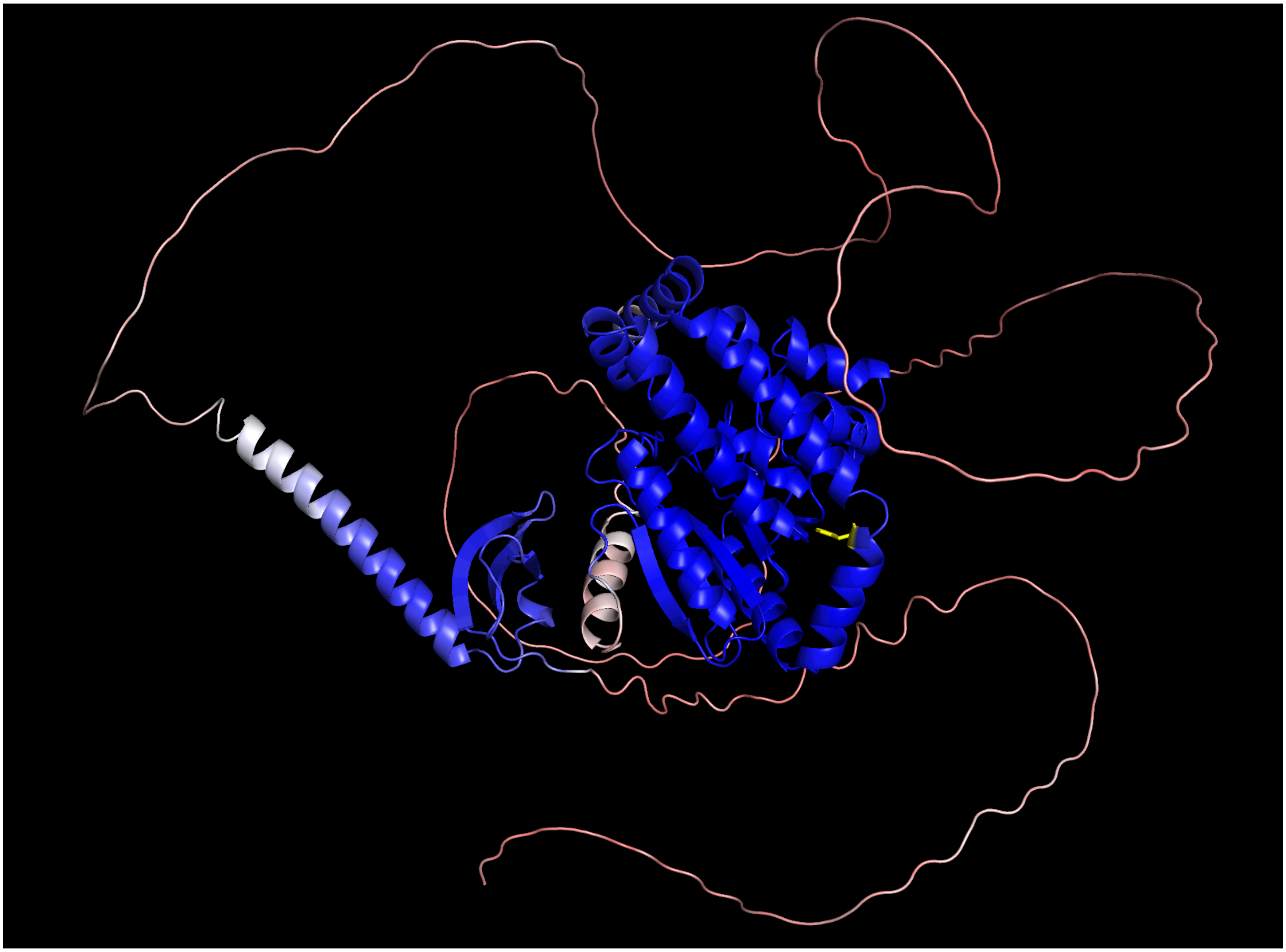

B

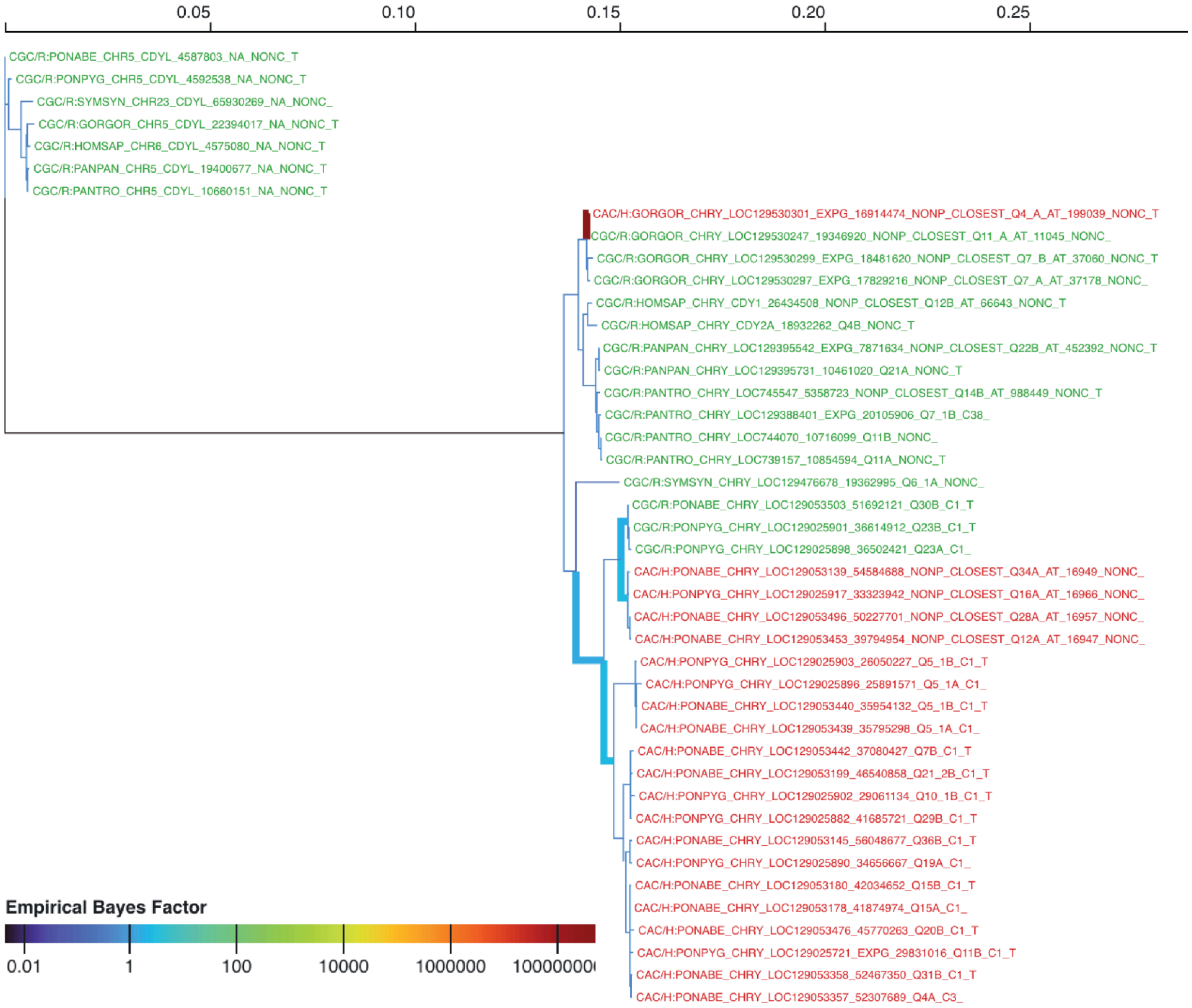
0.05
0.10
0.15
0.20
0.25
CGC/R:PONABE_CHR5_CDYL_4587803_NA_NONC_T
CGC/R:PONPYG_CHR5_CDYL_4592538_NA_NONC_T
CGC/R:SYMSYN_CHR23_CDYL_65930269_NA_NONC_
CGC/R:GORGOR_CHR5_CDYL_22394017_NA_NONC_T
CGC/R:HOMSAP_CHR6_CDYL_4575080_NA_NONC_T
CGC/R:PANPAN_CHR5_CDYL_19400677_NA_NONC_T
CGC/R:PANTRO_CHR5_CDYL_10660151_NA_NONC_T
CAC/H:GORGOR_CHRY_LOC129530301_EXPG_16914474_NONP_CLOSEST_Q4_A_AT_199039_NONC_T
CGC/R:GORGOR_CHRY_LOC129530247_19346920_NONP_CLOSEST_Q11_A_AT_11045_NONC_
CGC/R:GORGOR_CHRY_LOC129530299_EXPG_18481620_NONP_CLOSEST_Q7_B_AT_37060_NONC_T
CGC/R:GORGOR_CHRY_LOC129530297_EXPG_17829216_NONP_CLOSEST_Q7_A_AT_37178_NONC_
CGC/R:HOMSAP_CHRY_CDY1_26434508_NONP_CLOSEST_Q12B_AT_66643_NONC_T
CGC/R:HOMSAP_CHRY_CDY2A_18932262_Q4B_NONC_T
CGC/R:PANPAN_CHRY_LOC129395542_EXPG_7871634_NONP_CLOSEST_Q22B_AT_452392_NONC_T
CGC/R:PANPAN_CHRY_LOC129395731_10461020_Q21A_NONC_T
CGC/R:PANTRO_CHRY_LOC745547_5358723_NONP_CLOSEST_Q14B_AT_988449_NONC_T
CGC/R:PANTRO_CHRY_LOC129388401_EXPG_20105906_Q7_1B_C38_
CGC/R:PANTRO_CHRY_LOC744070_10716099_Q11B_NONC_
CGC/R:PANTRO_CHRY_LOC739157_10854594_Q11A_NONC_T
CGC/R:SYMSYN_CHRY_LOC129476678_19362995_Q6_1A_NONC_
CGC/R:PONABE_CHRY_LOC129053503_51692121_Q30B_C1_T
CGC/R:PONPYG_CHRY_LOC129025901_36614912_Q23B_C1_T
CGC/R:PONPYG_CHRY_LOC129025898_36502421_Q23A_C1_
CAC/H:PONABE_CHRY_LOC129053139_54584688_NONP_CLOSEST_Q34A_AT_16949_NONC_
CAC/H:PONPYG_CHRY_LOC129025917_33323942_NONP_CLOSEST_Q16A_AT_16966_NONC_
CAC/H:PONABE_CHRY_LOC129053496_50227701_NONP_CLOSEST_Q28A_AT_16957_NONC_
CAC/H:PONABE_CHRY_LOC129053453_39794954_NONP_CLOSEST_Q12A_AT_16947_NONC_
CAC/H:PONPYG_CHRY_LOC129025903_26050227_Q5_1B_C1_T
CAC/H:PONPYG_CHRY_LOC129025896_25891571_Q5_1A_C1_
CAC/H:PONABE_CHRY_LOC129053440_35954132_Q5_1B_C1_T
CAC/H:PONABE_CHRY_LOC129053439_35795298_Q5_1A_C1_
CAC/H:PONABE_CHRY_LOC129053442_37080427_Q7B_C1_T
CAC/H:PONABE_CHRY_LOC129053199_46540858_Q21_2B_C1_T
CAC/H:PONPYG_CHRY_LOC129025902_29061134_Q10_1B_C1_T
CAC/H:PONPYG_CHRY_LOC129025882_41685721_Q29B_C1_T
CAC/H:PONABE_CHRY_LOC129053145_56048677_Q36B_C1_T
CAC/H:PONPYG_CHRY_LOC129025890_34656667_Q19A_C1_
CAC/H:PONABE_CHRY_LOC129053180_42034652_Q15B_C1_T
CAC/H:PONABE_CHRY_LOC129053178_41874974_Q15A_C1_
CAC/H:PONABE_CHRY_LOC129053476_45770263_Q20B_C1_T
CAC/H:PONPYG_CHRY_LOC129025721_EXPG_29831016_Q11B_C1_T
CAC/H:PONABE_CHRY_LOC129053358_52467350_Q31B_C1_T
CAC/H:PONABE_CHRY_LOC129053357_52307689_Q4A_C3_
Empirical Bayes Factor
0.01
1
100
10000
1000000

#### Figure S12. Episodic diversifying selection on individual sites of the *RBMY* residue 13

(**A**) Residue 13 in the cartoon representation of the folder protein in PyMol (Schrödinger, LLC 2026). (**B**) Two branches under diversifying selection at residue 13 are drawn with a thicker lime green and dark red stroke.

A

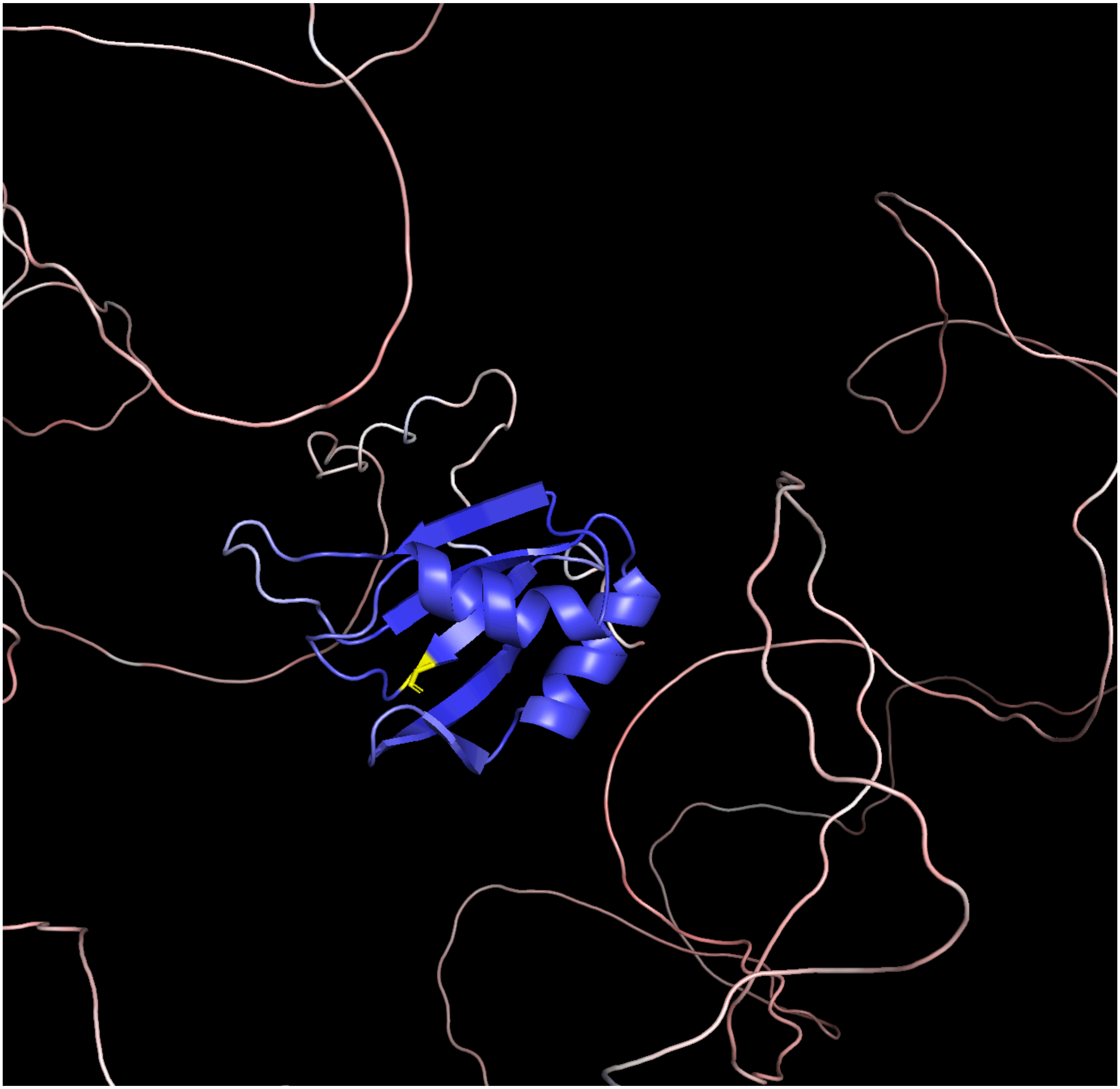

B

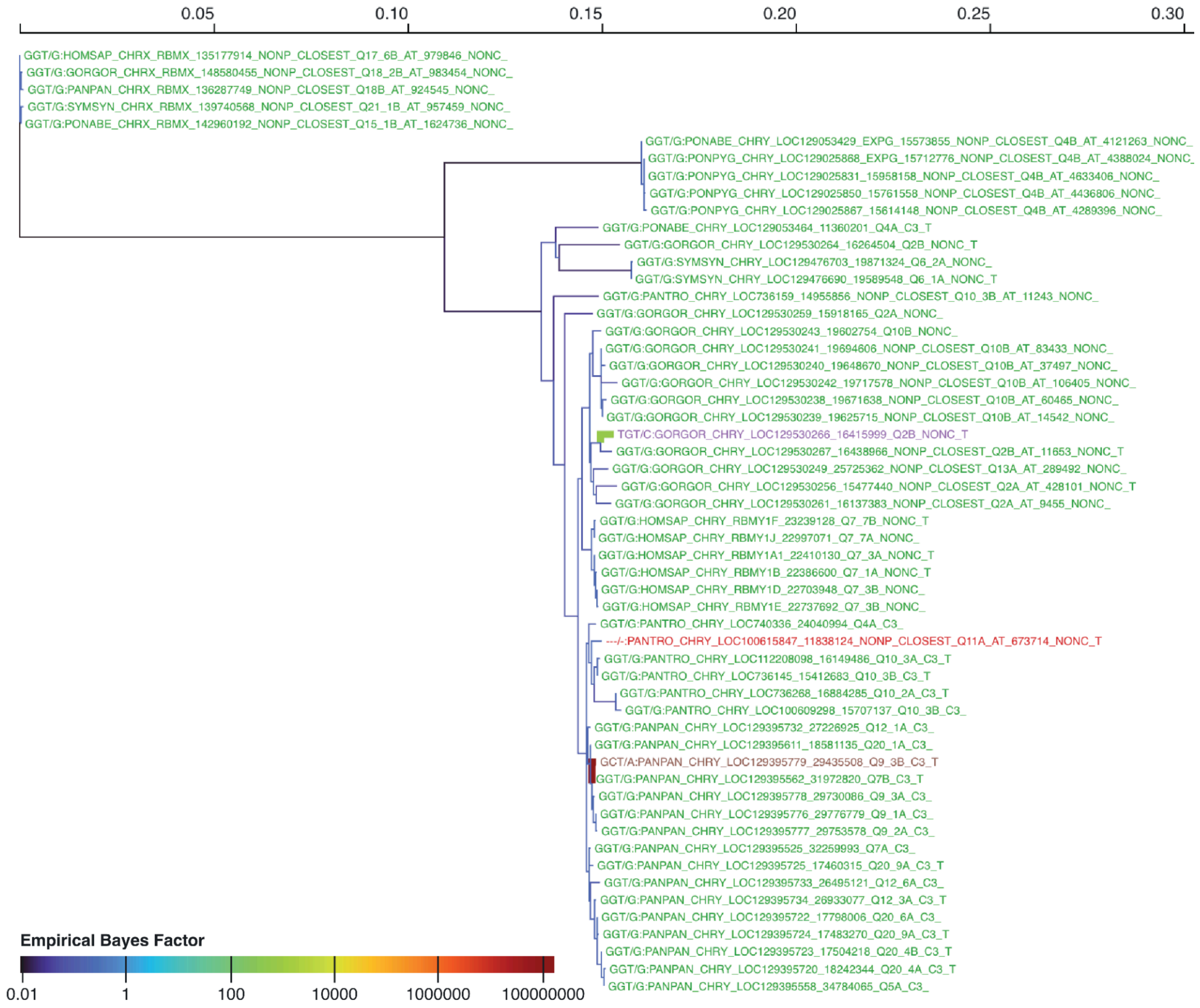

0.05
0.10
0.15
0.20
0.25
0.30
GGT/G:HOMSAP_CHRX_RBMX_135177914_NONP_CLOSEST_Q17_6B_AT_979846_NONC_
GGT/G:GORGOR_CHRX_RBMX_148580455_NONP_CLOSEST_Q18_2B_AT_983454_NONC_
GGT/G:PANPAN_CHRX_RBMX_136287749_NONP_CLOSEST_Q18B_AT_924545_NONC_
GGT/G:SYMSYN_CHRX_RBMX_139740568_NONP_CLOSEST_Q21_1B_AT_957459_NONC_
GGT/G:PONABE_CHRX_RBMX_142960192_NONP_CLOSEST_Q15_1B_AT_1624736_NONC_
GGT/G:PONABE_CHRY_LOC129053429_EXPG_15573855_NONP_CLOSEST_Q4B_AT_4121263_NONC_
GGT/G:PONPYG_CHRY_LOC129025888_EXPG_15712776_NONP_CLOSEST_Q4B_AT_4388024_NONC_
GGT/G:PONPYG_CHRY_LOC129025831_15958158_NONP_CLOSEST_Q4B_AT_4633406_NONC_
GGT/G:PONPYG_CHRY_LOC129025850_15761558_NONP_CLOSEST_Q4B_AT_4436806_NONC_
GGT/G:PONPYG_CHRY_LOC129025867_15614148_NONP_CLOSEST_Q4B_AT_4289396_NONC_
GGT/G:PONABE_CHRY_LOC129053464_11360201_Q4A_C3_T
GGT/G:GORGOR_CHRY_LOC129530264_16264504_Q2B_NONC_T
GGT/G:SYMSYN_CHRY_LOC129476703_19871324_Q6_2A_NONC_
GGT/G:SYMSYN_CHRY_LOC129476690_19589548_Q6_1A_NONC_T
GGT/G:PANTRO_CHRY_LOC736159_14955856_NONP_CLOSEST_Q10_3B_AT_11243_NONC_
GGT/G:GORGOR_CHRY_LOC129530259_15918165_Q2A_NONC_
GGT/G:GORGOR_CHRY_LOC129530243_19602754_Q10B_NONC_
GGT/G:GORGOR_CHRY_LOC129530241_19694606_NONP_CLOSEST_Q10B_AT_83433_NONC_
GGT/G:GORGOR_CHRY_LOC129530240_19648670_NONP_CLOSEST_Q10B_AT_37497_NONC_
GGT/G:GORGOR_CHRY_LOC129530242_19717578_NONP_CLOSEST_Q10B_AT_106405_NONC_
GGT/G:GORGOR_CHRY_LOC129530238_19671638_NONP_CLOSEST_Q10B_AT_60465_NONC_
GGT/G:GORGOR_CHRY_LOC129530239_19625715_NONP_CLOSEST_Q10B_AT_14542_NONC_
TGT/C:GORGOR_CHRY_LOC129530266_16415999_Q2B_NONC_T
GGT/G:GORGOR_CHRY_LOC129530267_16438966_NONP_CLOSEST_Q2B_AT_11653_NONC_T
GGT/G:GORGOR_CHRY_LOC129530249_25725362_NONP_CLOSEST_Q13A_AT_289492_NONC_
GGT/G:GORGOR_CHRY_LOC129530256_15477440_NONP_CLOSEST_Q2A_AT_428101_NONC_T
GGT/G:GORGOR_CHRY_LOC129530261_16137383_NONP_CLOSEST_Q2A_AT_9455_NONC_
GGT/G:HOMSAP_CHRY_RBMY1F_23239128_Q7_7B_NONC_T
GGT/G:HOMSAP_CHRY_RBMY1J_22997071_Q7_7A_NONC_
GGT/G:HOMSAP_CHRY_RBMY1A1_22410130_Q7_3A_NONC_T
GGT/G:HOMSAP_CHRY_RBMY1B_22386600_Q7_1A_NONC_T
GGT/G:HOMSAP_CHRY_RBMY1D_22703948_Q7_3B_NONC_
GGT/G:HOMSAP_CHRY_RBMY1E_22737692_Q7_3B_NONC_
GGT/G:PANTRO_CHRY_LOC740336_24040994_Q4A_C3_
---/-:PANTRO_CHRY_LOC100615847_11838124_NONP_CLOSEST_Q11A_AT_673714_NONC_T
GGT/G:PANTRO_CHRY_LOC112208098_16149486_Q10_3A_C3_T
GGT/G:PANTRO_CHRY_LOC736145_15412683_Q10_3B_C3_T
GGT/G:PANTRO_CHRY_LOC736268_16884285_Q10_2A_C3_T
GGT/G:PANTRO_CHRY_LOC100609298_15707137_Q10_3B_C3_
GGT/G:PANPAN_CHRY_LOC129395732_27226925_Q12_1A_C3_
GGT/G:PANPAN_CHRY_LOC129395611_18581135_Q20_1A_C3_
GCT/A:PANPAN_CHRY_LOC129395779_29435508_Q9_3B_C3_T
GGT/G:PANPAN_CHRY_LOC129395562_31972820_Q7B_C3_T
GGT/G:PANPAN_CHRY_LOC129395778_29730086_Q9_3A_C3_
GGT/G:PANPAN_CHRY_LOC129395776_29776779_Q9_1A_C3_
GGT/G:PANPAN_CHRY_LOC129395777_29753578_Q9_2A_C3_
GGT/G:PANPAN_CHRY_LOC129395525_32259993_Q7A_C3_
GGT/G:PANPAN_CHRY_LOC129395725_17460315_Q20_9A_C3_T
GGT/G:PANPAN_CHRY_LOC129395733_26495121_Q12_6A_C3_
GGT/G:PANPAN_CHRY_LOC129395734_26933077_Q12_3A_C3_T
GGT/G:PANPAN_CHRY_LOC129395722_17798006_Q20_6A_C3_
GGT/G:PANPAN_CHRY_LOC129395724_17483270_Q20_9A_C3_T
GGT/G:PANPAN_CHRY_LOC129395723_17504218_Q20_4B_C3_T
GGT/G:PANPAN_CHRY_LOC129395720_18242344_Q20_4A_C3_T
GGT/G:PANPAN_CHRY_LOC129395558_34784065_Q5A_C3_
Empirical Bayes Factor
0.01
1
100
10000
1000000
100000000

#### Figure S13. Episodic diversifying selection on individual sites of the *RBMY* residue 72

(**A**) Residue 72 in the cartoon representation of the folder protein in PyMol (Schrödinger, LLC 2026). (**B**) Two branches under diversifying selection at residue 72 are drawn with a thicker dark red stroke.

A

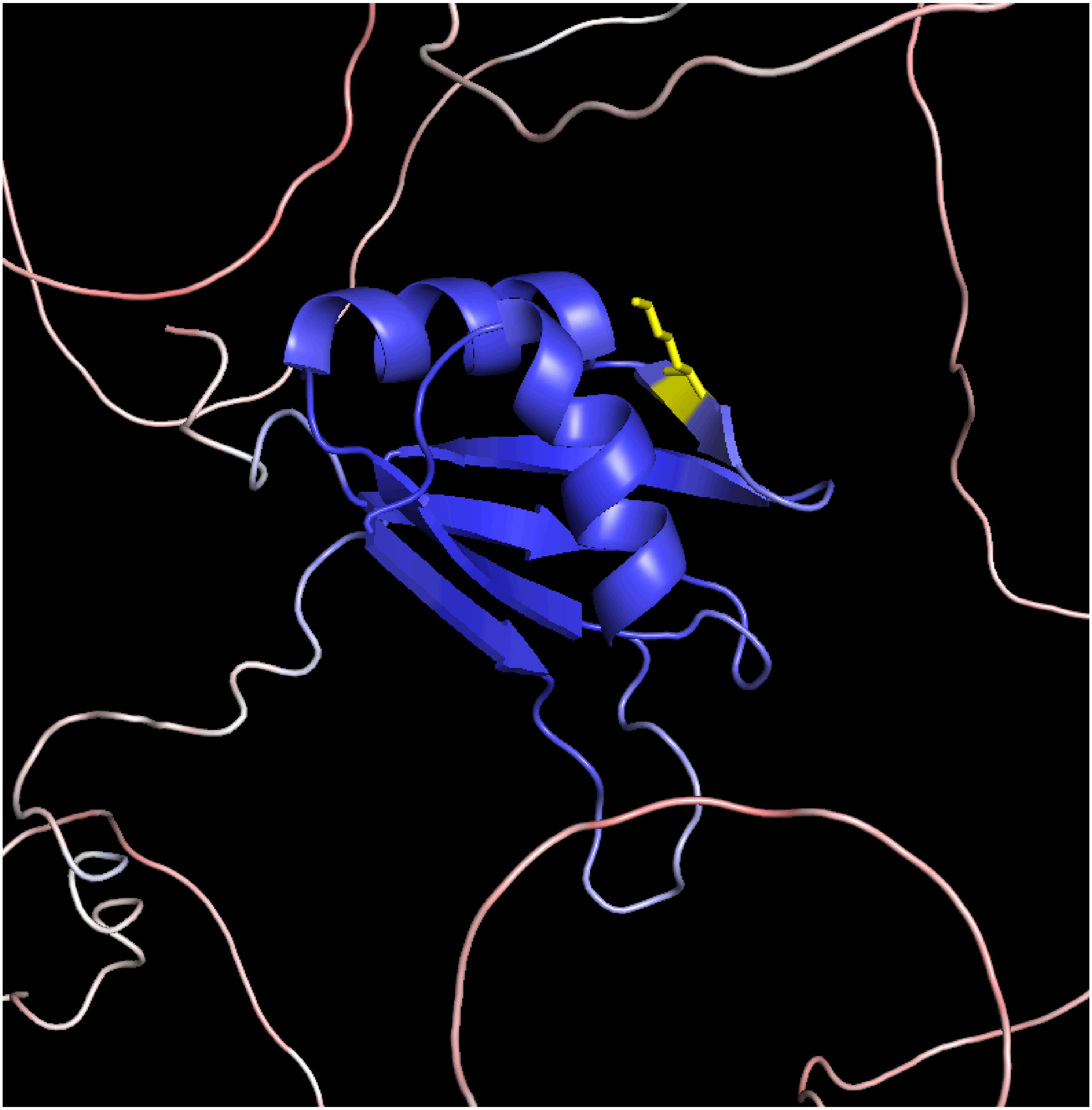

B

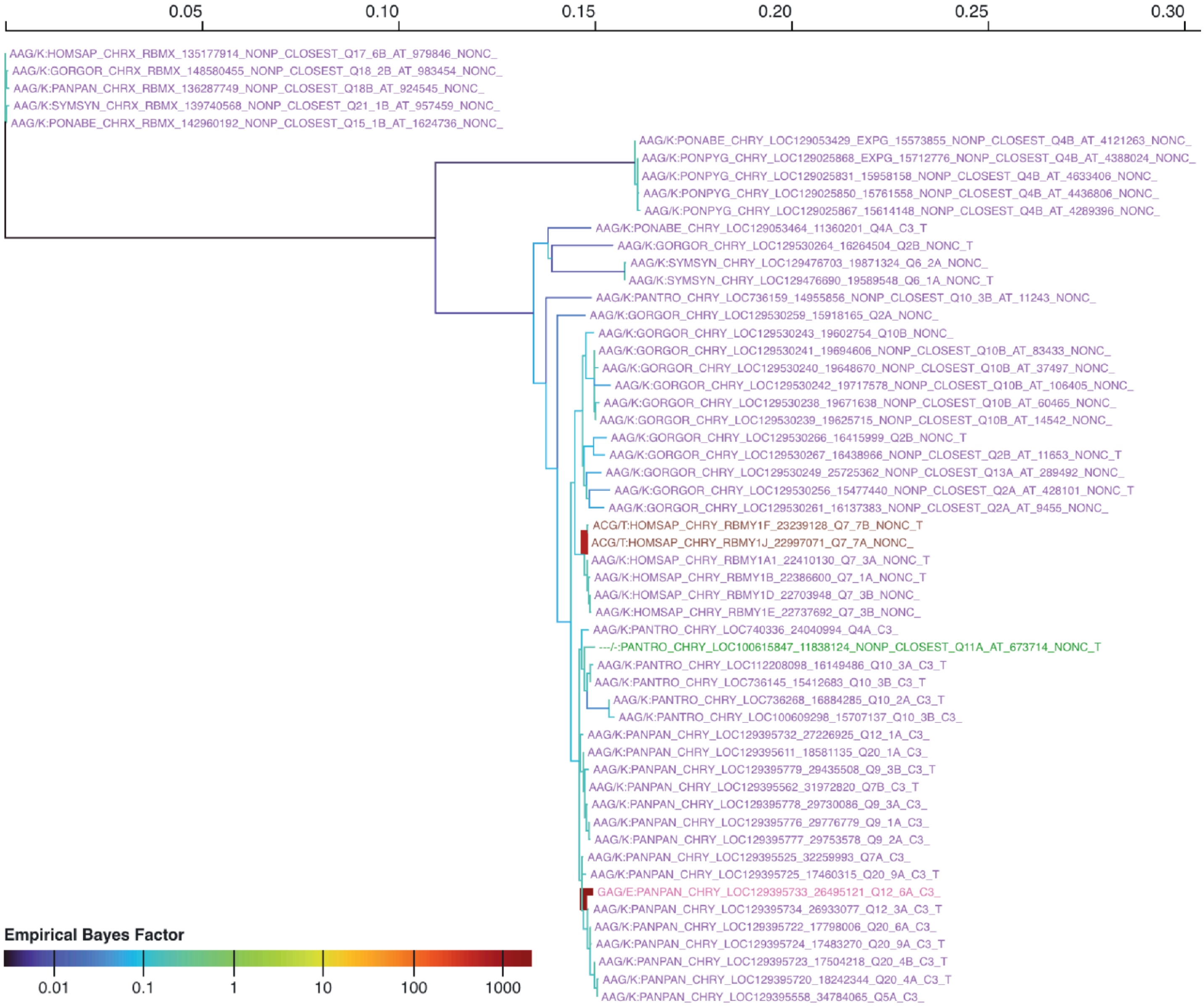

0.05
0.10
0.15
0.20
0.25
0.30
AAG/K:HOMSAP_CHRX_RBMX_135177914_NONP_CLOSEST_Q17_6B_AT_979846_NONC_
AAG/K:GORGOR_CHRX_RBMX_148580455_NONP_CLOSEST_Q18_2B_AT_983454_NONC_
AAG/K:PANPAN_CHRX_RBMX_136287749_NONP_CLOSEST_Q18B_AT_924545_NONC_
AAG/K:SYMSYN_CHRX_RBMX_139740568_NONP_CLOSEST_Q21_1B_AT_957459_NONC_
AAG/K:PONABE_CHRX_RBMX_142960192_NONP_CLOSEST_Q15_1B_AT_1624736_NONC_
AAG/K:PONABE_CHRY_LOC129053429_EXPG_15573855_NONP_CLOSEST_Q4B_AT_4121263_NONC_
AAG/K:PONPYG_CHRY_LOC129025868_EXPG_15712776_NONP_CLOSEST_Q4B_AT_4388024_NONC_
AAG/K:PONPYG_CHRY_LOC129025831_15958158_NONP_CLOSEST_Q4B_AT_4633406_NONC_
AAG/K:PONPYG_CHRY_LOC129025850_15761558_NONP_CLOSEST_Q4B_AT_4436806_NONC_
AAG/K:PONPYG_CHRY_LOC129025867_15614148_NONP_CLOSEST_Q4B_AT_4289396_NONC_
AAG/K:PONABE_CHRY_LOC129053464_11360201_Q4A_C3_T
AAG/K:GORGOR_CHRY_LOC129530264_16264504_Q2B_NONC_T
AAG/K:SYMSYN_CHRY_LOC129476703_19871324_Q6_2A_NONC_
AAG/K:SYMSYN_CHRY_LOC129476690_19589548_Q6_1A_NONC_T
AAG/K:PANTRO_CHRY_LOC736159_14955856_NONP_CLOSEST_Q10_3B_AT_11243_NONC_
AAG/K:GORGOR_CHRY_LOC129530259_15918165_Q2A_NONC_
AAG/K:GORGOR_CHRY_LOC129530243_19602754_Q10B_NONC_
AAG/K:GORGOR_CHRY_LOC129530241_19694606_NONP_CLOSEST_Q10B_AT_83433_NONC_
AAG/K:GORGOR_CHRY_LOC129530240_19648670_NONP_CLOSEST_Q10B_AT_37497_NONC_
AAG/K:GORGOR_CHRY_LOC129530242_19717578_NONP_CLOSEST_Q10B_AT_106405_NONC_
AAG/K:GORGOR_CHRY_LOC129530238_19671638_NONP_CLOSEST_Q10B_AT_60465_NONC_
AAG/K:GORGOR_CHRY_LOC129530239_19625715_NONP_CLOSEST_Q10B_AT_14542_NONC_
AAG/K:GORGOR_CHRY_LOC129530266_16415999_Q2B_NONC_T
AAG/K:GORGOR_CHRY_LOC129530267_16438966_NONP_CLOSEST_Q2B_AT_11653_NONC_T
AAG/K:GORGOR_CHRY_LOC129530249_25725362_NONP_CLOSEST_Q13A_AT_289492_NONC_
AAG/K:GORGOR_CHRY_LOC129530256_15477440_NONP_CLOSEST_Q2A_AT_428101_NONC_T
AAG/K:GORGOR_CHRY_LOC129530261_16137383_NONP_CLOSEST_Q2A_AT_9455_NONC_
ACG/T:HOMSAP_CHRY_RBMY1F_23239128_Q7_7B_NONC_T
ACG/T:HOMSAP_CHRY_RBMY1J_22997071_Q7_7A_NONC_
AAG/K:HOMSAP_CHRY_RBMY1A1_22410130_Q7_3A_NONC_T
AAG/K:HOMSAP_CHRY_RBMY1B_22386600_Q7_1A_NONC_T
AAG/K:HOMSAP_CHRY_RBMY1D_22703948_Q7_3B_NONC_
AAG/K:HOMSAP_CHRY_RBMY1E_22737692_Q7_3B_NONC_
AAG/K:PANTRO_CHRY_LOC740336_24040994_Q4A_C3_
---/-:PANTRO_CHRY_LOC100615847_11838124_NONP_CLOSEST_Q11A_AT_673714_NONC_T
AAG/K:PANTRO_CHRY_LOC112208098_16149486_Q10_3A_C3_T
AAG/K:PANTRO_CHRY_LOC736145_15412683_Q10_3B_C3_T
AAG/K:PANTRO_CHRY_LOC736268_16884285_Q10_2A_C3_T
AAG/K:PANTRO_CHRY_LOC100609298_15707137_Q10_3B_C3_
AAG/K:PANPAN_CHRY_LOC129395732_27226925_Q12_1A_C3_
AAG/K:PANPAN_CHRY_LOC129395611_18581135_Q20_1A_C3_
AAG/K:PANPAN_CHRY_LOC129395779_29435508_Q9_3B_C3_T
AAG/K:PANPAN_CHRY_LOC129395562_31972820_Q7B_C3_T
AAG/K:PANPAN_CHRY_LOC129395778_29730086_Q9_3A_C3_
AAG/K:PANPAN_CHRY_LOC129395776_29776779_Q9_1A_C3_
AAG/K:PANPAN_CHRY_LOC129395777_29753578_Q9_2A_C3_
AAG/K:PANPAN_CHRY_LOC129395525_32259993_Q7A_C3_
AAG/K:PANPAN_CHRY_LOC129395725_17460315_Q20_9A_C3_T
GAG/E:PANPAN_CHRY_LOC129395733_26495121_Q12_6A_C3_
AAG/K:PANPAN_CHRY_LOC129395734_26933077_Q12_3A_C3_T
AAG/K:PANPAN_CHRY_LOC129395722_17798006_Q20_6A_C3_
AAG/K:PANPAN_CHRY_LOC129395724_17483270_Q20_9A_C3_T
AAG/K:PANPAN_CHRY_LOC129395723_17504218_Q20_4B_C3_T
AAG/K:PANPAN_CHRY_LOC129395720_18242344_Q20_4A_C3_T
AAG/K:PANPAN_CHRY_LOC129395558_34784065_Q5A_C3_
Empirical Bayes Factor
0.01
0.1
1
10
100
1000

### Figure S14. Episodic diversifying selection on individual sites of the *RBMY* residue 92

(**A**) Residue 92 in the cartoon representation of the folder protein in PyMol (Schrödinger, LLC 2026). (**B**) Two branches under diversifying selection at residue 92 are drawn with a thicker dark red stroke.

A

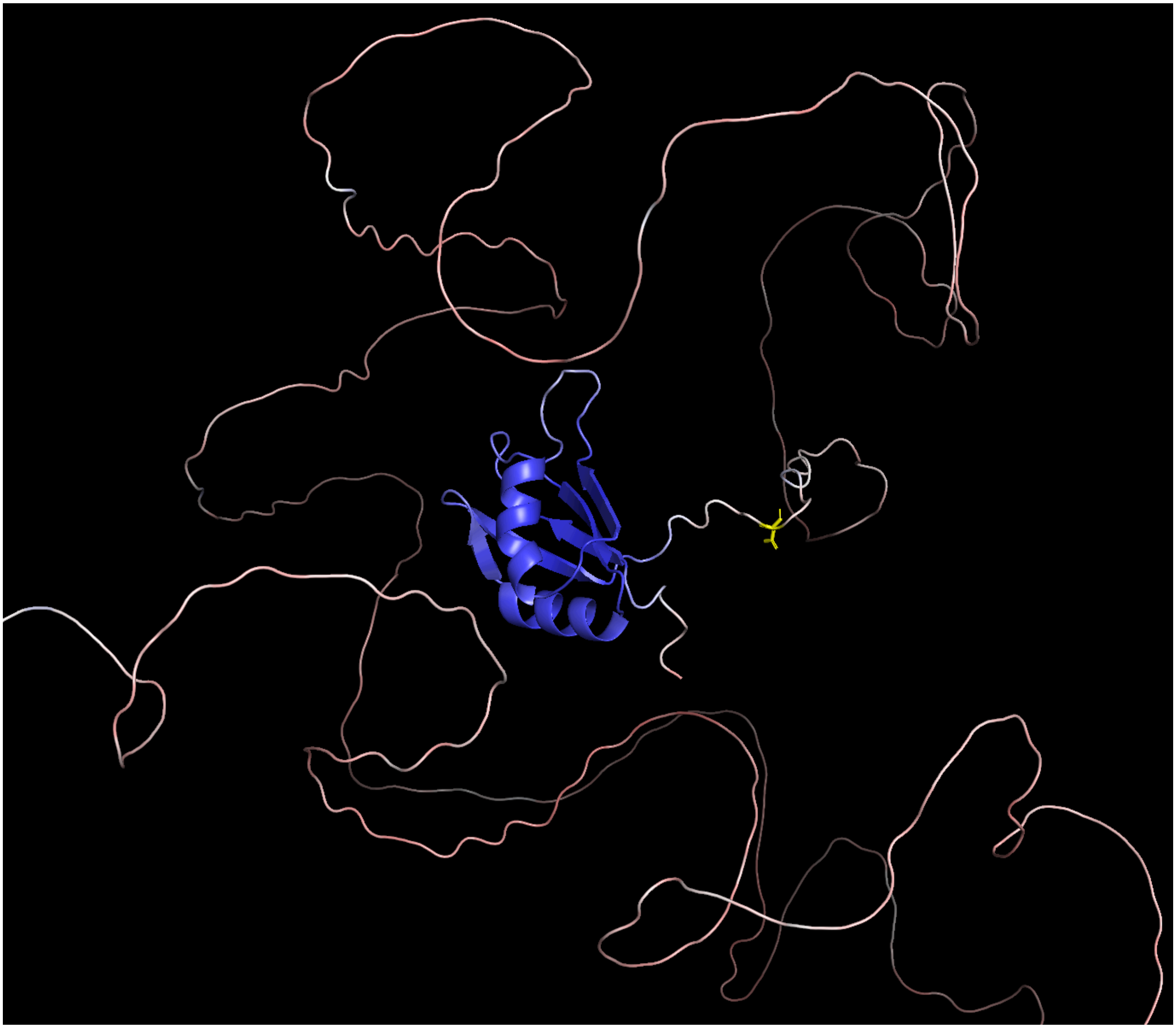

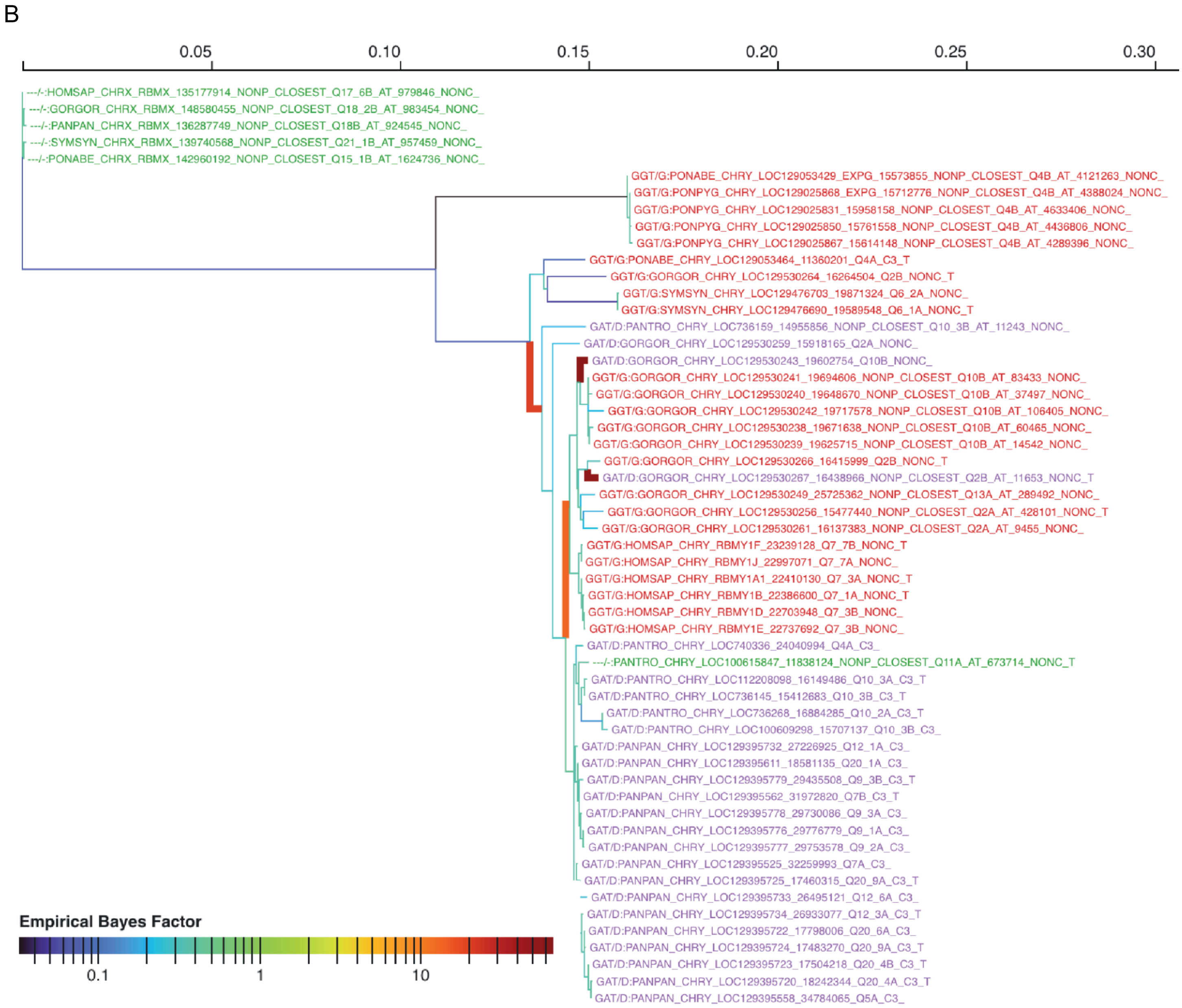
B
0.05
0.10
0.15
0.20
0.25
0.30
---/-:HOMSAP_CHRX_RBMX_135177914_NONP_CLOSEST_Q17_6B_AT_979846_NONC_
---/-:GORGOR_CHRX_RBMX_148580455_NONP_CLOSEST_Q18_2B_AT_983454_NONC_
---/-:PANPAN_CHRX_RBMX_136287749_NONP_CLOSEST_Q18B_AT_924545_NONC_
---/-:SYMSYN_CHRX_RBMX_139740568_NONP_CLOSEST_Q21_1B_AT_957459_NONC_
---/-:PONABE_CHRX_RBMX_142960192_NONP_CLOSEST_Q15_1B_AT_1624736_NONC_
GGT/G:PONABE_CHRY_LOC129053429_EXPG_15573855_NONP_CLOSEST_Q4B_AT_4121263_NONC_
GGT/G:PONPYG_CHRY_LOC129025868_EXPG_15712776_NONP_CLOSEST_Q4B_AT_4388024_NONC_
GGT/G:PONPYG_CHRY_LOC129025831_15958158_NONP_CLOSEST_Q4B_AT_4633406_NONC_
GGT/G:PONPYG_CHRY_LOC129025850_15761558_NONP_CLOSEST_Q4B_AT_4436806_NONC_
GGT/G:PONPYG_CHRY_LOC129025867_15614148_NONP_CLOSEST_Q4B_AT_4289396_NONC_
GGT/G:PONABE_CHRY_LOC129053464_11360201_Q4A_C3_T
GGT/G:GORGOR_CHRY_LOC129530264_16264504_Q2B_NONC_T
GGT/G:SYMSYN_CHRY_LOC129476703_19871324_Q6_2A_NONC_
GGT/G:SYMSYN_CHRY_LOC129476690_19589548_Q6_1A_NONC_T
GAT/D:PANTRO_CHRY_LOC736159_14955856_NONP_CLOSEST_Q10_3B_AT_11243_NONC_
GAT/D:GORGOR_CHRY_LOC129530259_15918165_Q2A_NONC_
GAT/D:GORGOR_CHRY_LOC129530243_19602754_Q10B_NONC_
GGT/G:GORGOR_CHRY_LOC129530241_19694606_NONP_CLOSEST_Q10B_AT_83433_NONC_
GGT/G:GORGOR_CHRY_LOC129530240_19648670_NONP_CLOSEST_Q10B_AT_37497_NONC_
GGT/G:GORGOR_CHRY_LOC129530242_19717578_NONP_CLOSEST_Q10B_AT_106405_NONC_
GGT/G:GORGOR_CHRY_LOC129530238_19671638_NONP_CLOSEST_Q10B_AT_60465_NONC_
GGT/G:GORGOR_CHRY_LOC129530239_19625715_NONP_CLOSEST_Q10B_AT_14542_NONC_
GGT/G:GORGOR_CHRY_LOC129530266_16415999_Q2B_NONC_T
GAT/D:GORGOR_CHRY_LOC129530267_16438966_NONP_CLOSEST_Q2B_AT_11653_NONC_T
GGT/G:GORGOR_CHRY_LOC129530249_25725362_NONP_CLOSEST_Q13A_AT_289492_NONC_
GGT/G:GORGOR_CHRY_LOC129530256_15477440_NONP_CLOSEST_Q2A_AT_428101_NONC_T
GGT/G:GORGOR_CHRY_LOC129530261_16137383_NONP_CLOSEST_Q2A_AT_9455_NONC_
GGT/G:HOMSAP_CHRY_RBMY1F_23239128_Q7_7B_NONC_T
GGT/G:HOMSAP_CHRY_RBMY1J_22997071_Q7_7A_NONC_
GGT/G:HOMSAP_CHRY_RBMY1A1_22410130_Q7_3A_NONC_T
GGT/G:HOMSAP_CHRY_RBMY1B_22386600_Q7_1A_NONC_T
GGT/G:HOMSAP_CHRY_RBMY1D_22703948_Q7_3B_NONC_
GGT/G:HOMSAP_CHRY_RBMY1E_22737692_Q7_3B_NONC_
GAT/D:PANTRO_CHRY_LOC740336_24040994_Q4A_C3_
---/-:PANTRO_CHRY_LOC100615847_11838124_NONP_CLOSEST_Q11A_AT_673714_NONC_T
GAT/D:PANTRO_CHRY_LOC112208098_16149486_Q10_3A_C3_T
GAT/D:PANTRO_CHRY_LOC736145_15412683_Q10_3B_C3_T
GAT/D:PANTRO_CHRY_LOC736268_16884285_Q10_2A_C3_T
GAT/D:PANTRO_CHRY_LOC100609298_15707137_Q10_3B_C3_
GAT/D:PANPAN_CHRY_LOC129395732_27226925_Q12_1A_C3_
GAT/D:PANPAN_CHRY_LOC129395611_18581135_Q20_1A_C3_
GAT/D:PANPAN_CHRY_LOC129395779_29435508_Q9_3B_C3_T
GAT/D:PANPAN_CHRY_LOC129395562_31972820_Q7B_C3_T
GAT/D:PANPAN_CHRY_LOC129395778_29730086_Q9_3A_C3_
GAT/D:PANPAN_CHRY_LOC129395776_29776779_Q9_1A_C3_
GAT/D:PANPAN_CHRY_LOC129395777_29753578_Q9_2A_C3_
GAT/D:PANPAN_CHRY_LOC129395525_32259993_Q7A_C3_
GAT/D:PANPAN_CHRY_LOC129395725_17460315_Q20_9A_C3_T
GAT/D:PANPAN_CHRY_LOC129395733_26495121_Q12_6A_C3_
GAT/D:PANPAN_CHRY_LOC129395734_26933077_Q12_3A_C3_T
GAT/D:PANPAN_CHRY_LOC129395722_17798006_Q20_6A_C3_
GAT/D:PANPAN_CHRY_LOC129395724_17483270_Q20_9A_C3_T
GAT/D:PANPAN_CHRY_LOC129395723_17504218_Q20_4B_C3_T
GAT/D:PANPAN_CHRY_LOC129395720_18242344_Q20_4A_C3_T
GAT/D:PANPAN_CHRY_LOC129395558_34784065_Q5A_C3_
Empirical Bayes Factor
0.1
1
10

### Figure S15. Episodic diversifying selection on individual sites of the *TSPY* residue 76

(**A**) Residue 76 in the cartoon representation of the folder protein in PyMol (Schrödinger, LLC 2026). (**B**) Four branches under diversifying selection at residue 76 are drawn with a thicker lime-green stroke.

A

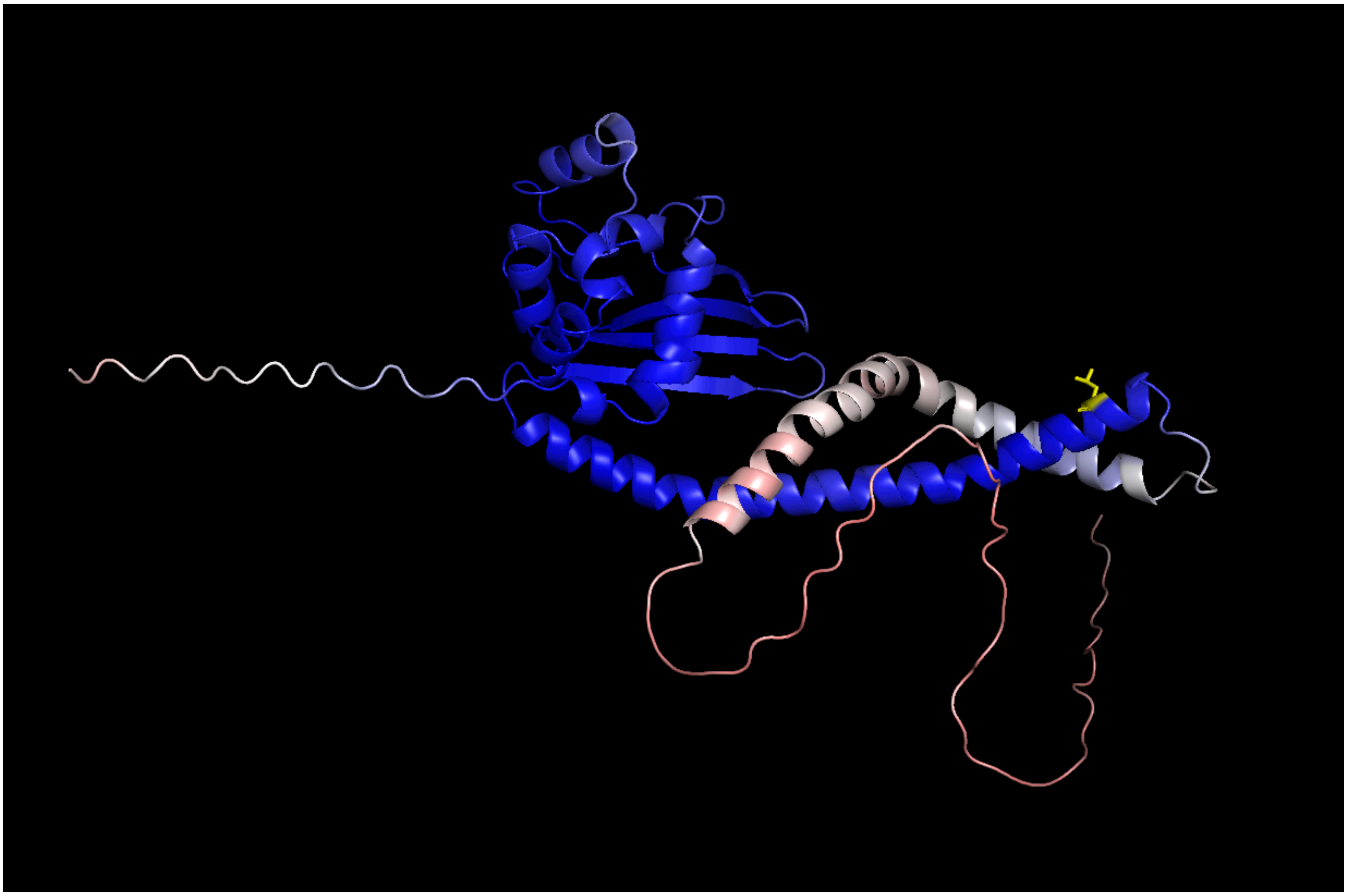

B

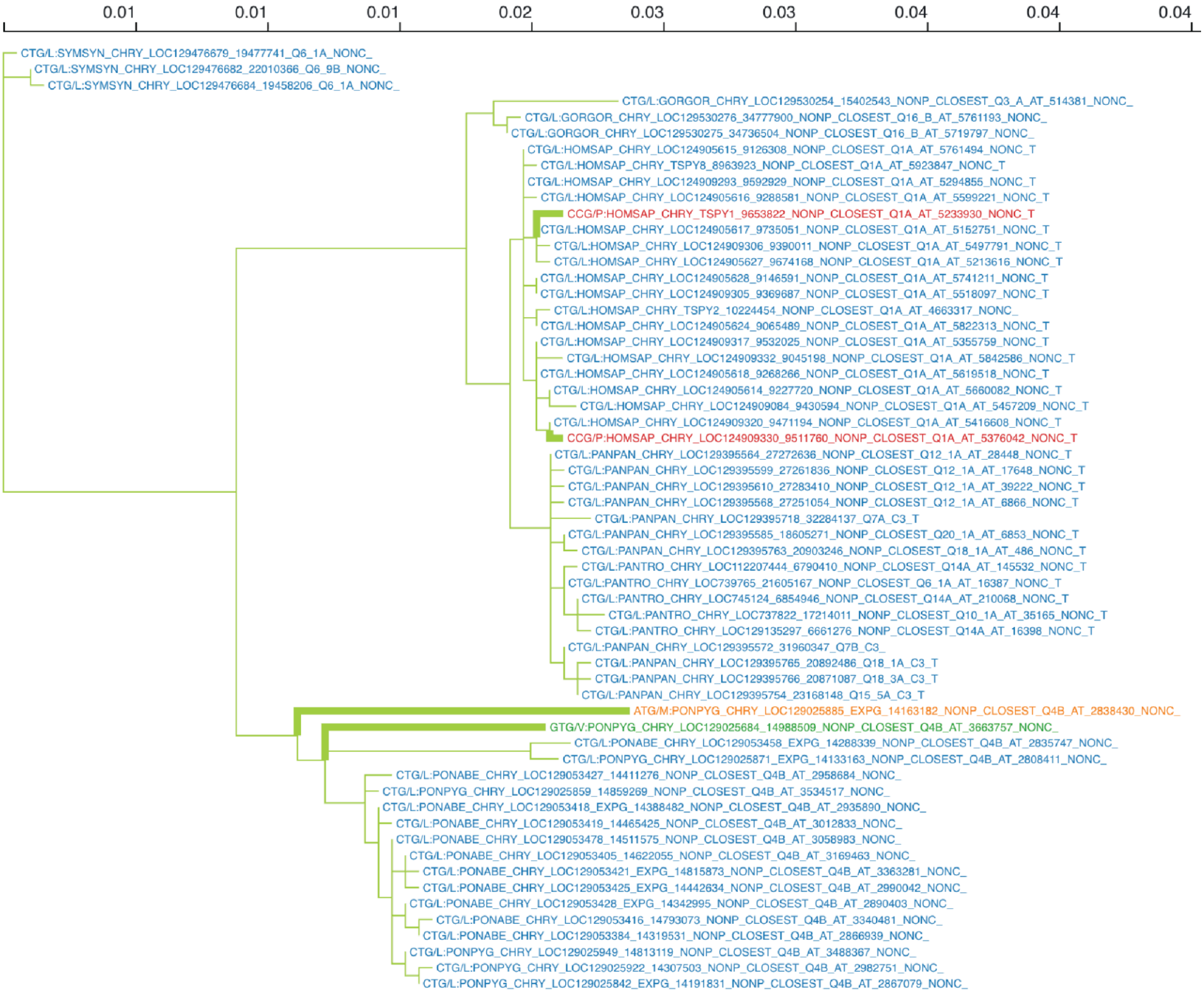
0.01
0.01
0.01
0.02
0.03
0.03
0.04
0.04
0.04
CTG/L:SYMSYN_CHRY_LOC129476679_19477741_Q6_1A_NONC_
CTG/L:SYMSYN_CHRY_LOC129476682_22010366_Q6_9B_NONC_
CTG/L:SYMSYN_CHRY_LOC129476684_19458206_Q6_1A_NONC_
CTG/L:GORGOR_CHRY_LOC129530254_15402543_NONP_CLOSEST_Q3_A_AT_514381_NONC_
CTG/L:GORGOR_CHRY_LOC129530276_34777900_NONP_CLOSEST_Q16_B_AT_5761193_NONC_
CTG/L:GORGOR_CHRY_LOC129530275_34736504_NONP_CLOSEST_Q16_B_AT_5719797_NONC_
CTG/L:HOMSAP_CHRY_LOC124905615_9126308_NONP_CLOSEST_Q1A_AT_5761494_NONC_T
CTG/L:HOMSAP_CHRY_TSPY8_8963923_NONP_CLOSEST_Q1A_AT_5923847_NONC_T
CTG/L:HOMSAP_CHRY_LOC124909293_9592929_NONP_CLOSEST_Q1A_AT_5294855_NONC_T
CTG/L:HOMSAP_CHRY_LOC124905616_9288581_NONP_CLOSEST_Q1A_AT_5599221_NONC_T
CCG/P:HOMSAP_CHRY_TSPY1_9653822_NONP_CLOSEST_Q1A_AT_5233930_NONC_T
CTG/L:HOMSAP_CHRY_LOC124905617_9735051_NONP_CLOSEST_Q1A_AT_5152751_NONC_T
CTG/L:HOMSAP_CHRY_LOC124909306_9390011_NONP_CLOSEST_Q1A_AT_5497791_NONC_T
CTG/L:HOMSAP_CHRY_LOC124905627_9674168_NONP_CLOSEST_Q1A_AT_5213616_NONC_T
CTG/L:HOMSAP_CHRY_LOC124905628_9146591_NONP_CLOSEST_Q1A_AT_5741211_NONC_T
CTG/L:HOMSAP_CHRY_LOC124909305_9369687_NONP_CLOSEST_Q1A_AT_5518097_NONC_T
CTG/L:HOMSAP_CHRY_TSPY2_10224454_NONP_CLOSEST_Q1A_AT_4663317_NONC_
CTG/L:HOMSAP_CHRY_LOC124905624_9065489_NONP_CLOSEST_Q1A_AT_5822313_NONC_T
CTG/L:HOMSAP_CHRY_LOC124909317_9532025_NONP_CLOSEST_Q1A_AT_5355759_NONC_T
CTG/L:HOMSAP_CHRY_LOC124909332_9045198_NONP_CLOSEST_Q1A_AT_5842586_NONC_T
CTG/L:HOMSAP_CHRY_LOC124905618_9268266_NONP_CLOSEST_Q1A_AT_5619518_NONC_T
CTG/L:HOMSAP_CHRY_LOC124905614_9227720_NONP_CLOSEST_Q1A_AT_5660082_NONC_T
CTG/L:HOMSAP_CHRY_LOC124909084_9430594_NONP_CLOSEST_Q1A_AT_5457209_NONC_T
CTG/L:HOMSAP_CHRY_LOC124909320_9471194_NONP_CLOSEST_Q1A_AT_5416608_NONC_T
CCG/P:HOMSAP_CHRY_LOC124909330_9511760_NONP_CLOSEST_Q1A_AT_5376042_NONC_T
CTG/L:PANPAN_CHRY_LOC129395564_27272636_NONP_CLOSEST_Q12_1A_AT_28448_NONC_T
CTG/L:PANPAN_CHRY_LOC129395599_27261836_NONP_CLOSEST_Q12_1A_AT_17648_NONC_T
CTG/L:PANPAN_CHRY_LOC129395610_27283410_NONP_CLOSEST_Q12_1A_AT_39222_NONC_T
CTG/L:PANPAN_CHRY_LOC129395568_27251054_NONP_CLOSEST_Q12_1A_AT_6866_NONC_T
CTG/L:PANPAN_CHRY_LOC129395718_32284137_Q7A_C3_T
CTG/L:PANPAN_CHRY_LOC129395585_18605271_NONP_CLOSEST_Q20_1A_AT_6853_NONC_T
CTG/L:PANPAN_CHRY_LOC129395763_20903246_NONP_CLOSEST_Q18_1A_AT_486_NONC_T
CTG/L:PANTRO_CHRY_LOC112207444_6790410_NONP_CLOSEST_Q14A_AT_145532_NONC_T
CTG/L:PANTRO_CHRY_LOC739765_21605167_NONP_CLOSEST_Q6_1A_AT_16387_NONC_T
CTG/L:PANTRO_CHRY_LOC745124_6854946_NONP_CLOSEST_Q14A_AT_210068_NONC_T
CTG/L:PANTRO_CHRY_LOC737822_17214011_NONP_CLOSEST_Q10_1A_AT_35165_NONC_T
CTG/L:PANTRO_CHRY_LOC129135297_6661276_NONP_CLOSEST_Q14A_AT_16398_NONC_T
CTG/L:PANPAN_CHRY_LOC129395572_31960347_Q7B_C3_
CTG/L:PANPAN_CHRY_LOC129395765_20892486_Q18_1A_C3_T
CTG/L:PANPAN_CHRY_LOC129395766_20871087_Q18_3A_C3_T
CTG/L:PANPAN_CHRY_LOC129395754_23168148_Q15_5A_C3_T
ATG/M:PONPYG_CHRY_LOC129025885_EXPG_14163182_NONP_CLOSEST_Q4B_AT_2838430_NONC_
GTG/V:PONPYG_CHRY_LOC129025684_14988509_NONP_CLOSEST_Q4B_AT_3663757_NONC_
CTG/L:PONABE_CHRY_LOC129053458_EXPG_14288339_NONP_CLOSEST_Q4B_AT_2835747_NONC_
CTG/L:PONPYG_CHRY_LOC129025871_EXPG_14133163_NONP_CLOSEST_Q4B_AT_2808411_NONC_
CTG/L:PONABE_CHRY_LOC129053427_14411276_NONP_CLOSEST_Q4B_AT_2958684_NONC_
CTG/L:PONPYG_CHRY_LOC129025859_14859269_NONP_CLOSEST_Q4B_AT_3534517_NONC_
CTG/L:PONABE_CHRY_LOC129053418_EXPG_14388482_NONP_CLOSEST_Q4B_AT_2935890_NONC_
CTG/L:PONABE_CHRY_LOC129053419_14465425_NONP_CLOSEST_Q4B_AT_3012833_NONC_
CTG/L:PONABE_CHRY_LOC129053478_14511575_NONP_CLOSEST_Q4B_AT_3058983_NONC_
CTG/L:PONABE_CHRY_LOC129053405_14622055_NONP_CLOSEST_Q4B_AT_3169463_NONC_
CTG/L:PONABE_CHRY_LOC129053421_EXPG_14815873_NONP_CLOSEST_Q4B_AT_3363281_NONC_
CTG/L:PONABE_CHRY_LOC129053425_EXPG_14442634_NONP_CLOSEST_Q4B_AT_2990042_NONC_
CTG/L:PONABE_CHRY_LOC129053428_EXPG_14342995_NONP_CLOSEST_Q4B_AT_2890403_NONC_
CTG/L:PONABE_CHRY_LOC129053416_14793073_NONP_CLOSEST_Q4B_AT_3340481_NONC_
CTG/L:PONABE_CHRY_LOC129053384_14319531_NONP_CLOSEST_Q4B_AT_2866939_NONC_
CTG/L:PONPYG_CHRY_LOC129025949_14813119_NONP_CLOSEST_Q4B_AT_3488367_NONC_
CTG/L:PONPYG_CHRY_LOC129025922_14307503_NONP_CLOSEST_Q4B_AT_2982751_NONC_
CTG/L:PONPYG_CHRY_LOC129025842_EXPG_14191831_NONP_CLOSEST_Q4B_AT_2867079_NONC_

### Figure S16. Episodic diversifying selection on residue 475 of *CDY*

(**A**) Residue 475 in the cartoon representation of the folder protein in PyMol (Schrödinger, LLC 2026). (**B**) Two branches under diversifying selection at residue 475 drawn with a thicker (lime green and dark red) stroke.

A

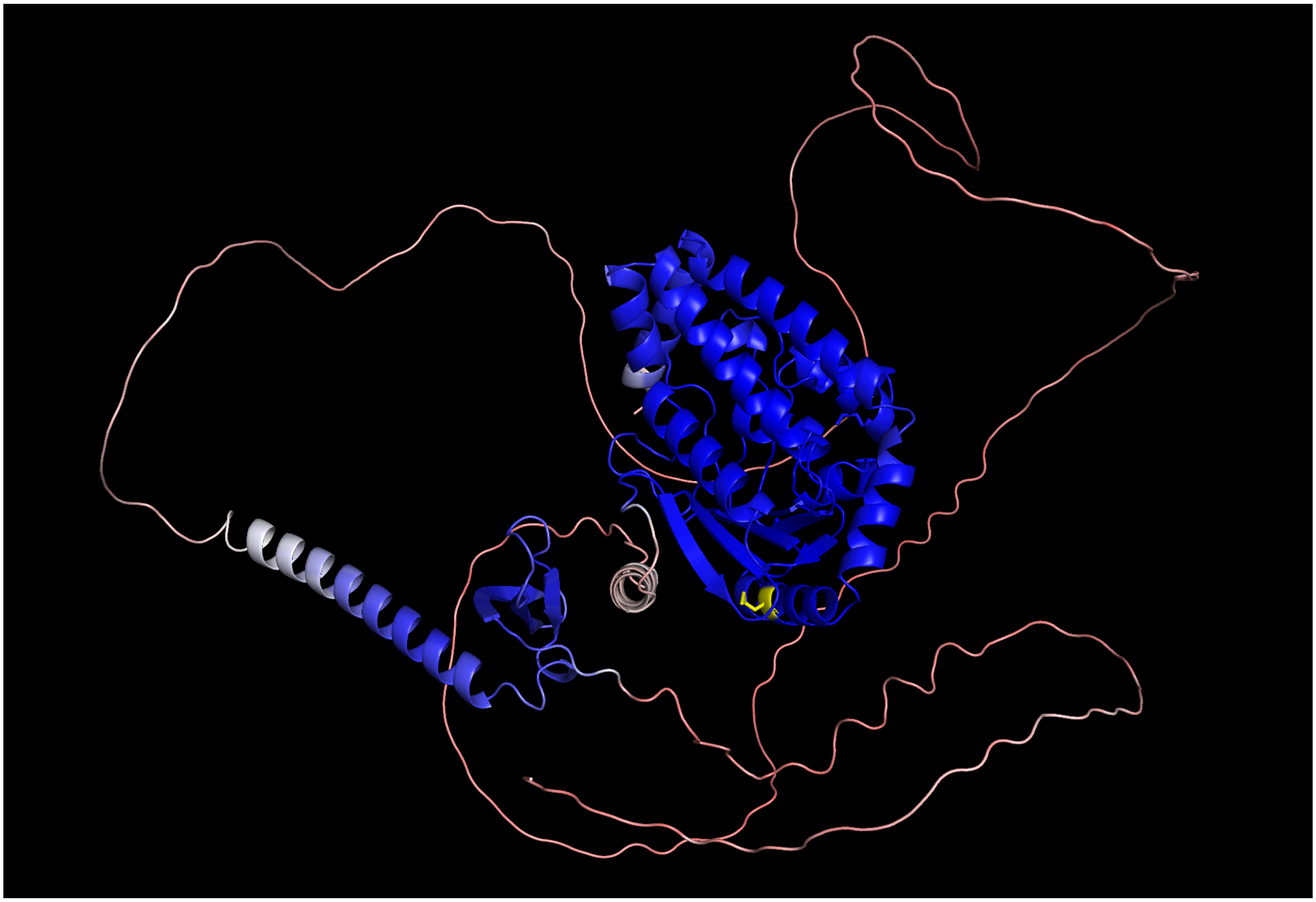

B

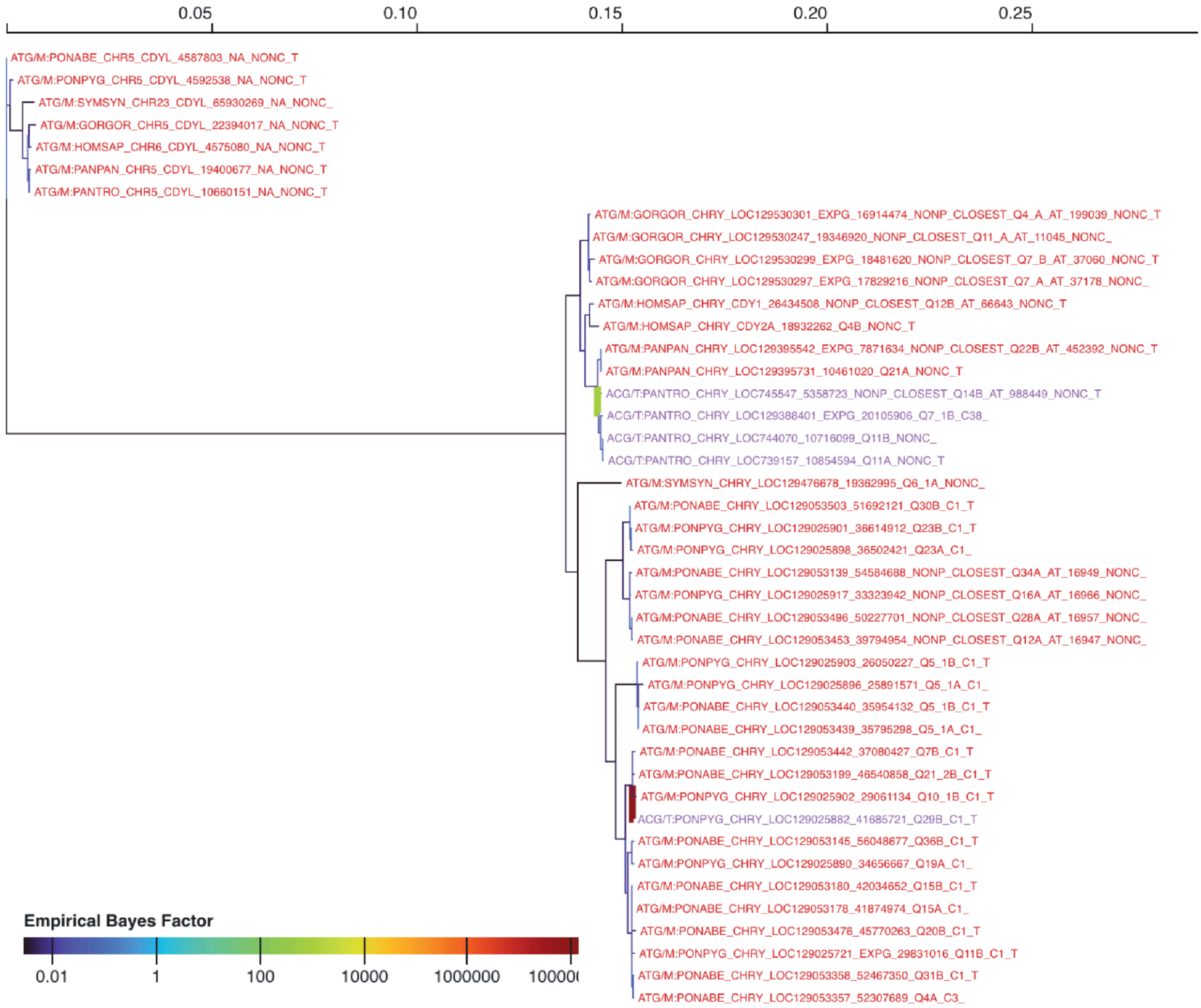

0.05
0.10
0.15
0.20
0.25
ATG/M:PONABE_CHR5_CDYL_4587803_NA_NONC_T
ATG/M:PONPYG_CHR5_CDYL_4592538_NA_NONC_T
ATG/M:SYMSYN_CHR23_CDYL_65930269_NA_NONC_
ATG/M:GORGOR_CHR5_CDYL_22394017_NA_NONC_T
ATG/M:HOMSAP_CHR6_CDYL_4575080_NA_NONC_T
ATG/M:PANPAN_CHR5_CDYL_19400677_NA_NONC_T
ATG/M:PANTRO_CHR5_CDYL_10660151_NA_NONC_T
ATG/M:GORGOR_CHRY_LOC129530301_EXPG_16914474_NONP_CLOSEST_Q4_A_AT_199039_NONC_T
ATG/M:GORGOR_CHRY_LOC129530247_19346920_NONP_CLOSEST_Q11_A_AT_11045_NONC_
ATG/M:GORGOR_CHRY_LOC129530299_EXPG_18481620_NONP_CLOSEST_Q7_B_AT_37060_NONC_T
ATG/M:GORGOR_CHRY_LOC129530297_EXPG_17829216_NONP_CLOSEST_Q7_A_AT_37178_NONC_
ATG/M:HOMSAP_CHRY_CDY1_26434508_NONP_CLOSEST_Q12B_AT_66643_NONC_T
ATG/M:HOMSAP_CHRY_CDY2A_18932262_Q4B_NONC_T
ATG/M:PANPAN_CHRY_LOC129395542_EXPG_7871634_NONP_CLOSEST_Q22B_AT_452392_NONC_T
ATG/M:PANPAN_CHRY_LOC129395731_10461020_Q21A_NONC_T
ACG/T:PANTRO_CHRY_LOC745547_5358723_NONP_CLOSEST_Q14B_AT_988449_NONC_T
ACG/T:PANTRO_CHRY_LOC129388401_EXPG_20105906_Q7_1B_C38_
ACG/T:PANTRO_CHRY_LOC744070_10716099_Q11B_NONC_
ACG/T:PANTRO_CHRY_LOC739157_10854594_Q11A_NONC_T
ATG/M:SYMSYN_CHRY_LOC129476678_19362995_Q6_1A_NONC_
ATG/M:PONABE_CHRY_LOC129053503_51692121_Q30B_C1_T
ATG/M:PONPYG_CHRY_LOC129025901_36614912_Q23B_C1_T
ATG/M:PONPYG_CHRY_LOC129025898_36502421_Q23A_C1_
ATG/M:PONABE_CHRY_LOC129053139_54584688_NONP_CLOSEST_Q34A_AT_16949_NONC_
ATG/M:PONPYG_CHRY_LOC129025917_33323942_NONP_CLOSEST_Q16A_AT_16966_NONC_
ATG/M:PONABE_CHRY_LOC129053496_50227701_NONP_CLOSEST_Q28A_AT_16957_NONC_
ATG/M:PONABE_CHRY_LOC129053453_39794954_NONP_CLOSEST_Q12A_AT_16947_NONC_
ATG/M:PONPYG_CHRY_LOC129025903_26050227_Q5_1B_C1_T
ATG/M:PONPYG_CHRY_LOC129025896_25891571_Q5_1A_C1_
ATG/M:PONABE_CHRY_LOC129053440_35954132_Q5_1B_C1_T
ATG/M:PONABE_CHRY_LOC129053439_35795298_Q5_1A_C1_
ATG/M:PONABE_CHRY_LOC129053442_37080427_Q7B_C1_T
ATG/M:PONABE_CHRY_LOC129053199_46540858_Q21_2B_C1_T
ATG/M:PONPYG_CHRY_LOC129025902_29061134_Q10_1B_C1_T
ACG/T:PONPYG_CHRY_LOC129025882_41685721_Q29B_C1_T
ATG/M:PONABE_CHRY_LOC129053145_56048677_Q36B_C1_T
ATG/M:PONPYG_CHRY_LOC129025890_34656667_Q19A_C1_
ATG/M:PONABE_CHRY_LOC129053180_42034652_Q15B_C1_T
ATG/M:PONABE_CHRY_LOC129053178_41874974_Q15A_C1_
ATG/M:PONABE_CHRY_LOC129053476_45770263_Q20B_C1_T
ATG/M:PONPYG_CHRY_LOC129025721_EXPG_29831016_Q11B_C1_T
ATG/M:PONABE_CHRY_LOC129053358_52467350_Q31B_C1_T
ATG/M:PONABE_CHRY_LOC129053357_52307689_Q4A_C3_
Empirical Bayes Factor
0.01
1
100
10000
1000000
100000

### Figure S17. Anomaly detected by the BUSTED-E test

(**A**) Branch leading to possible error affecting single sequence highlighted in burgundy. (**B**) Multisequence alignment with possible detected error highlighted in a red rectangle.

A

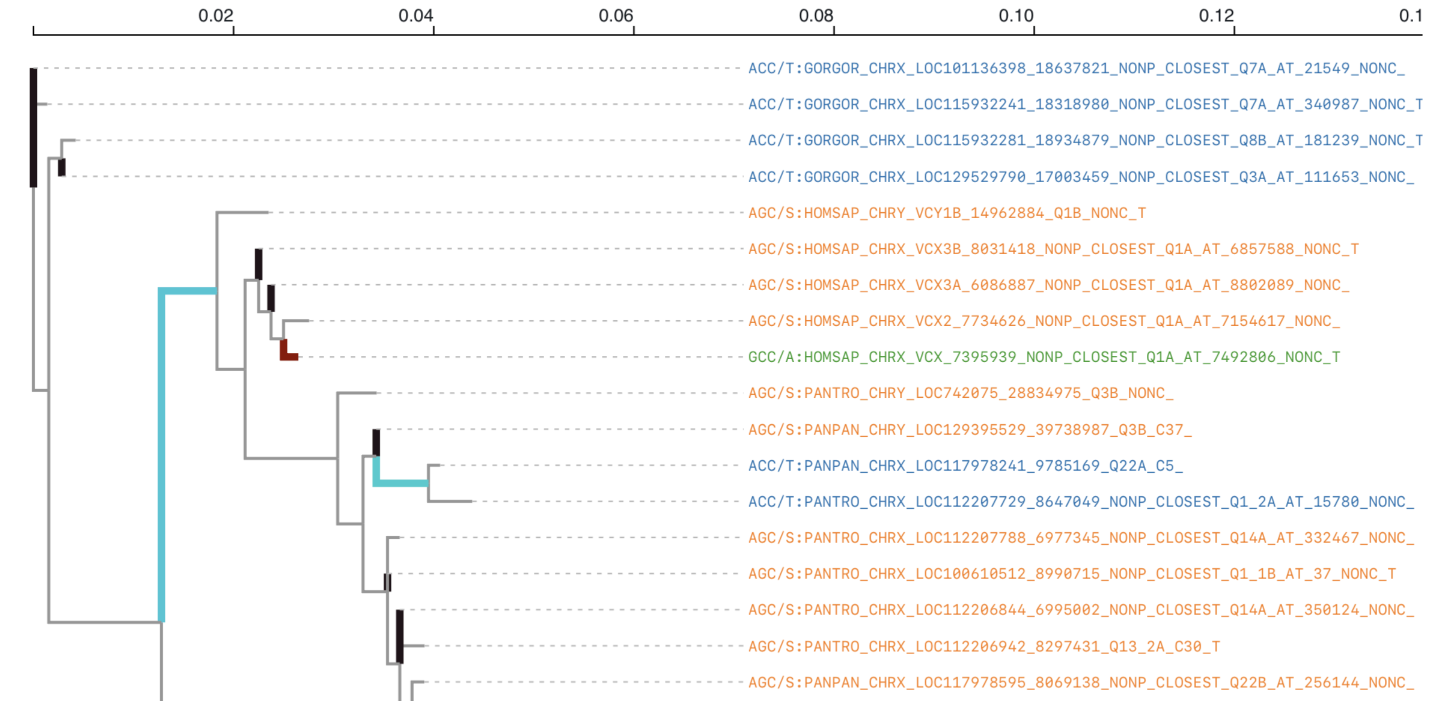

B

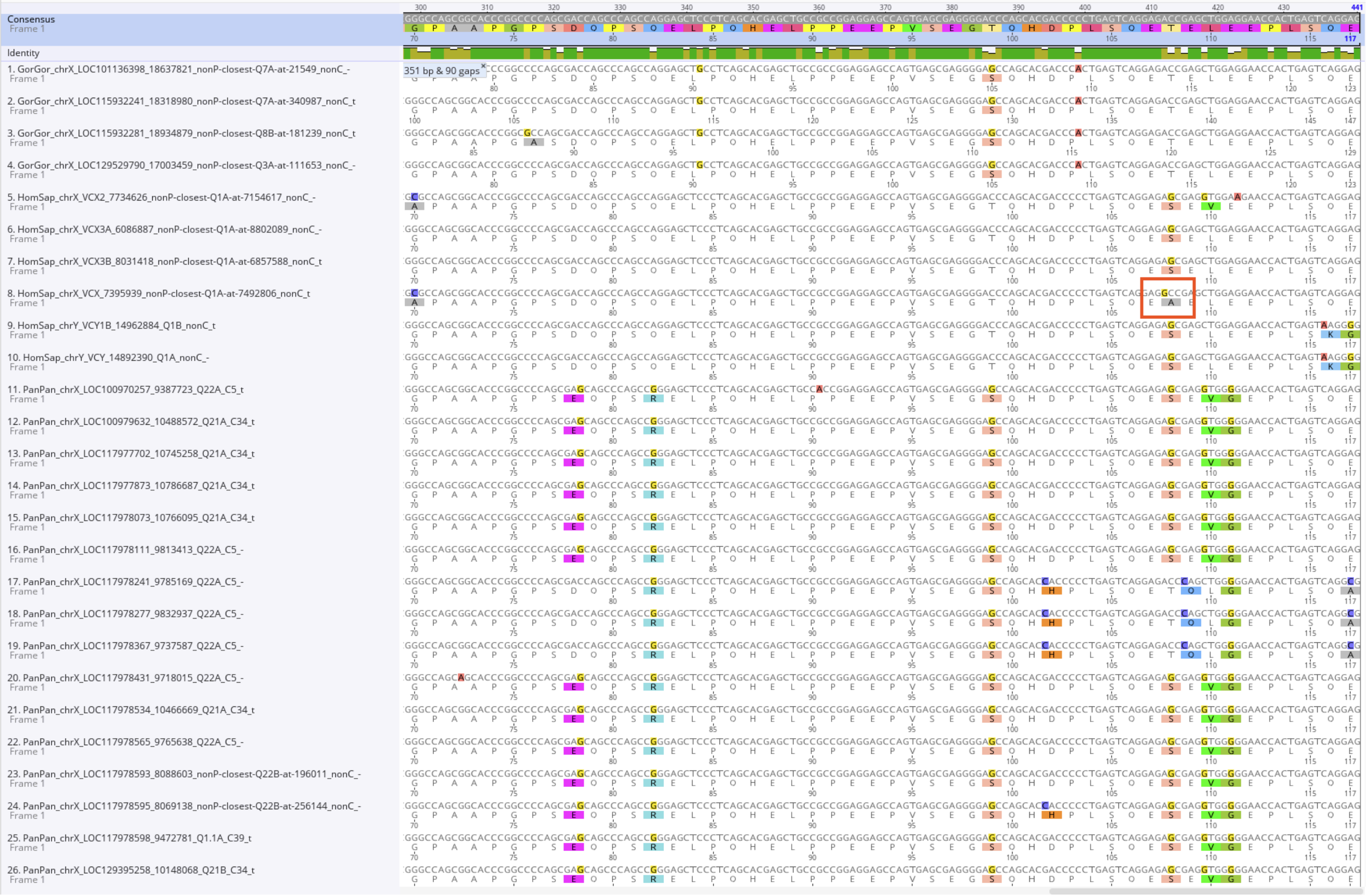

### Figure S18. The phylogenetic relationship among *RBMX*/*RBMY* copies

Sequences were aligned with `MAFFT` (Katoh and Standley 2013), and trees were constructed with the neighbor-joining method (Saitou and Nei 1987) with Jukes-Cantor (Jukes and Cantor 1969) correction. Trees were visualized with iTOL (Letunic and Bork 2024). Visualization excludes XP_063457564.1::LOC134729763::chrX::PanPan, XP_054400209.1::LOC100937012::chrX::PonAbe, and XP_054327929.1::LOC129024757::chrX::PonPyg due to their long branches (beyond plot resolution).

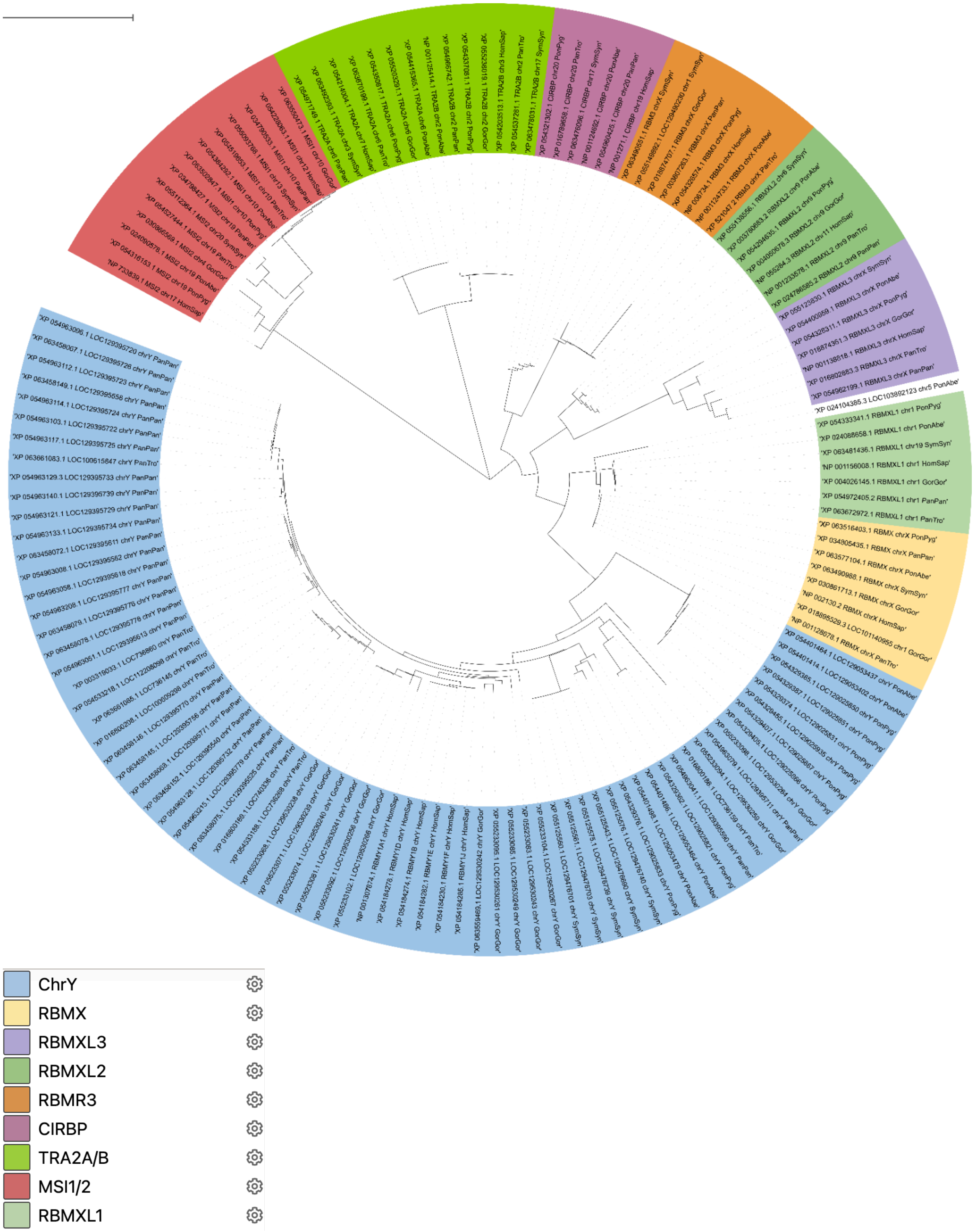

### Figure S19. Orangutan genus Y chromosomes, repeat, and selected gene content

(**A**) `MODDOTPLOT (Sweeten, Schatz, and Phillippy 2024)` visualization of the Bornean orangutan Y chromosome with the following genes highlighted: *DAZ* (green), *CDY* (yellow), *RBMY* (red), *TSPY* (grey), *HSFY* (cyan). (**B**) `MODDOTPLOT` (Sweeten, Schatz, and Phillippy 2024) visualization of the Sumatran orangutan Y chromosome. (**C**) `MODDOTPLOT` (Sweeten, Schatz, and Phillippy 2024) in comparative view with Bornean orangutan on the X-axis, and Sumatran orangutan on the Y-axis.

A

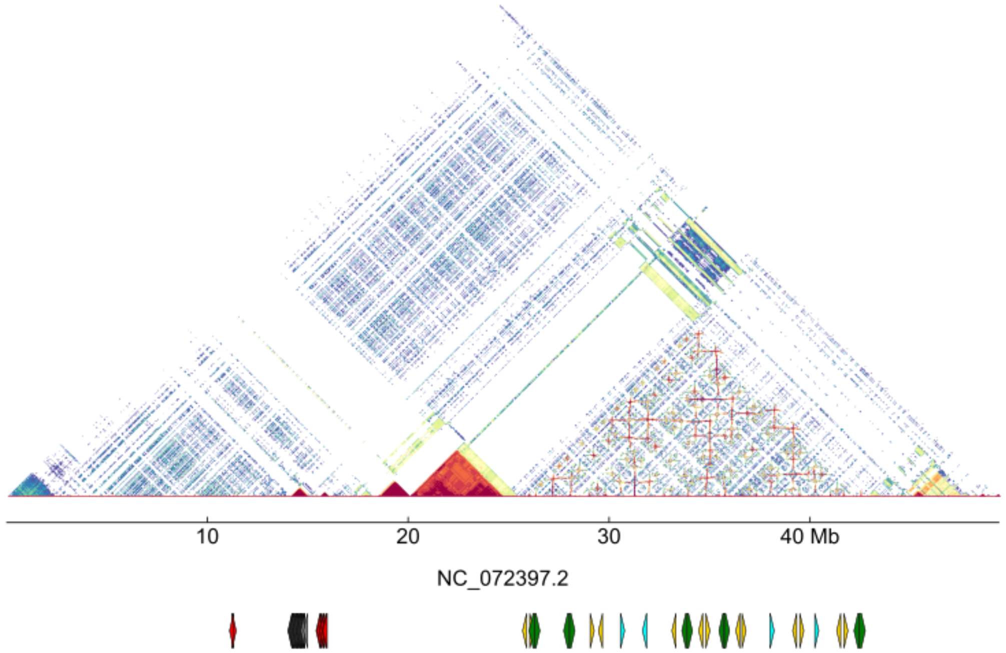

B

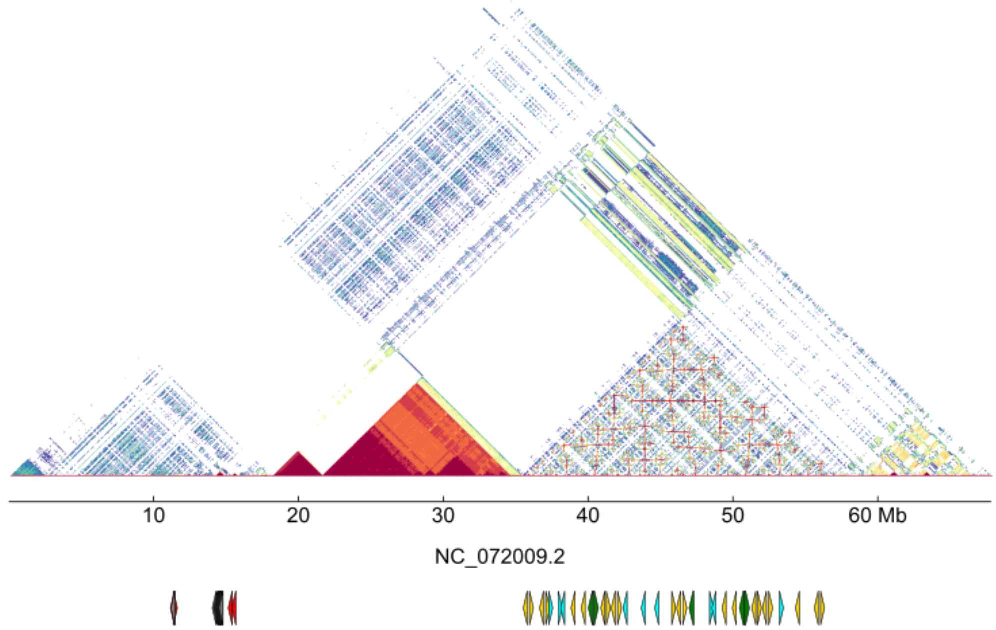

C

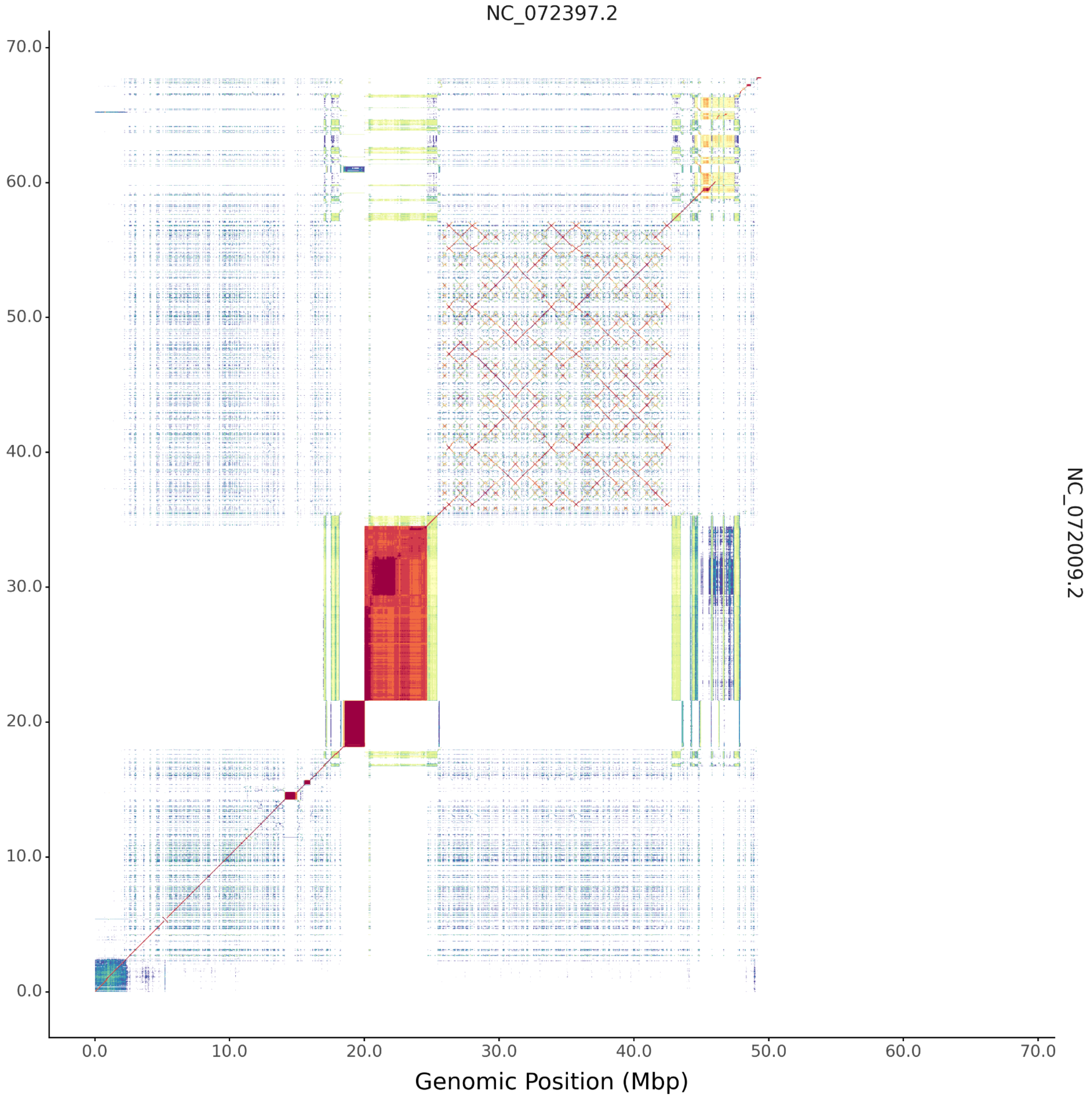

NC_072397.2
70.0
60.0
50.0
40.0
30.0
20.0
10.0
0.0
0.0
10.0
20.0
30.0
40.0
50.0
60.0
70.0
Genomic Position (Mbp)
NC_072009.2

Figure S20. Percentage identity between RRM domains of two palindromic *DAZ* copies of Bornean orangutan.

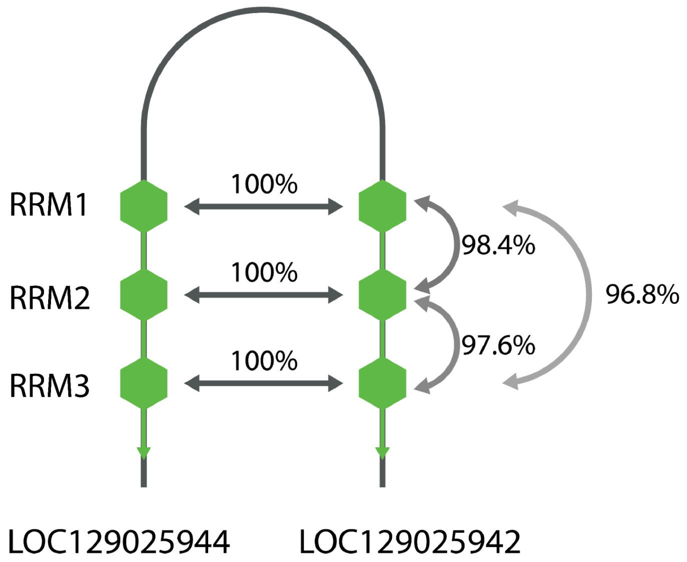

### Figure S21. *Pan* genus Y chromosomes, repeat, and selected gene content

(**A**) `MODDOTPLOT` (Sweeten, Schatz, and Phillippy 2024) visualization of the bonobo Y chromosome with the following genes highlighted: *DAZ* (green), *CDY* (yellow), *RBMY* (red), *TSPY* (grey). (**B**) `MODDOTPLOT` (Sweeten, Schatz, and Phillippy 2024) visualization of the chimpanzee Y chromosome. (**C**) `MODDOTPLOT` (Sweeten, Schatz, and Phillippy 2024) in comparative view with bonobo on the X-axis, and chimpanzee on the Y-axis.

A

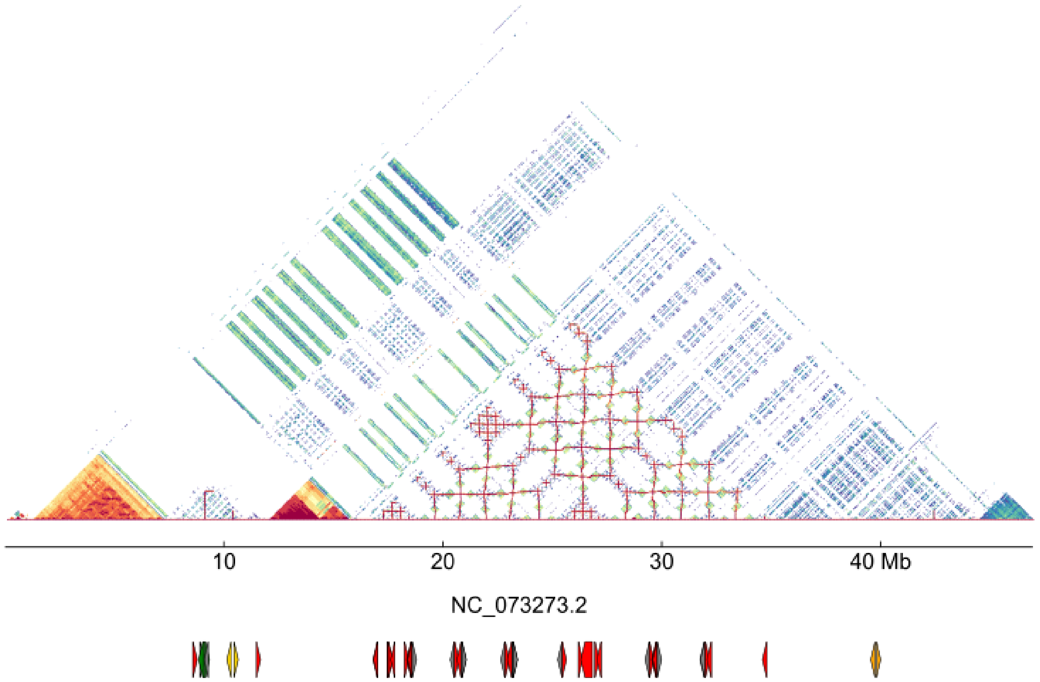

B

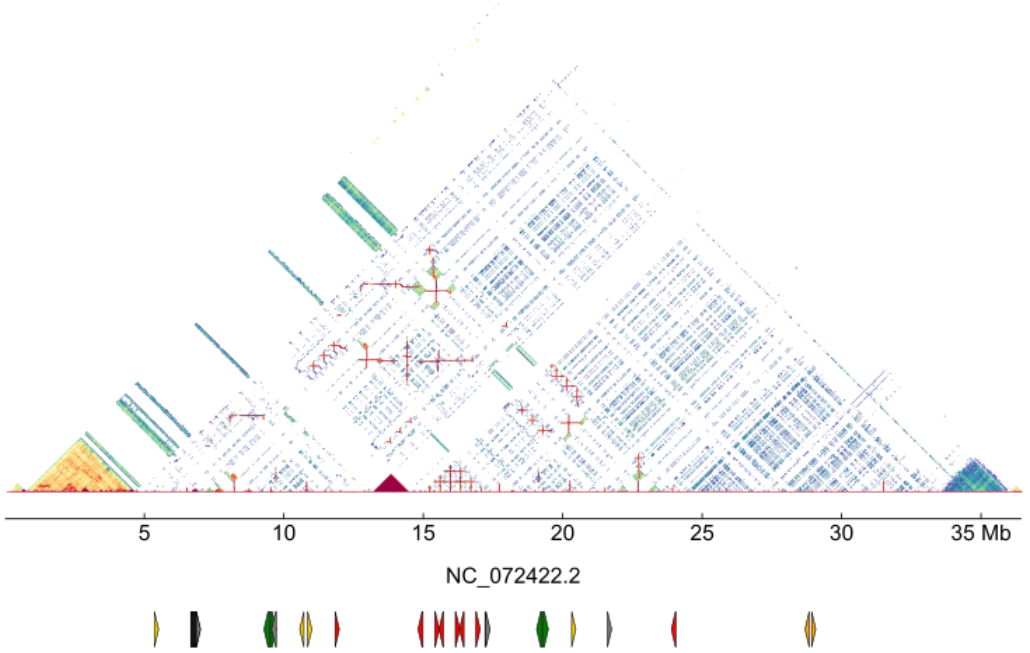

C

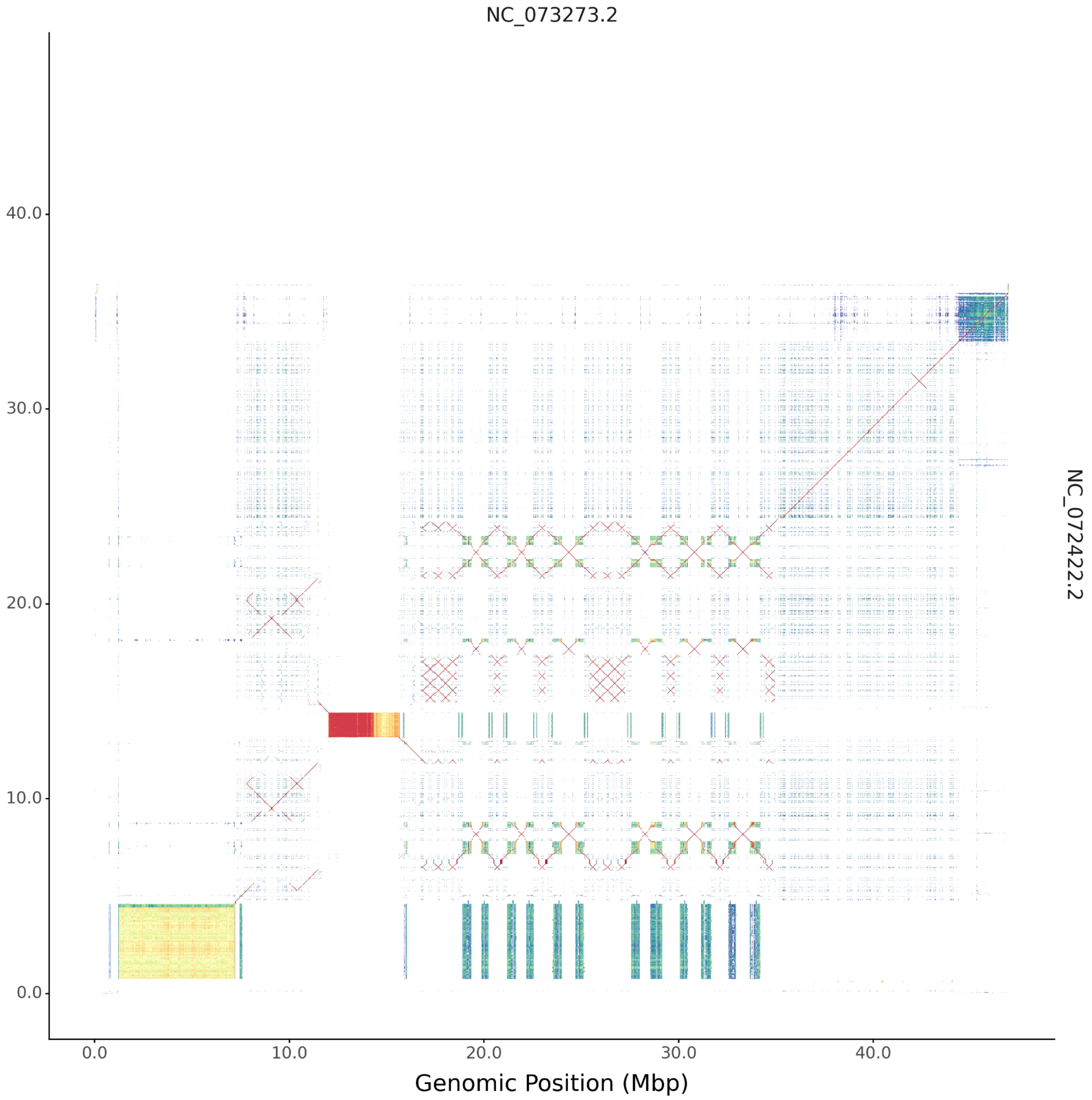

NC_073273.2
NC_072422.2
40.0
30.0
20.0
10.0
0.0
0.0
10.0
20.0
30.0
40.0
Genomic Position (Mbp)

### Figure S22. Bonobo-specific clades of the *RBMY* gene family

(**A**) A fragment of the *RBMY* gene family phylogenetic tree with only the bonobo copies and one additional chimpanzee *RBMY* copy used for tree rooting. Visualized in `ITOL` (Letunic and Bork 2024). Nodes from clade A marked in blue rectangle, and nodes from clade B marked in red. (**B**) Distribution of clade-specific alleles (Alleles that are at least 80% clade specific). (**C**) Distribution of *RBMY* gene copies. Clade A in blue, clade B marked in red.

A

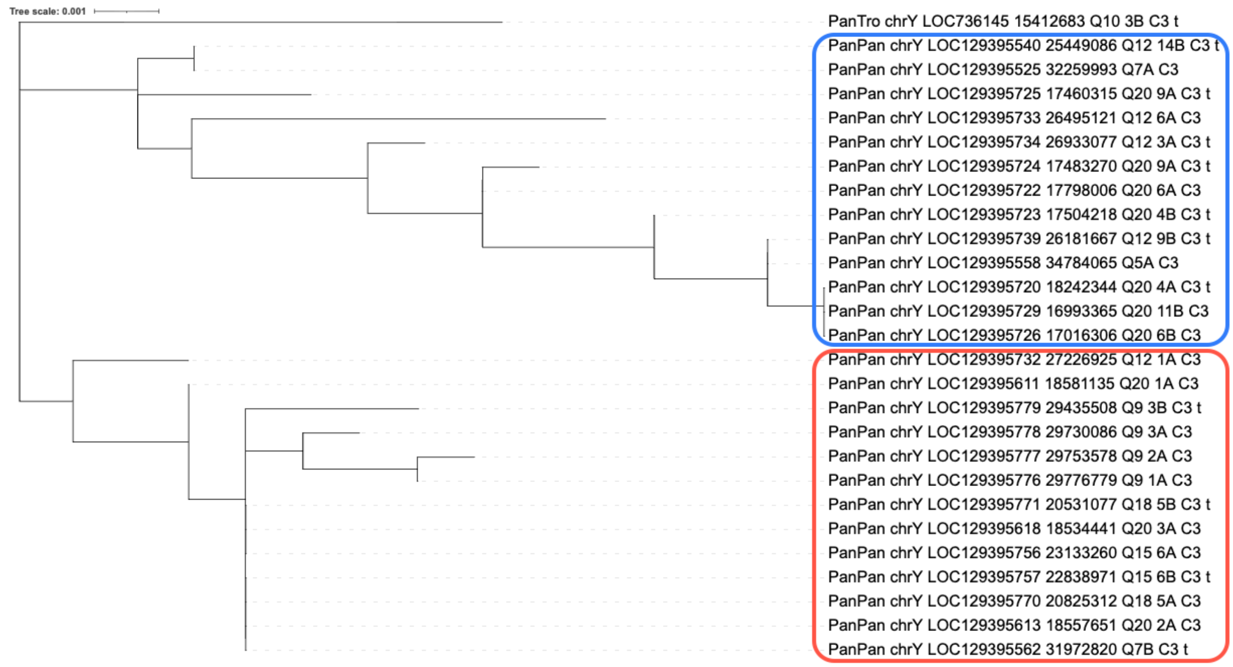

B

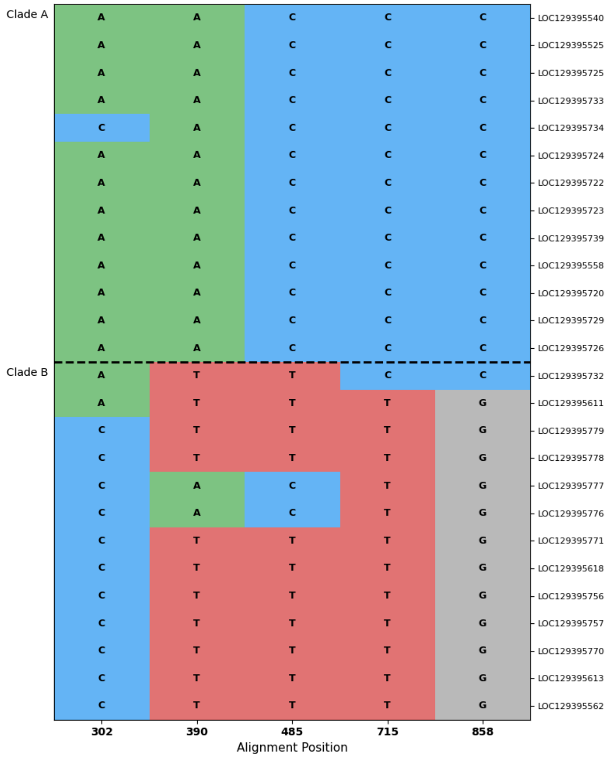

C

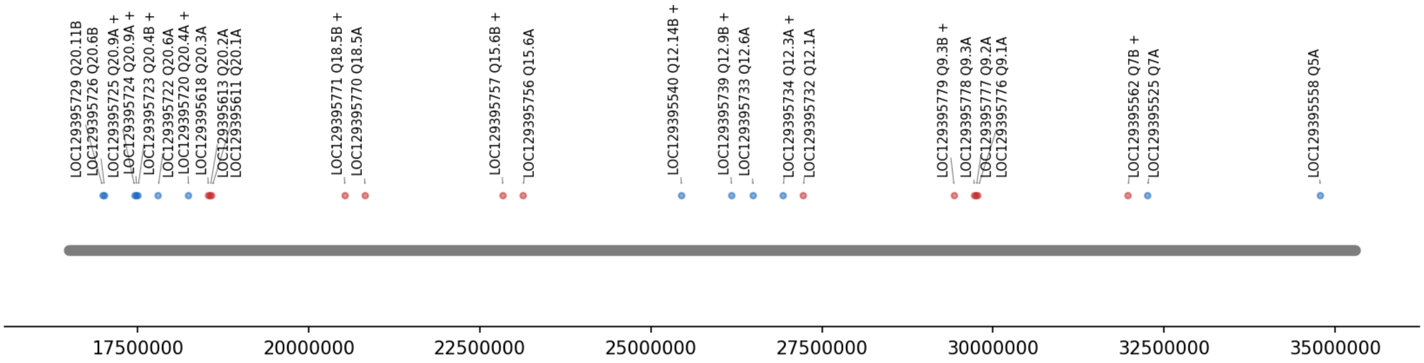
LOC129395729 Q20.11B
LOC129395726 Q20.6B
LOC129395725 Q20.9A +
LOC129395724 Q20.9A +
LOC129395723 Q20.4B +
LOC129395722 Q20.6A
LOC129395720 Q20.4A +
LOC129395618 Q20.3A
LOC129395613 Q20.2A
LOC129395611 Q20.1A
LOC129395771 Q18.5B +
LOC129395770 Q18.5A
LOC129395757 Q15.6B +
LOC129395756 Q15.6A
LOC129395540 Q12.14B +
LOC129395739 Q12.9B +
LOC129395733 Q12.6A
LOC129395734 Q12.3A +
LOC129395732 Q12.1A
LOC129395779 Q9.3B +
LOC129395778 Q9.3A
LOC129395777 Q9.2A
LOC129395776 Q9.1A
LOC129395562 Q7B +
LOC129395525 Q7A
LOC129395558 Q5A
17500000
20000000
22500000
25000000
27500000
30000000
32500000
35000000

### Figure S23. Palindrome containing two *RBMY* copies and an unbalanced *TSPY* array in bonobo

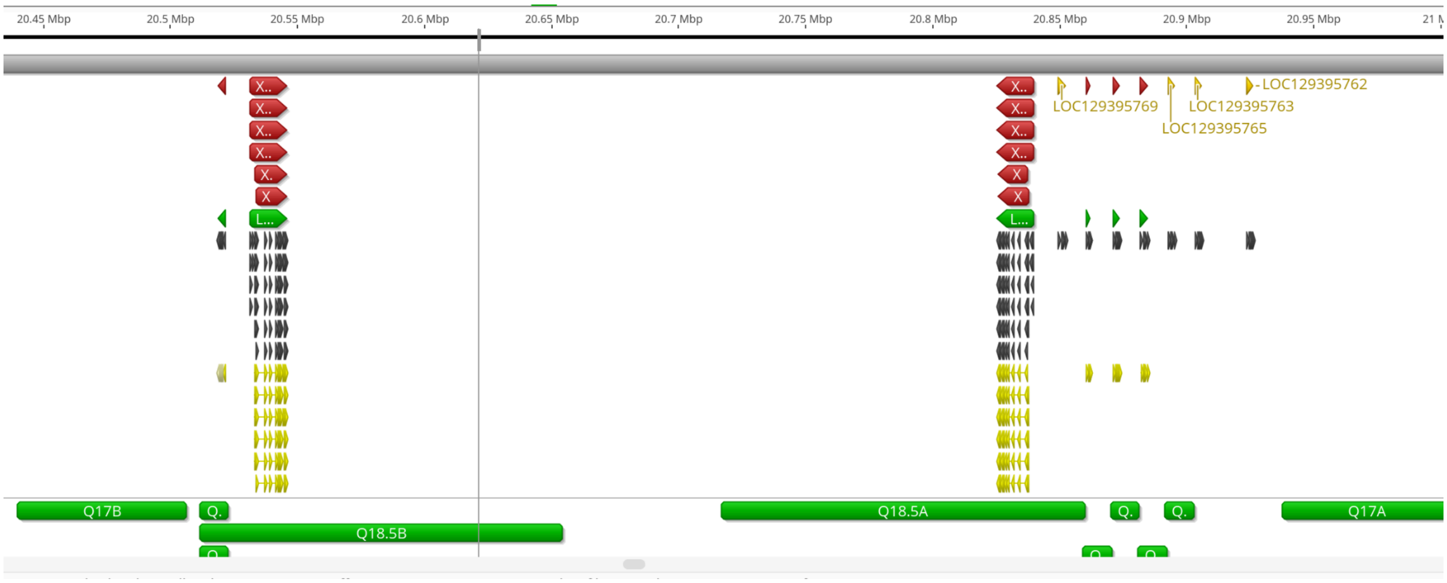

### Figure S24. Branch-specific diversifying selection in *RBMY*

Branch-specific diversifying selection identified by aBSREL (Smith et al. 2015). (**A**) A branch with identified diversifying selection drawn with a thicker gray stroke. (**B**) Sequence of gene copy identified under the branch-specific diversification test is highlighted in a red rectangle.

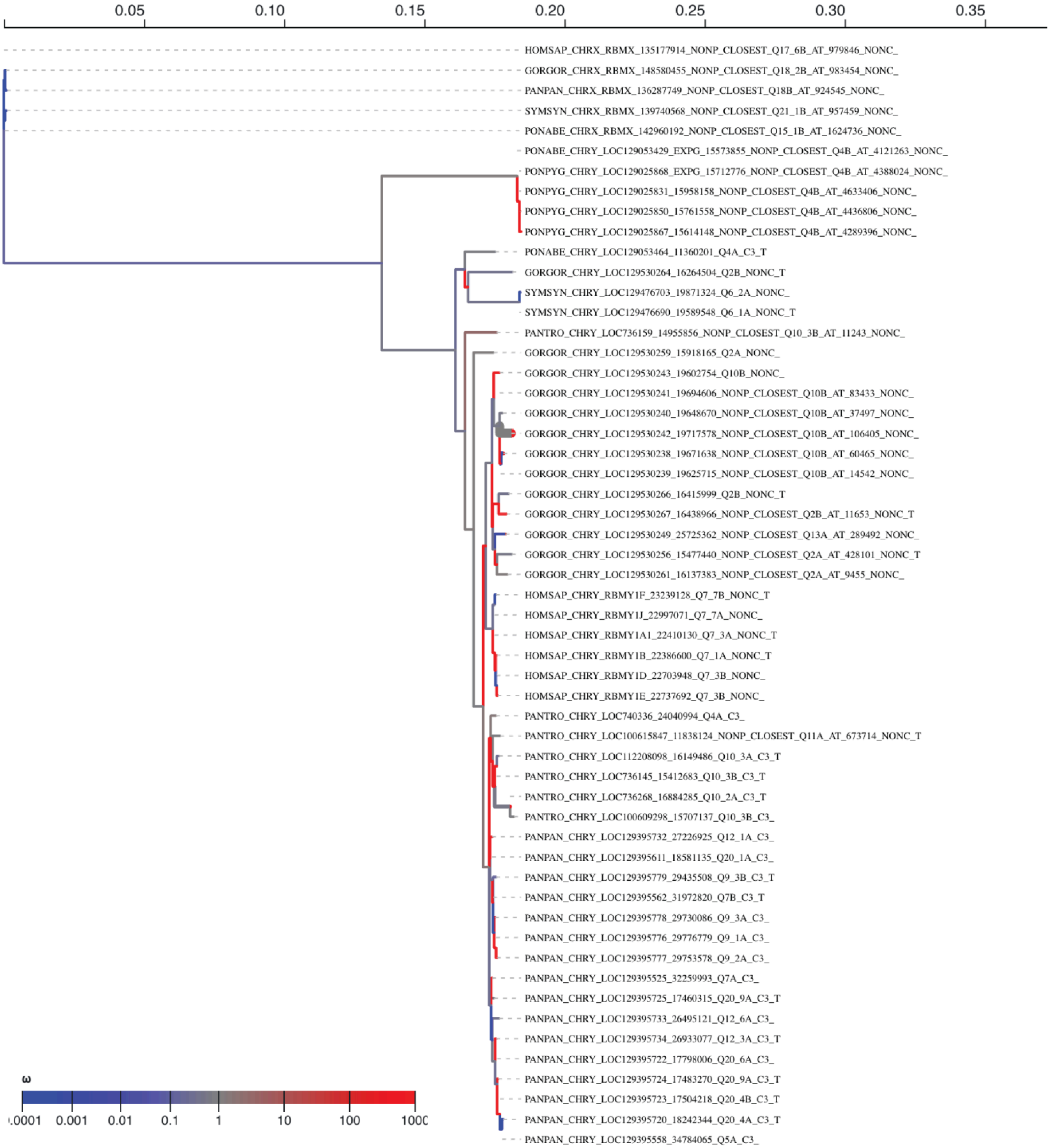

B

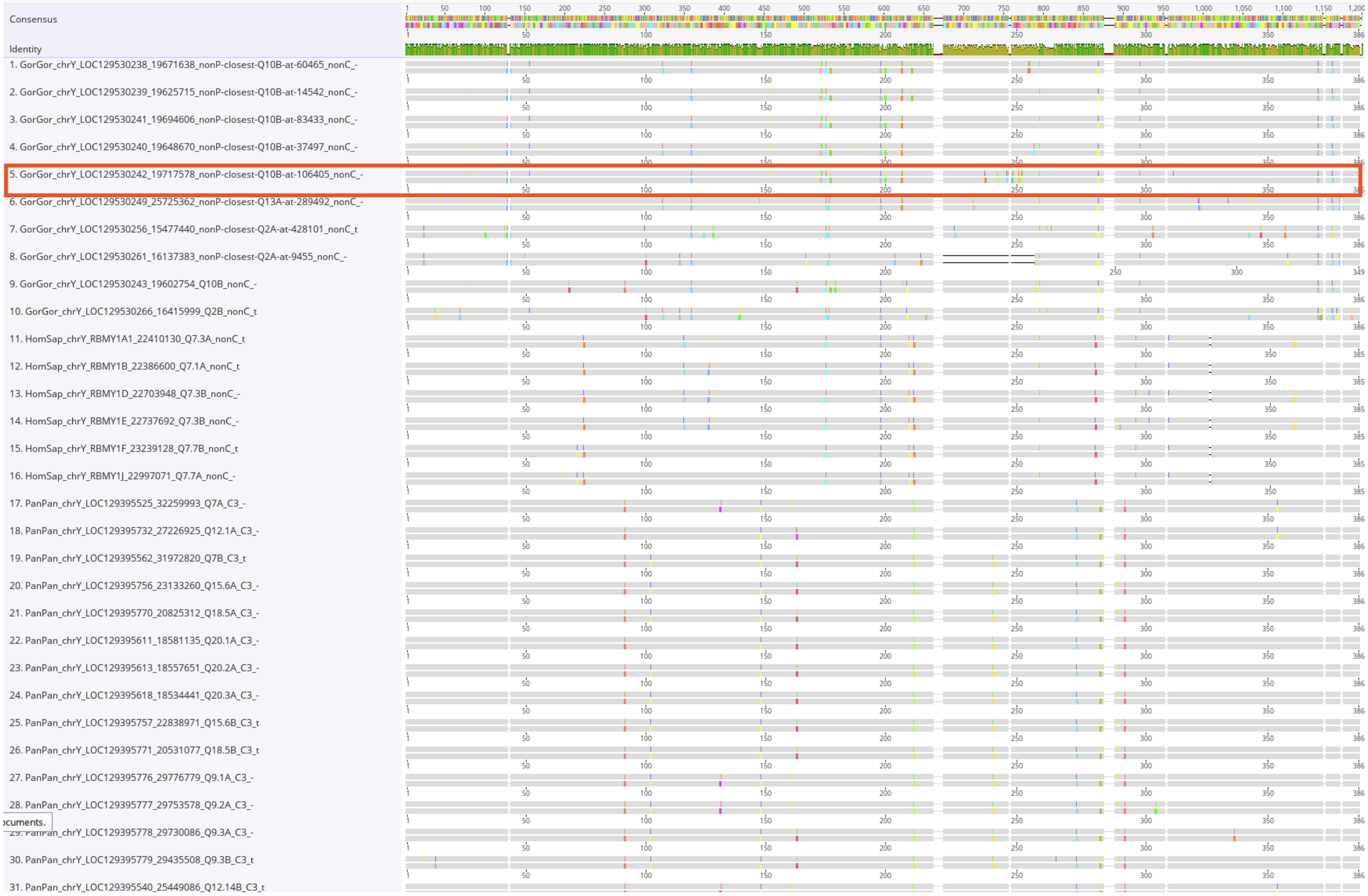

Consensus
Identity
1. GorGor_chrY_LOC129530238_19671638_nonP-closest-Q10B-at-60465_nonC_-
2. GorGor_chrY_LOC129530239_19625715_nonP-closest-Q10B-at-14542_nonC_-
3. GorGor_chrY_LOC129530241_19694606_nonP-closest-Q10B-at-83433_nonC_-
4. GorGor_chrY_LOC129530240_19648670_nonP-closest-Q10B-at-37497_nonC_-
5. GorGor_chrY_LOC129530242_19717578_nonP-closest-Q10B-at-106405_nonC_-
6. GorGor_chrY_LOC129530249_25725362_nonP-closest-Q13A-at-289492_nonC_-
7. GorGor_chrY_LOC129530256_15477440_nonP-closest-Q2A-at-428101_nonC_t
8. GorGor_chrY_LOC129530261_16137383_nonP-closest-Q2A-at-9455_nonC_-
9. GorGor_chrY_LOC129530243_19602754_Q10B_nonC_-
10. GorGor_chrY_LOC129530266_16415999_Q2B_nonC_t
11. HomSap_chrY_RBMY1A1_22410130_Q7.3A_nonC_t
12. HomSap_chrY_RBMY1B_22386600_Q7.1A_nonC_t
13. HomSap_chrY_RBMY1D_22703948_Q7.3B_nonC_-
14. HomSap_chrY_RBMY1E_22737692_Q7.3B_nonC_-
15. HomSap_chrY_RBMY1F_23239128_Q7.7B_nonC_t
16. HomSap_chrY_RBMY1J_22997071_Q7.7A_nonC_-
17. PanPan_chrY_LOC129395525_32259993_Q7A_C3_-
18. PanPan_chrY_LOC129395732_27226925_Q12.1A_C3_-
19. PanPan_chrY_LOC129395562_31972820_Q7B_C3_t
20. PanPan_chrY_LOC129395756_23133260_Q15.6A_C3_-
21. PanPan_chrY_LOC129395770_20825312_Q18.5A_C3_-
22. PanPan_chrY_LOC129395611_18581135_Q20.1A_C3_-
23. PanPan_chrY_LOC129395613_18557651_Q20.2A_C3_-
24. PanPan_chrY_LOC129395618_18534441_Q20.3A_C3_-
25. PanPan_chrY_LOC129395757_22838971_Q15.6B_C3_t
26. PanPan_chrY_LOC129395771_20531077_Q18.5B_C3_t
27. PanPan_chrY_LOC129395776_29776779_Q9.1A_C3_-
28. PanPan_chrY_LOC129395777_29753578_Q9.2A_C3_-
29. PanPan_chrY_LOC129395778_29730086_Q9.3A_C3_-
30. PanPan_chrY_LOC129395779_29435508_Q9.3B_C3_t
31. PanPan_chrY_LOC129395540_25449086_Q12.14B_C3_t

### Additional Data Files

#### Additional Data File 1. Signatures vs structural isoforms

Tables with rows representing sequence signatures and columns representing transcript structural isoforms

#### Additional Data File 2. DAZ repeats

Tables of individual *DAZ* repeats identified from sequencing reads. Tables of *DAZ* repeat composition found sequencing reads.

#### Additional Data File 3. Gene Clusters

The initial set of gene copies and homologs for all YAGs.

#### Additional Data File 4. YAGs gene_review

Table of genes reviewed after manual curation.

#### Additional Data File 5. YAG GFFs

The `GFF` files contain all YAGs, including those identified via manual curation.

#### Additional Data File 6. Sequence isoforms count tables

Count tables summarizing the frequency distributions of signatures across samples

Bhowmick, Bejon Kumar, Yoko Satta, and Naoyuki Takahata. 2007. “The Origin and Evolution of Human Ampliconic Gene Families and Ampliconic Structure.” *Genome Research* 17 (4): 441–50.

Cao, Peng-Rong, Lei Wang, Yu-Chao Jiang, Yin-Sha Yi, Fang Qu, Tao-Cheng Liu, and Yuan Lv. 2015. “De Novo Origin of VCY2 from Autosome to Y-Transposed Amplicon.” *PloS One* 10 (3): e0119651.

Delbridge, Margaret L., Guy Longepied, Danielle Depetris, Marie-Genevieve Mattei, Christine M. Disteche, Jennifer A. Marshall Graves, and Michael J. Mitchell. 2004. “TSPY, the Candidate Gonadoblastoma Gene on the Human Y Chromosome, Has a Widely Expressed Homologue on the X - Implications for Y Chromosome Evolution.” *Chromosome Research: An International Journal on the Molecular, Supramolecular and Evolutionary Aspects of Chromosome Biology* 12 (4): 345–56.

Dorus, Steve, Sandra L. Gilbert, Michele L. Forster, Robert J. Barndt, and Bruce T. Lahn. 2003. “The CDY-Related Gene Family: Coordinated Evolution in Copy Number, Expression Profile and Protein Sequence.” *Human Molecular Genetics* 12 (14): 1643–50.

Elliott, David J., Caroline Dalgliesh, Gerald Hysenaj, and Ingrid Ehrmann. 2019. “RBMX Family Proteins Connect the Fields of Nuclear RNA Processing, Disease and Sex Chromosome Biology.” *The International Journal of Biochemistry & Cell Biology* 108 (March):1–6.

Hughes, Jennifer F., Helen Skaletsky, and David C. Page. 2012. “Sequencing of Rhesus Macaque Y Chromosome Clarifies Origins and Evolution of the DAZ (Deleted in AZoospermia) Genes.” *BioEssays: News and Reviews in Molecular, Cellular and Developmental Biology* 34 (12): 1035–44.

Hughes, Jennifer F., Helen Skaletsky, Tatyana Pyntikova, Natalia Koutseva, Terje Raudsepp, Laura G. Brown, Daniel W. Bellott, Ting-Jan Cho, Shannon Dugan-Rocha, Ziad Khan, Colin Kremitzki, Catrina Fronick, Tina A. Graves-Lindsay, Lucinda Fulton, Wesley C. Warren, Richard K. Wilson, Elaine Owens, James E. Womack, William J. Murphy, Donna M. Muzny, Kim C. Worley, Bhanu P. Chowdhary, Richard A. Gibbs, and David C. Page. 2020. “Sequence Analysis in Bos Taurus Reveals Pervasiveness of X-Y Arms Races in Mammalian Lineages.” *Genome Research* 30 (12): 1716–26.

Iwase, Mineyo, Yoko Satta, Hirohisa Hirai, Yuriko Hirai, and Naoyuki Takahata. 2010. “Frequent Gene Conversion Events between the X and Y Homologous Chromosomal Regions in Primates.” *BMC Evolutionary Biology* 10 (July):225.

Jukes, Thomas H., and Charles R. Cantor. 1969. “Evolution of Protein Molecules.” In *Mammalian Protein Metabolism*, 21–132. Elsevier.

Jumper, John, Richard Evans, Alexander Pritzel, Tim Green, Michael Figurnov, Olaf Ronneberger, Kathryn Tunyasuvunakool, Russ Bates, Augustin Žídek, Anna Potapenko, Alex Bridgland, Clemens Meyer, Simon A. A. Kohl, Andrew J. Ballard, Andrew Cowie, Bernardino Romera-Paredes, Stanislav Nikolov, Rishub Jain, Jonas Adler, Trevor Back, Stig Petersen, David Reiman, Ellen Clancy, Michal Zielinski, Martin Steinegger, Michalina Pacholska, Tamas Berghammer, Sebastian Bodenstein, David Silver, Oriol Vinyals, Andrew W. Senior, Koray Kavukcuoglu, Pushmeet Kohli, and Demis Hassabis. 2021. “Highly Accurate Protein Structure Prediction with AlphaFold.” *Nature* 596 (7873): 583–89.

Katoh, Kazutaka, and Daron M. Standley. 2013. “MAFFT Multiple Sequence Alignment Software Version 7: Improvements in Performance and Usability.” *Molecular Biology and Evolution* 30 (4): 772–80.

Lahn, B. T., and D. C. Page. 1999. “Retroposition of Autosomal mRNA Yielded Testis-Specific Gene Family on Human Y Chromosome.” *Nature Genetics* 21 (4): 429–33.

Letunic, Ivica, and Peer Bork. 2024. “Interactive Tree of Life (iTOL) v6: Recent Updates to the Phylogenetic Tree Display and Annotation Tool.” *Nucleic Acids Research* 52 (W1): W78–82.

Lingenfelter, P. A., M. L. Delbridge, S. Thomas, H. E. Hoekstra, M. J. Mitchell, J. A. Graves, and C. M. Disteche. 2001. “Expression and Conservation of Processed Copies of the RBMX Gene.” *Mammalian Genome: Official Journal of the International Mammalian Genome Society* 12 (7): 538–45.

Makova, Kateryna D., Brandon D. Pickett, Robert S. Harris, Gabrielle A. Hartley, Monika Cechova, Karol Pal, Sergey Nurk, Dongahn Yoo, Qiuhui Li, Prajna Hebbar, Barbara C. McGrath, Francesca Antonacci, Margaux Aubel, Arjun Biddanda, Matthew Borchers, Erich Bornberg-Bauer, Gerard G. Bouffard, Shelise Y. Brooks, Lucia Carbone, Laura Carrel, Andrew Carroll, Pi-Chuan Chang, Chen-Shan Chin, Daniel E. Cook, Sarah J. C. Craig, Luciana de Gennaro, Mark Diekhans, Amalia Dutra, Gage H. Garcia, Patrick G. S. Grady, Richard E. Green, Diana Haddad, Pille Hallast, William T. Harvey, Glenn Hickey, David A. Hillis, Savannah J. Hoyt, Hyeonsoo Jeong, Kaivan Kamali, Sergei L. Kosakovsky Pond, Troy M. LaPolice, Charles Lee, Alexandra P. Lewis, Yong-Hwee E. Loh, Patrick Masterson, Kelly M. McGarvey, Rajiv C. McCoy, Paul Medvedev, Karen H. Miga, Katherine M. Munson, Evgenia Pak, Benedict Paten, Brendan J.

Pinto, Tamara Potapova, Arang Rhie, Joana L. Rocha, Fedor Ryabov, Oliver A. Ryder, Samuel Sacco, Kishwar Shafin, Valery A. Shepelev, Viviane Slon, Steven J. Solar, Jessica M. Storer, Peter H. Sudmant, Sweetalana, Alex Sweeten, Michael G. Tassia, Françoise Thibaud-Nissen, Mario Ventura, Melissa A. Wilson, Alice C. Young, Huiqing Zeng, Xinru Zhang, Zachary A. Szpiech, Christian D. Huber, Jennifer L. Gerton, Soojin V. Yi, Michael C. Schatz, Ivan A. Alexandrov, Sergey Koren, Rachel J. O'Neill, Evan E. Eichler, and Adam M. Phillippy. 2024. "The Complete Sequence and Comparative Analysis of Ape Sex Chromosomes." *Nature* 630 (8016): 401–11.

Saitou, N., and M. Nei. 1987. "The Neighbor-Joining Method: A New Method for Reconstructing Phylogenetic Trees." *Molecular Biology and Evolution* 4 (4): 406–25.

Saxena, R., L. G. Brown, T. Hawkins, R. K. Alagappan, H. Skaletsky, M. P. Reeve, R. Reijo, S. Rozen, M. B. Dinulos, C. M. Disteche, and D. C. Page. 1996. "The DAZ Gene Cluster on the Human Y Chromosome Arose from an Autosomal Gene That Was Transposed, Repeatedly Amplified and Pruned." *Nature Genetics* 14 (3): 292–99.

Schrödinger, LLC. 2026. *The PyMOL Molecular Graphics System, Version 3.0*. http://www.pymol.org/pymol.

Skaletsky, Helen, Tomoko Kuroda-Kawaguchi, Patrick J. Minx, Holland S. Cordum, Ladeana Hillier, Laura G. Brown, Sjoerd Repping, Tatyana Pyntikova, Johar Ali, Tamberlyn Bieri, Asif Chinwalla, Andrew Delehaunty, Kim Delehaunty, Hui Du, Ginger Fewell, Lucinda Fulton, Robert Fulton, Tina Graves, Shun-Fang Hou, Philip Latrielle, Shawn Leonard, Elaine Mardis, Rachel Maupin, John McPherson, Tracie Miner, William Nash, Christine Nguyen, Philip Ozersky, Kymberlie Pepin, Susan Rock, Tracy Rohlfing, Kelsi Scott, Brian Schultz, Cindy Strong, Aye Tin-Wollam, Shiaw-Pyng Yang, Robert H. Waterston, Richard K. Wilson, Steve Rozen, and David C. Page. 2003. "The Male-Specific Region of the Human Y Chromosome Is a Mosaic of Discrete Sequence Classes." *Nature* 423 (6942): 825–37.

Smith, Martin D., Joel O. Wertheim, Steven Weaver, Ben Murrell, Konrad Scheffler, and Sergei L. Kosakovsky Pond. 2015. "Less Is More: An Adaptive Branch-Site Random Effects Model for Efficient Detection of Episodic Diversifying Selection." *Molecular Biology and Evolution* 32 (5): 1342–53.

Sweeten, Alexander P., Michael C. Schatz, and Adam M. Phillippy. 2024. "ModDotPlot-Rapid and Interactive Visualization of Tandem Repeats." *Bioinformatics (Oxford, England)* 40 (8). https://doi.org/10.1093/bioinformatics/btae493.

Vallender, Eric J., and Bruce T. Lahn. 2004. "How Mammalian Sex Chromosomes Acquired Their Peculiar Gene Content." *BioEssays: News and Reviews in Molecular, Cellular and Developmental Biology* 26 (2): 159–69.

Xu, E. Y., F. L. Moore, and R. A. Pera. 2001. "A Gene Family Required for Human Germ Cell Development Evolved from an Ancient Meiotic Gene Conserved in Metazoans." *Proceedings of the National Academy of Sciences of the United States of America* 98 (13): 7414–19.
